# Multi-Peptide Prompting Enables In-Context Learning in Protein Language Models

**DOI:** 10.64898/2026.08.26.747074

**Authors:** Joshua Almonte, Minh Triet Vu, Andrew Ahn, Erik H. Thiede

## Abstract

We show that single-sequence PLMs can perform in-context peptide learning without gradient updates, task-specific retraining, or architectural modification. We introduce multi-peptide example prompts (MPEPs), in which demonstration peptides are concatenated with glycine spacers and used as context for scoring query peptides by their prompted probability. We evaluate this approach across multiple tasks, using both encoder-only ESM-2 models and decoder-only ProGen2 models. Across most tasks, performance improves with the number of peptide examples and with model scale, indicating that PLMs can extract shared properties from prompted examples. We further introduce a difference score that contrasts positive-example and negative-example MPEPs, reducing compositional biases in raw PLM probabilities and substantially improving classification. Perhaps surprisingly, when applied to few-shot peptide learning problems MPEP-based classification is competitive with classifiers trained on frozen ESM-2 embeddings, despite requiring no training. These results reveal an unexpected in-context inference capability in single-sequence PLMs and establish in-context learning as a potential lightweight strategy for low-data peptide classification.

## 1 Introduction

Peptides are an increasingly important therapeutic modality because they can provide the high target specificity and affinity of biologics with lower production cost and immunogenicity risk (Fosgerau & Hoffmann, 2015; Wang et al., 2022). However, peptide discovery remains fundamentally a search problem over an enormous combinatorial space. A peptide of length 12 spans more than 4 × 10^15^ possible primary sequences, making exhaustive experimental exploration impossible. Moreover, experimental characterization is costly, and for many peptide properties only limited data is available (Zhou et al., 2024). As a result, modern peptide discovery pipelines rely heavily on computational methods to prioritize candidates prior to synthesis and experimental characterization (Ansari & D. White, 2024; Goles et al., 2024; Wu et al., 2024).

Protein language models (PLMs) such as ESM-2 (Lin et al., 2023) and ProGen2 (Nijkamp et al., 2023) have emerged as a powerful technology for analyzing amino acid sequences, and hold promise as tools for peptide discovery. PLMs use neural network architectures initially developed for natural language and are pretrained on large protein sequence corpora. They learn sequence representations that capture structural and functional information useful for downstream prediction tasks (Schmirler et al., 2024; Vieira et al., 2025; Brixi et al., 2023; Chen et al., 2025). For instance, sequence-level scores derived from PLM amino-acid probabilities have been used to rank and prioritize candidate sequences (Chen et al., 2025; Kantroo et al., 2025a; Nijkamp et al., 2023; Notin et al., 2022).

The similarity between natural language and protein sequence models raises a fundamental question: can prompting strategies used to adapt pretrained LLMs in the low-data regime be used to adapt PLMs? We focus on an approach that has seen notable success for LLMs: in-context learning. Rather than updating model parameters, in-context learning conditions a pretrained language model on a small set of examples supplied directly in the prompt, allowing the model’s behavior to adapt at inference time (Brown et al., 2020; Dong et al., 2024). The examples act as demonstrations that influence subsequent predictions without any gradient updates. Subsequent work has shown that large language models use this mechanism to learn across a broad variety of tasks, including forecasting dynamical systems (Gruver et al., 2023) and performing density estimation (Liu et al., 2025). But in contrast to LLMs, most PLMs are trained on individual protein sequences and are never explicitly trained to compare multiple sequences or aggregate information across demonstrations. Some recent efforts have addressed this problem by incorporate multi-sequence conditioning through dedicated architectures or training schemes (Notin et al., 2023; Beck et al., 2025; Truong Jr & Bepler, 2023; Ma et al., 2025). We complement this work by asking whether off-the-shelf, single-sequence PLMs already support in-context inference.

We hypothesize that PLMs can use multiple peptide examples provided at inference time as contextual information and apply information extracted from those examples when evaluating a query peptide. To test this hypothesis, we developed the **Multi-Peptide Example Prompt (MPEP)**, a prompt format in which peptide demonstrations are concatenated using glycine spacers and supplied to a pretrained PLM. The model’s native token probabilities are then used to score the query, and a score threshold is applied to enable peptide classification without gradient updates, parameter modification, or task-specific retraining.

### Main Contributions

We demonstrate that pretrained, single-sequence PLMs are capable of incontext learning.

1. Multi-peptide example prompts improve peptide classification performance on multiple tasks, including secondary-structure classification and MHC-II binding prediction tasks, in both a masked and autoregressive protein language model.
2. A log-probability-difference score, which contrasts a positive-example MPEP against a negative-example MPEP, helps isolate context-dependent signal from the model’s intrinsic compositional preferences and substantially improves classification.
3. Classification performance increases both with the number of examples and model size, similar to in-context learning in natural language models.

Our results give new insight into the capabilities of protein language models and reveal a simple and practical strategy for low-data peptide classification.

## 2 Method

The core of our method is the construction of a multi-peptide example prompt (MPEP) comprised of example peptides that share a common property. Concretely, given *k* example peptides {*E*_1_, …, *E*_*k*_} we form a prompt by concatenating the sequences together, delimited by a spacer. In this work, we use a five-glycine spacer; we leave exploration and optimization of the spacer for future work.

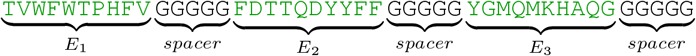

We use this prompt to evaluate query peptides by computing their conditional log-probability under the pretrained protein language model.

### 2.1 Conditional Peptide Probabilities using MPEPs

We apply our prompts to two architecturally distinct PLM families: an encoder-only masked language model (ESM-2) and a decoder-only autoregressive model (ProGen2). The prompt layout is shared; what differs between the two is how the query positions are realized and how the score is computed. We depict both scoring procedures in Figure 1.

**Figure 1:**
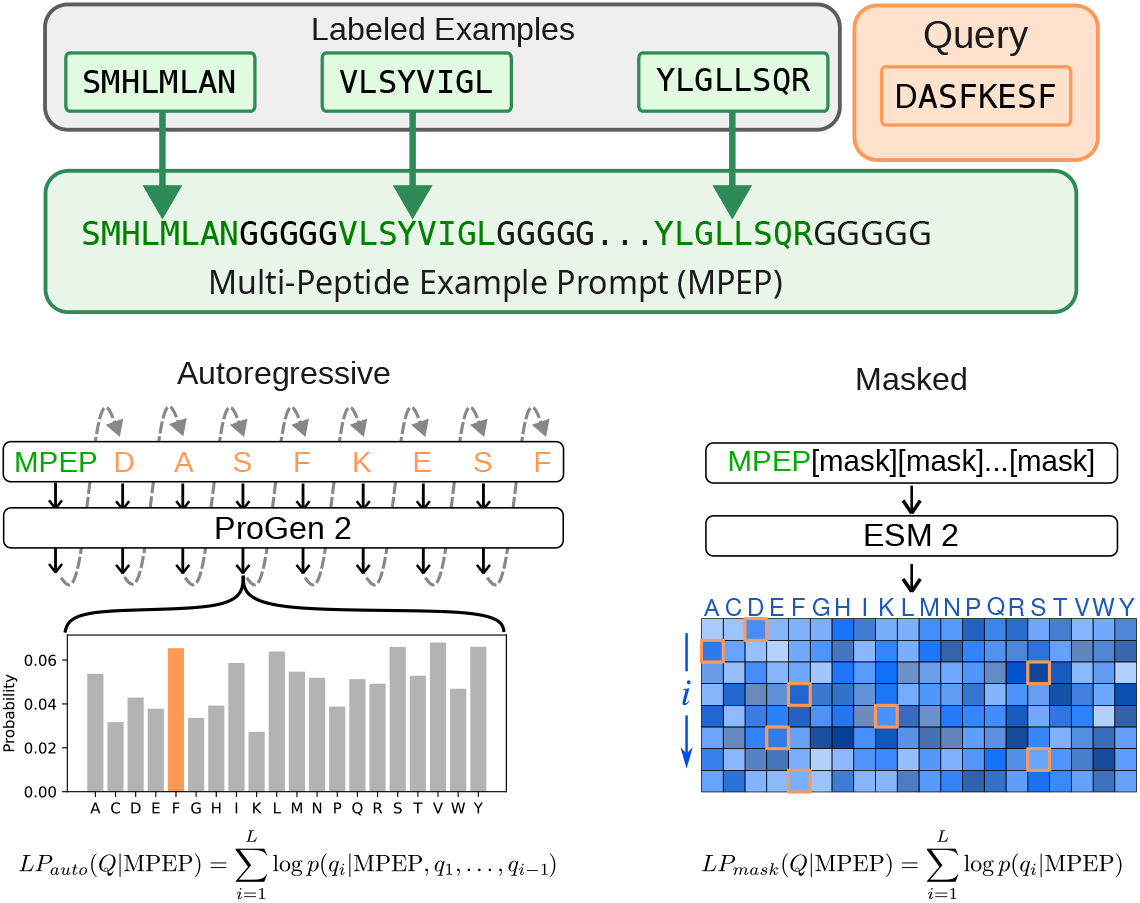
Schematic depicting how Multi-Peptide Example Prompts (MPEPs) are constructed, as well as how they are used in both autoregressive and masked language models.

#### Prompted Log-Probabilities from Masked Language Models

Masked language models predict amino acids at [MASK] positions conditioned on the rest of the sequence (Lin et al., 2023). Consequently, we append *L* = |*Q*| contiguous [MASK] tokens to the prompt.

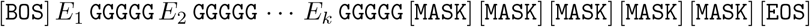

where [BOS] is the beginning of sequence token, [EOS] is the end of sequence token, *E*_*i*_ is the *i*th demonstration peptide. In this example, the length of the query peptide is five amino acids. ESM-2 is run in a single forward pass per prompt, yielding categorical distributions over the vocabulary at every mask position. We then calculate the log-probability of the query peptide *Q* by summing the log-probabilities at each masked position:

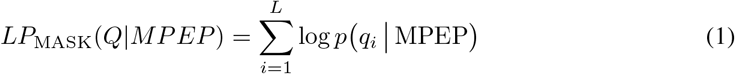

A high *LP*_MASK_ means that, having seen the context examples, the model considers the query likely. Note that this model attaches a probability to the amino acid identity at each position: effectively, our masked language model has constructed a position-specific scoring matrix (Gribskov et al., 1987).

#### Prompted Log-probabilities from Autoregressive PLMs

For decoder-only autoregressive PLMs such as ProGen2(Nijkamp et al., 2023) there are no mask tokens. Instead, the model factorizes the probability of a sequence as a product of left-to-right conditional probabilities. To enable MPEP conditioning in this setting, we construct an MPEP using the same demonstration-plus-spacer layout.

For a query peptide *Q* = (*q*_0_, *q*_1_, …, *q*_*L−*1_) of length *L*, we construct *L* prompts

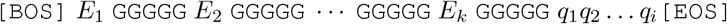

for 1 ≤ *i* ≤ *L*. We then calculate the peptide’s log-probability by summing over the sequence

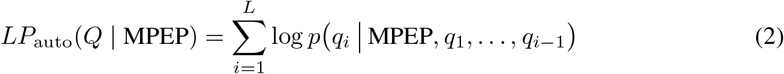

In theory, this can be evaluated in a single forward pass of the model due to attention causality. In contrast to log-probabilities evaluated on masked language models this allows for conditional dependence between amino acids.

### 2.2 Peptide Classification using MPEP-derived Scores

To convert log-probabilities into classifications, we employ two scoring variants: a *positive-only score* that uses only demonstrations of the target class, and a *difference score*. For our positive-only score, we set our score to be the prompted log-probability

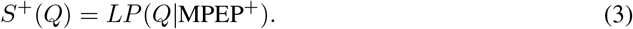

where MPEP^+^ is an MPEP made of only positive examples. While straightforward, the positive-only score conflates three distinct signals: the model’s unconditioned preference for certain residue compositions, the presence of a linker sequence (polyglycine for most of our experiments), and the contribution of the in-context examples. To help isolate the latter, we construct an additional prompt MPEP^*−*^ with the same number of examples and linkers as MPEP^+^ but using negative examples. We then score peptides by the difference in the two log-probabilities,

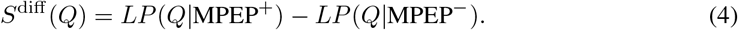

Code implementing these methods and links to datasets are available at https://github.com/thiedelab/peptide_icl.

## 3 Experiments

We evaluated in-context learning capability on three experiments: a synthetic pattern-completion task that probes whether a PLM can use repeated structure within an MPEP despite biological implausibility, a MHC-II peptide binder classification task demonstrating the phenomenon on a biologically relevant dataset, and a random sequence task designed to detect the presence of induction heads. Additional experiments demonstrating in-context learning on other biochemically relevant datasets are given in the supplement and described at the end of the section.

### 3.1 Pattern completion

We first test the simplest prerequisite of in-context learning: can a PLM exploit a repeated sequence pattern within its input to learn out-of-distribution sequences? To demonstrate this task, we employ a simplified prompt, consisting of two homogeneous amino-acids, *B*_1_ and *B*_2_, where |*B*_1_|, |*B*_2_| ∈ [1, 10], repeated *r* ∈ {1, …, 5} times:

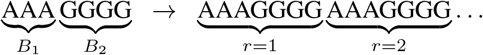

For ESM-2, we then append this prompt with |*B*_1_| mask tokens; for ProGen2, we test for the next sequence autoregressively. We score each (|*B*_1_ |, |*B*_2_ |, *r*) cell by averaging top-1 per-residue accuracy over all ordered amino-acid pairs (*a, b*) from the 20-letter alphabet. For ESM-2 we use the masked-language-model head; for ProGen2 we use argmax decoding under the autoregressive distribution. This experiment closely resembles prior work on how repeated sequences in protein sequences can affect PLM output (Kantroo et al., 2025b). While this work focused on the dangers of in-context learning distorting model fitness predictions, we focus on assessing whether in-context learning can lead to productive classification.

We summarize each model’s behavior by averaging over block sizes and plotting accuracy versus *r* and versus model size in Figure 2A. At *r* = 1, every model performs near the chance level of 1*/*20 = 5%, indicating that a single preceding block carries negligible signal. As *r* grows, accuracy increases monotonically for all models. We see the same phenomena for the decoder-only autoregressive model ProGen2-xl, suggesting the phenomenon is not unique to ESM2. Moreover, figure 2B shows that larger models convert repeats into larger accuracy gains. This scaling with both model size and example count is characteristic of in-context learning (Brown et al., 2020). Moreover, it cannot be attributed to memorized completions from the training data because the synthetic blocks are not realistic protein sequences. As an example, we show predicted amino acid probabilities conditioned on alternating tryptophan W and tyrosine Y repeats in Figure 2C Both are uncommon amino acids whose bulky, hydrophobic character makes such a repeat unlikely to occur in a real biological sequence. Despite this, probability mass shifts onto the residues that continue the W/Y pattern, so the model completes the pattern rather than reverting to a more biologically plausible sequence. We believe this reflects the model’s ability to learn from the in-context repeats.

**Figure 2:**
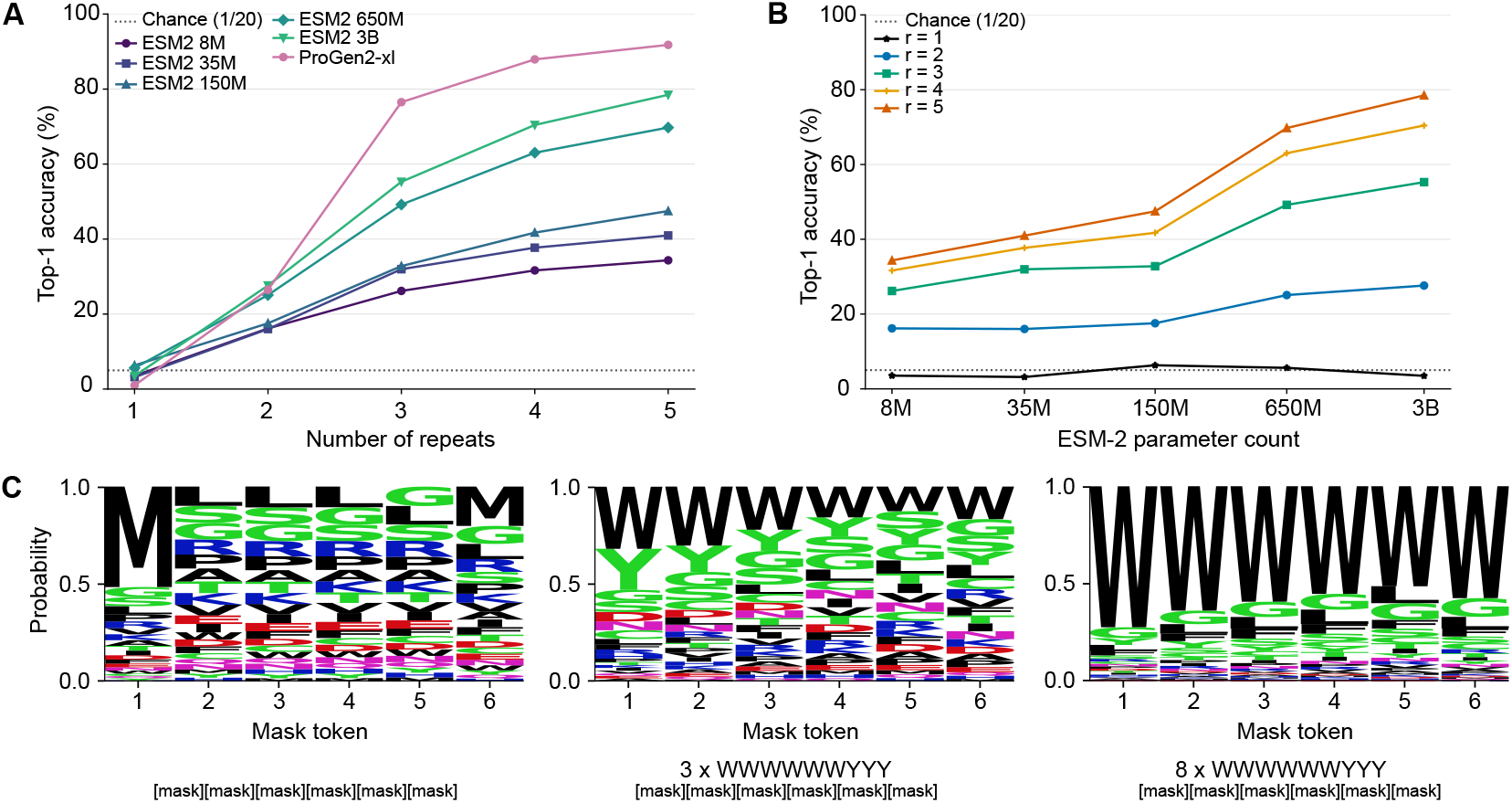
(**A**) Per-residue top-1 accuracy on the synthetic pattern-completion task as a function of the number of preceding repeats *r*. All models begin near chance (1*/*20 = 5%) at *r* = 1 and improve monotonically as more repeats are provided. (**B**) Per-residue top-1 accuracy for the same task as a function of ESM-2 model parameter count. Larger models benefit more from additional prompted examples. (**C**) Sequence logos showing how amino acids probabilities change with an increased repetition of tryptophan / tyrosine blocks.

### 3.2 Secondary-structure classification

We next ask asks whether MPEP can recover a more complex property: the propensity of a peptide to adopt either an *α*-helical fold or a *β*-hairpin fold. The *α*-helix and *β*-hairpin datasets were derived from (Tsai et al., 2022) and (DuPai et al., 2021), respectively. We restricted both datasets to peptides of length 12 and reduced sequence redundancy at a 50% identity threshold. After random train/test splitting, the *α*-helix dataset contains 3,461 training and 865 test sequences, and the *β*-hairpin dataset contains 2,366 training and 591 test sequences. For each value of *k* ∈ {0, 1, 2, 4, 8, 16, 32}, we draw 10 independent random subsets of size *k* from the training set, build positive (and, for the difference score, negative) MPEP prompts, and report mean ROC-AUC ± standard deviation across the draws. Figure 3 reports ROC-AUC ± as a function of the number of in-context examples for both classes and both scoring variants on ESM-2. We focus on ESM-2 for simplicity in this Subsection; in Appendix Fig. 7 we show that ProGen2-xlarge exhibits similar behavior.

**Figure 3:**
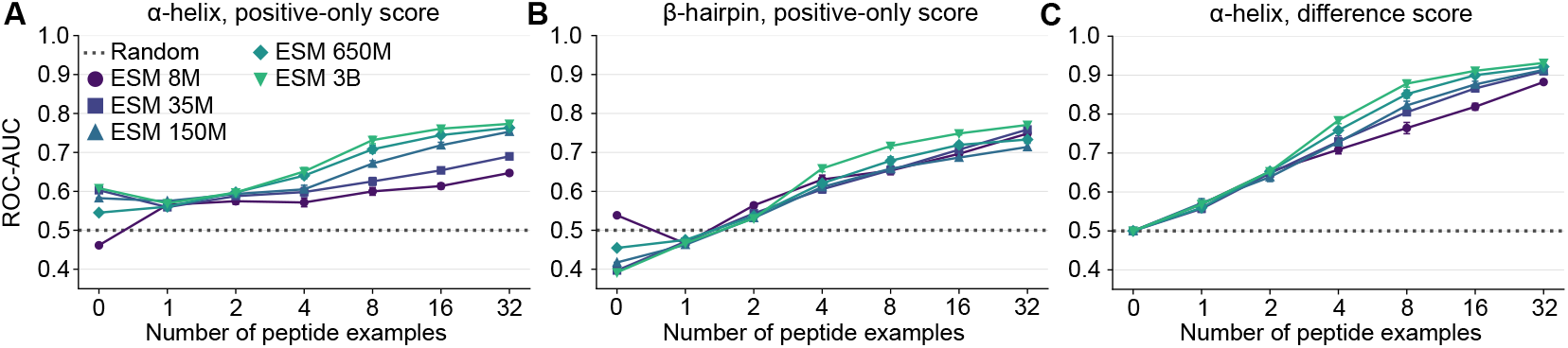
Performance of MPEP-based secondary structure classification as a function of in-context example count *k* using the ESM-2 models. Error bars are standard deviations over 10 random draws.

For *α*-helix classification, all ESM-2 models achieve above-chance ROC-AUC even at *k* = 0 when scored with the positive-only score (ROC-AUC ≈0.55–0.61, with larger models doing better). Overall performance on *α*-helix classification, (**A**), is better than *β*-hairpin classification, (**B**), using the positive-only scoring method. This indicates that *α*-helical peptides have a distinctive amino-acid composition that ESM-2’s marginal distribution already captures. The difference score (**C**; plot is the same for *α*-helix and *β*-sheet classification by symmetry) removes this zero-shot bias, bringing all models to ROC-AUC = 0.5 at *k* = 0. For all experiments performance increases monotonically with example count, with larger models retaining a consistent advantage.

### 3.3 MHC-II binder classification

To evaluate the performance of in-context learning on a nontrivial task of biomedical relevance, we attempt to predict peptide binding to the Major Histocompatibility Complex Class II (MHC-II) protein. MHC-II binding depends jointly on backbone geometry and side-chain contacts within the binding groove. To construct our dataset, we used the similarity reduced peptide binder dataset for the HLA-DR1*0101 allele from (Wang et al., 2010), ensuring minimal sequence homology between examples. We selected all peptides of length 15 and sorted them by ascending IC_50_ values. We labeled the top 250 as binders, the bottom 250 as non-binders and discarding the rest. The same evaluation protocol as the secondary-structure task was used: 10 independent draws of *k* training examples per condition for ESM-2, and 5 independent iterations for ProGen2 (following the more expensive autoregressive scoring procedure).

Figure 4 reports ROC-AUC for MHC-II binder classification under both scoring approaches for both ESM-2 and ProGen2-xl. We again see that performance improves with both the number of examples and model size. Intriguingly, we achieve effective classification despite providing no explicit information about the substrate protein. In fact, a prompt consisting of the HLA-DR1*0101 *α* and *β* chains, a five glycine spacer, and 15 mask tokens achieves an ROC-AUC of only 0.52 for MPEP-ESM2: providing the PLM with the binding partner does *not* lead to effective classification.

**Figure 4:**
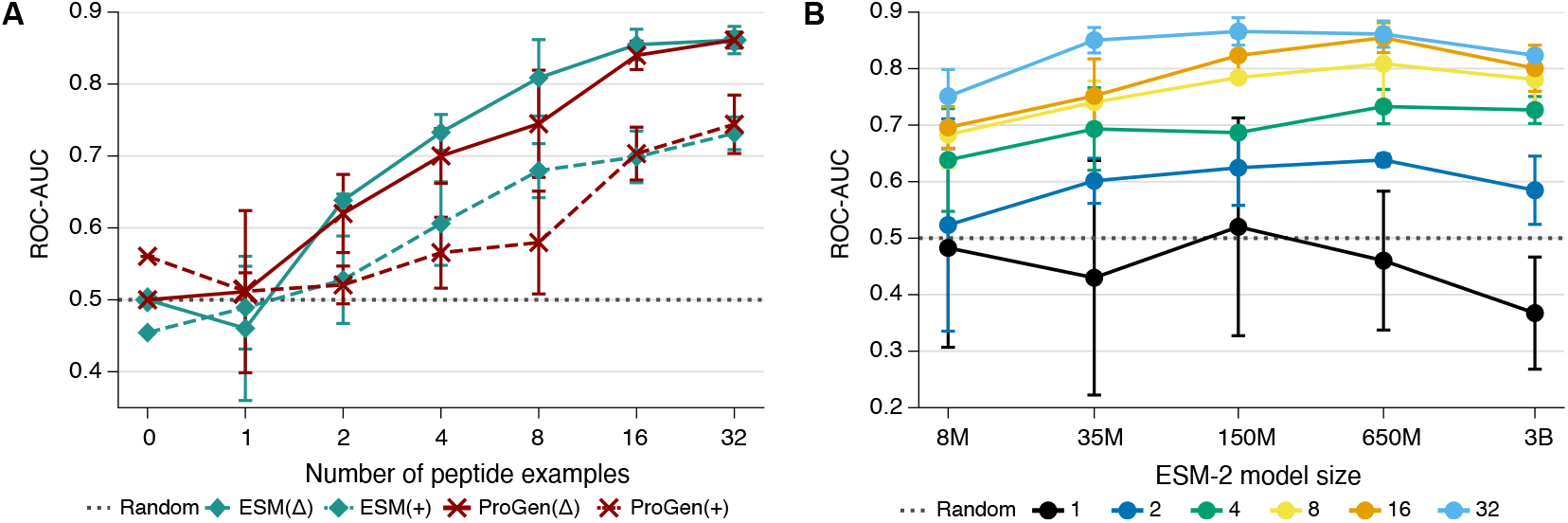
(**A**) MHC-II binder classification task with the ESM 2 650M model in green and ProGen2-xl model in red. The difference score, denoted by Δ, performs better than the positive score, denoted by +, for MPEPs with ≥ 2 peptide binder examples. (**B**) Performance on the MHC-II binder classification task as a function of ESM 2 model size. Lines give ROC-AUC for MPEPs built with either 1, 2, 4, 8, 16, or 32 peptide examples using the difference scoring method.

As with secondary structure prediction, we see that positive-only score achieves non-random performance with zero examples. We investigate this further in Table 2 of the supplement, where we show that 0-shot ROC-AUC correlates directly with how well the model’s unprompted amino acids probabilities matches the frequency of amino acids in the binder and nonbinder datasets. The difference score reduces this bias: all models have an ROC-AUC of one half with zero examples, by construction. Moreover, as we increase example count we see that the difference score substantially outperforms positive-only scoring: with 32 examples ESM-2 650M and 150M reach ROC-AUC ≈0.86, compared to ≈0.73 under the positive-only score. Despite autoregressive scoring allowing for conditional dependence between amino acids and therefore being theoretically powerful, we do not see better performance from ProGen. Further benchmarks assessing the effect of example selection and ordering are given in the Supplementary Information.

#### Benchmark against competing classifiers

To gauge the relative strength of in-context learning for PLMs, we compare the performance of ESM using MPEP with two families of baselines. First, we compare with light-weight baselines with comparable evaluation time. These include position-specific scoring matrices (PSSM) and regression from one-hot encodings. Second, we compare with classifiers trained on mean-pooled sequence embeddings from ESM. While these require more compute to evaluate, they present a stronger performance baseline. Hyperparameters were tuned using an additional validation dataset of 16 positive and 16 negative examples.

Results comparing the performance for all models are presented in Table 1. MPEP-based classification outperforms all models with comparable runtime, specifically PSSM and one-hot encoding-based models. Outperforming PSSM suggests that the model is not simply counting amino acid frequencies in the examples to perform inference. While embedding-based classifiers outperform in-context learning, they typically do so by only a small margin. MPEP-based screens are often close to, or within, the confidence intervals for classifiers trained on ESM embeddings. Moreover, embedding-based classifiers are substantially more expensive. MPEP-based screens require no explicit training steps and evaluating (1) only requires a single forward pass of ESM, as opposed to embedding-based classifiers that require one forward pass per test sequence. We graph this in Figure 5 where we show that MPEP-based screens can be three orders of magnitude faster than embedding-based classifiers.

**Table 1:** Comparison of MPEP-ESM2, a position-specific scoring matrix (PSSM), and embedding-based classifiers on the MHC-II HLA-DR1*0101 binary classification dataset. We report mean ROC-AUC ± sample standard deviation across 10 data splits. Bold indicates the highest mean in each column across all methods.

| Method / model | $k$ training examples in each class | | | | | |
| --- | --- | --- | --- | --- | --- | --- |
|  | 1 | 2 | 4 | 8 | 16 | 32 |
| **(A) MPEP** |  |  |  |  |  |  |
| MPEP-ESM-2 8M | 0.64 ± 0.14 | 0.73 ± 0.06 | 0.70 ± 0.08 | 0.77 ± 0.04 | 0.79 ± 0.03 | 0.84 ± 0.03 |
| MPEP-ESM-2 35M | **0.67 ± 0.13** | 0.73 ± 0.07 | 0.72 ± 0.06 | 0.75 ± 0.06 | 0.80 ± 0.05 | 0.86 ± 0.02 |
| MPEP-ESM-2 150M | 0.65 ± 0.14 | 0.74 ± 0.06 | 0.73 ± 0.05 | 0.79 ± 0.05 | 0.83 ± 0.04 | 0.86 ± 0.02 |
| MPEP-ESM-2 650M | 0.65 ± 0.13 | 0.73 ± 0.05 | 0.74 ± 0.05 | 0.78 ± 0.03 | 0.84 ± 0.04 | 0.86 ± 0.02 |
| **(B) PSSM and One-hot** |  |  |  |  |  |  |
| PSSM: $S^+$ | 0.57 ± 0.05 | 0.56 ± 0.05 | 0.60 ± 0.06 | 0.65 ± 0.03 | 0.70 ± 0.04 | 0.75 ± 0.03 |
| PSSM: $S^{\text{diff}}$ | 0.56 ± 0.05 | 0.56 ± 0.05 | 0.60 ± 0.05 | 0.66 ± 0.04 | 0.71 ± 0.05 | 0.77 ± 0.02 |
| One-hot: Logistic regression | 0.56 ± 0.05 | 0.56 ± 0.04 | 0.60 ± 0.05 | 0.65 ± 0.04 | 0.70 ± 0.05 | 0.74 ± 0.03 |
| One-hot: Nearest neighbor | 0.54 ± 0.03 | 0.53 ± 0.04 | 0.55 ± 0.03 | 0.56 ± 0.03 | 0.59 ± 0.04 | 0.60 ± 0.03 |
| **(C) ESM2 embedding-based classifiers** |  |  |  |  |  |  |
| ESM-2 8M: K-NN | 0.53 ± 0.06 | 0.69 ± 0.08 | 0.71 ± 0.04 | 0.76 ± 0.03 | 0.81 ± 0.03 | 0.82 ± 0.03 |
| ESM-2 8M: Logistic regression | 0.55 ± 0.09 | 0.71 ± 0.10 | 0.71 ± 0.08 | 0.80 ± 0.05 | 0.84 ± 0.02 | 0.86 ± 0.01 |
| ESM-2 8M: 3-layer MLP | 0.58 ± 0.05 | 0.73 ± 0.04 | 0.71 ± 0.07 | 0.78 ± 0.05 | 0.84 ± 0.01 | 0.85 ± 0.02 |
| ESM-2 35M: K-NN | 0.51 ± 0.08 | 0.72 ± 0.06 | 0.73 ± 0.08 | 0.78 ± 0.03 | 0.81 ± 0.03 | 0.83 ± 0.02 |
| ESM-2 35M: Logistic regression | 0.55 ± 0.15 | 0.74 ± 0.05 | 0.73 ± 0.11 | 0.80 ± 0.03 | 0.84 ± 0.03 | 0.85 ± 0.01 |
| ESM-2 35M: 3-layer MLP | 0.57 ± 0.13 | 0.73 ± 0.08 | **0.75 ± 0.06** | 0.79 ± 0.04 | 0.83 ± 0.01 | 0.85 ± 0.02 |
| ESM-2 150M: K-NN | 0.54 ± 0.08 | 0.75 ± 0.04 | 0.74 ± 0.06 | 0.81 ± 0.03 | 0.82 ± 0.03 | 0.83 ± 0.03 |
| ESM-2 150M: Logistic regression | 0.55 ± 0.13 | **0.77 ± 0.05** | **0.75 ± 0.08** | **0.82 ± 0.04** | 0.85 ± 0.02 | 0.87 ± 0.01 |
| ESM-2 150M: 3-layer MLP | 0.55 ± 0.12 | 0.75 ± 0.07 | **0.75 ± 0.08** | **0.82 ± 0.03** | 0.84 ± 0.02 | 0.86 ± 0.01 |
| ESM-2 650M: K-NN | 0.54 ± 0.09 | 0.72 ± 0.11 | 0.73 ± 0.08 | 0.81 ± 0.02 | 0.84 ± 0.02 | 0.85 ± 0.02 |
| ESM-2 650M: Logistic regression | 0.57 ± 0.12 | 0.76 ± 0.11 | 0.74 ± 0.12 | **0.82 ± 0.03** | **0.87 ± 0.02** | **0.88 ± 0.01** |
| ESM-2 650M: 3-layer MLP | 0.59 ± 0.11 | 0.74 ± 0.13 | 0.74 ± 0.10 | 0.81 ± 0.05 | 0.86 ± 0.03 | 0.87 ± 0.01 |

**Figure 5:**
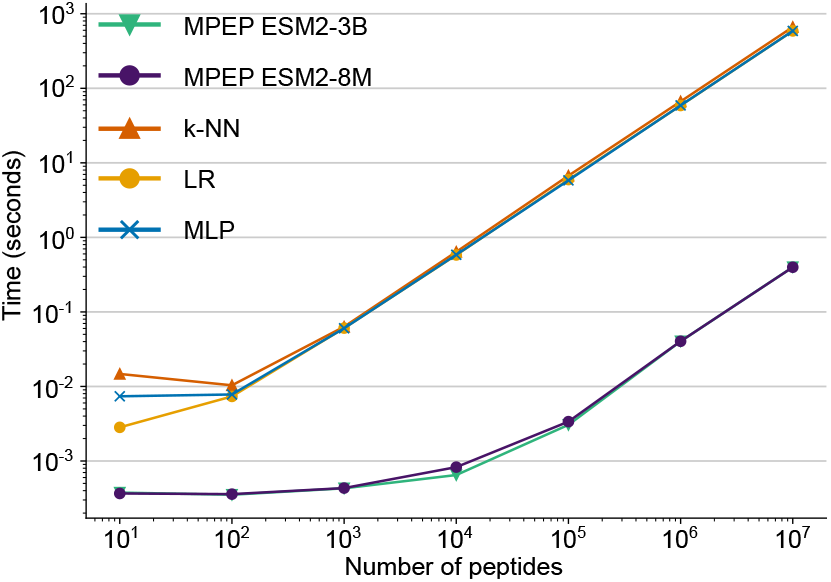
Peptide screening time comparison between MPEP and trained embedding baselines.

### 3.4 Induction Heads

Olsson et al. (2022) proposed that specific circuits known as “induction heads” are critical for incontext learning in natural language models. Here, we test for the presence of induction heads in autoregressive PLMs. For ease of presentation we only report results for ProGen2-xlarge; results for other models are given in the Supplement. Experimental designs, heuristics and scoring mechanisms are constructed as described in (Olsson et al., 2022).

A head qualifies as an induction if it exhibits *prefix matching* and *copying* capabilities. To measure these heuristics across all ProGen2 model sizes, we mimiced the prompt design by sampling 25 random tokens from the 20-aa vocabulary, excluding the most and the least popular ones. A complete sequence contains the 25-mer repeated 4 times, then prepended by “1” (BOS). We classify any head whose prefix matching and copying score exceeds 0.3 and 0.2, respectively, as induction. With these separation boundaries, we found 3-5 induction heads in each of the ProGen2 models. Scores for all heads for ProGen-xlarge are shown in Fig. 6A.

**Figure 6:**
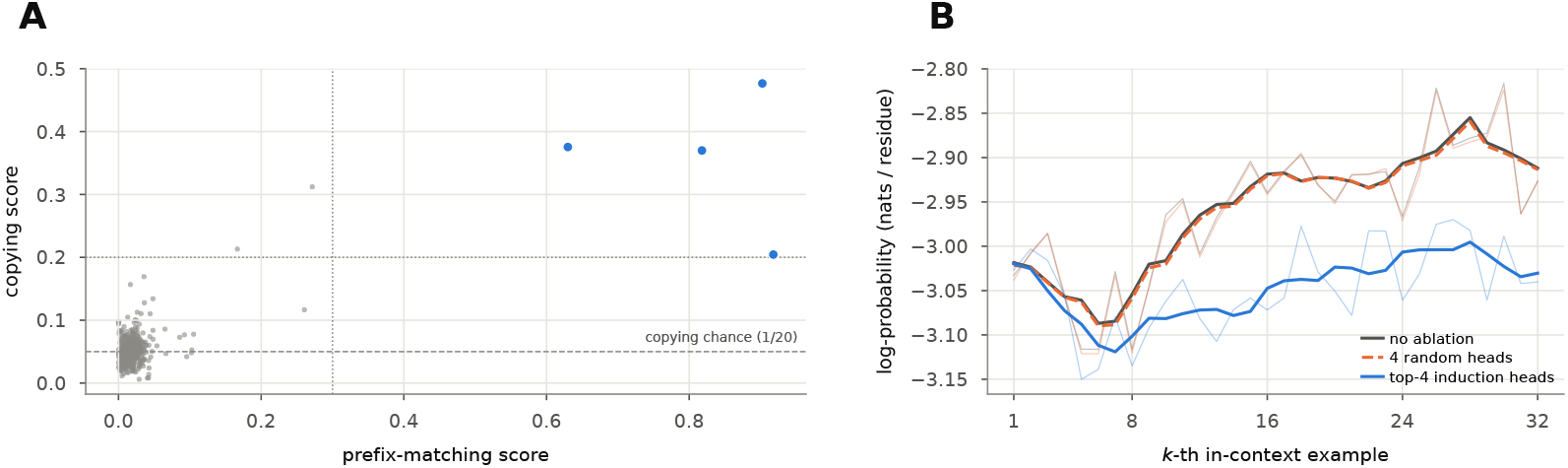
(A) Scatter plot of all heads by prefix matching and copying scores for ProGen-xlarge. (B) Test showing loss of in-context learning ability (per residue) upon ablation of the identified induction heads, averaged over 10 runs.

**Figure 7:**
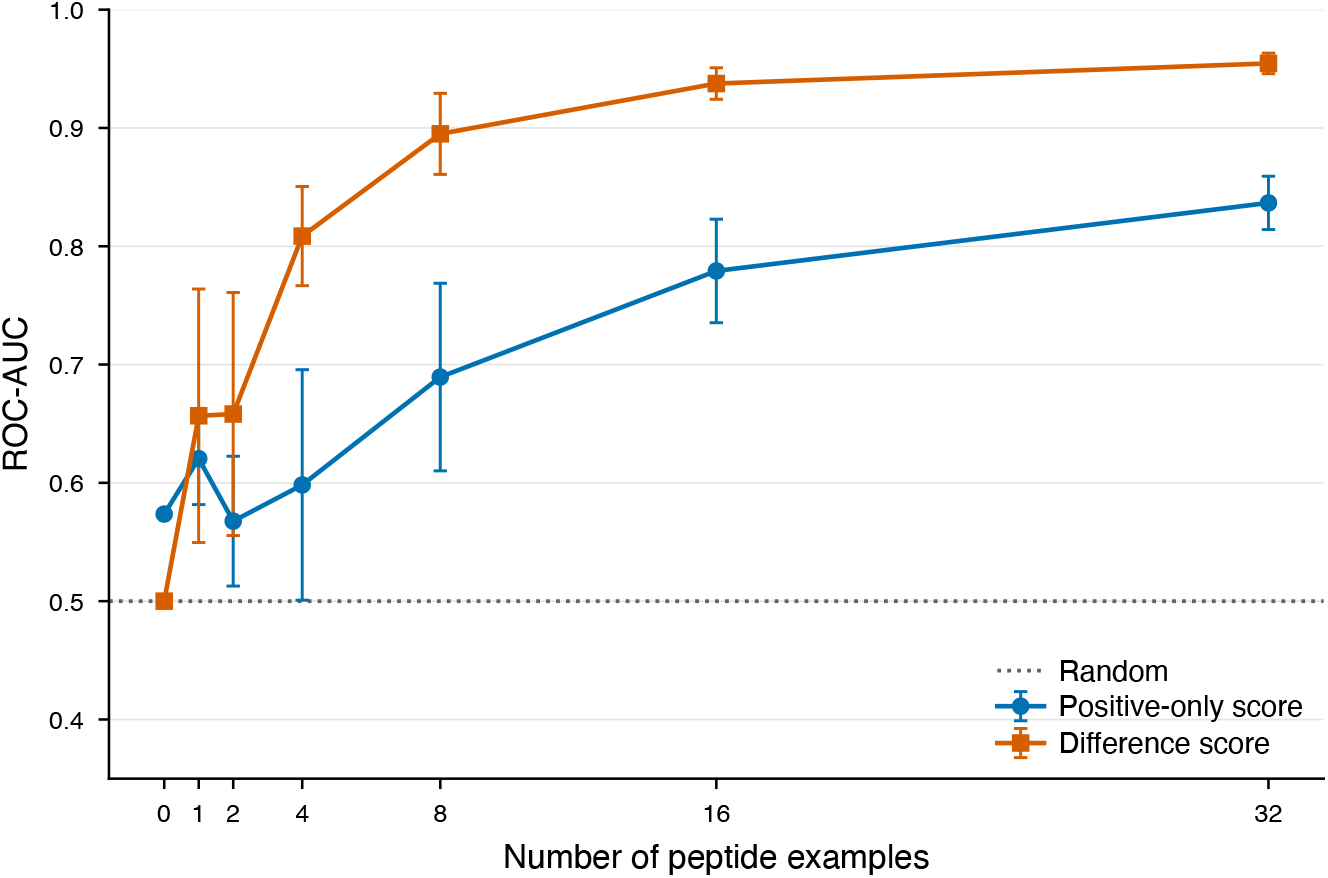
ROC-AUC for secondary-structure classification with ProGen2-xlarge as a function of in-context example count *k*. The qualitative pattern matches ESM-2: classification improves monotonically with *k*, and the difference score (orange squares) removes the zero-shot bias visible in the positive-only score (blue circles).

**Figure 8:**
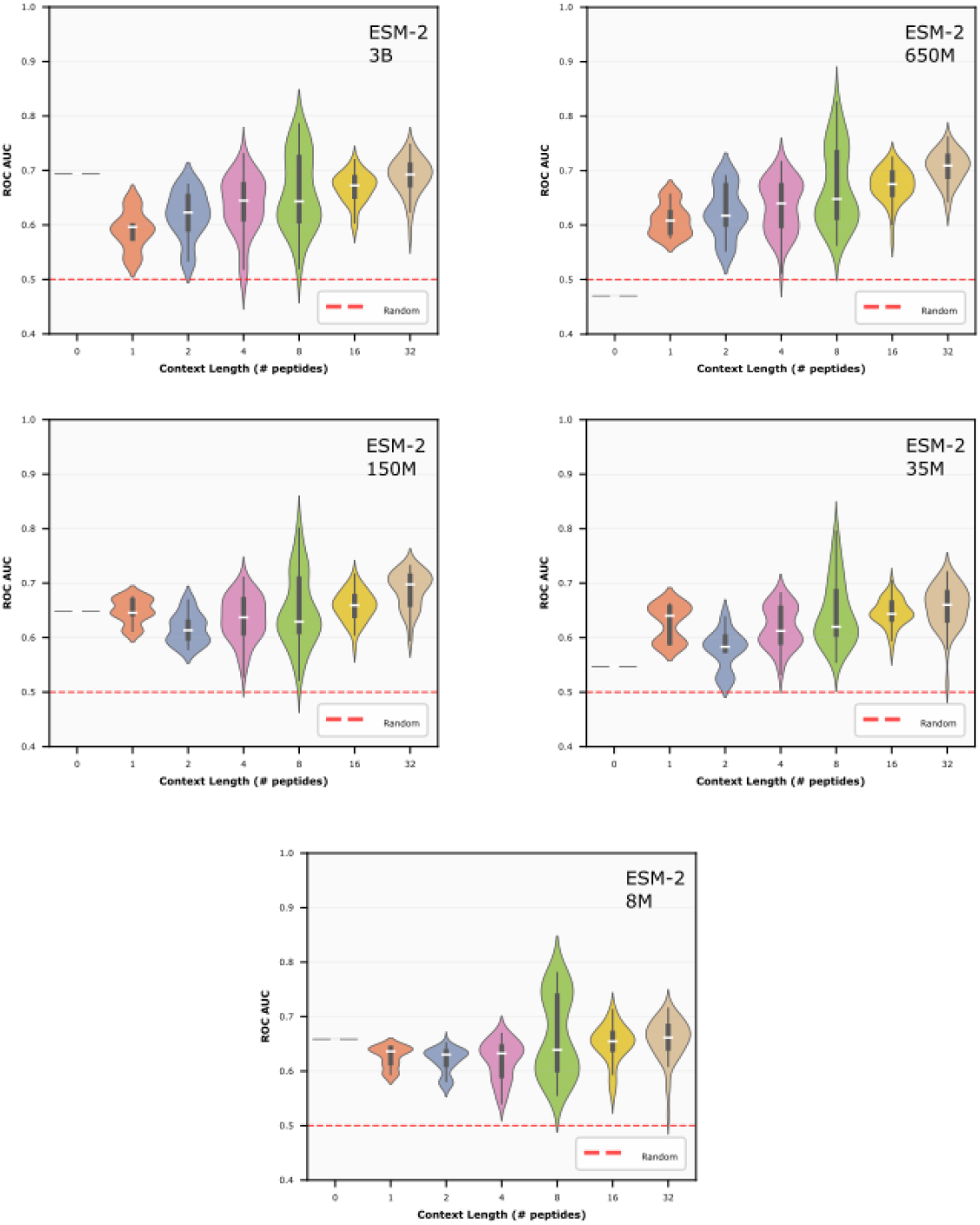
Distribution of ROC-AUC over different example selections and orderings for MHC-II binder classification, for each ESM-2 model size. For each *k*, we randomly drew 10 example subsets and computed all possible orderings (capped at 500 for *k* ≥ 8). Violins show the full distribution; white dots show the median. At low *k*, distributions are wide and some draws fall near or below chance; as *k* increases, distributions narrow and the median rises.

**Figure 9:**
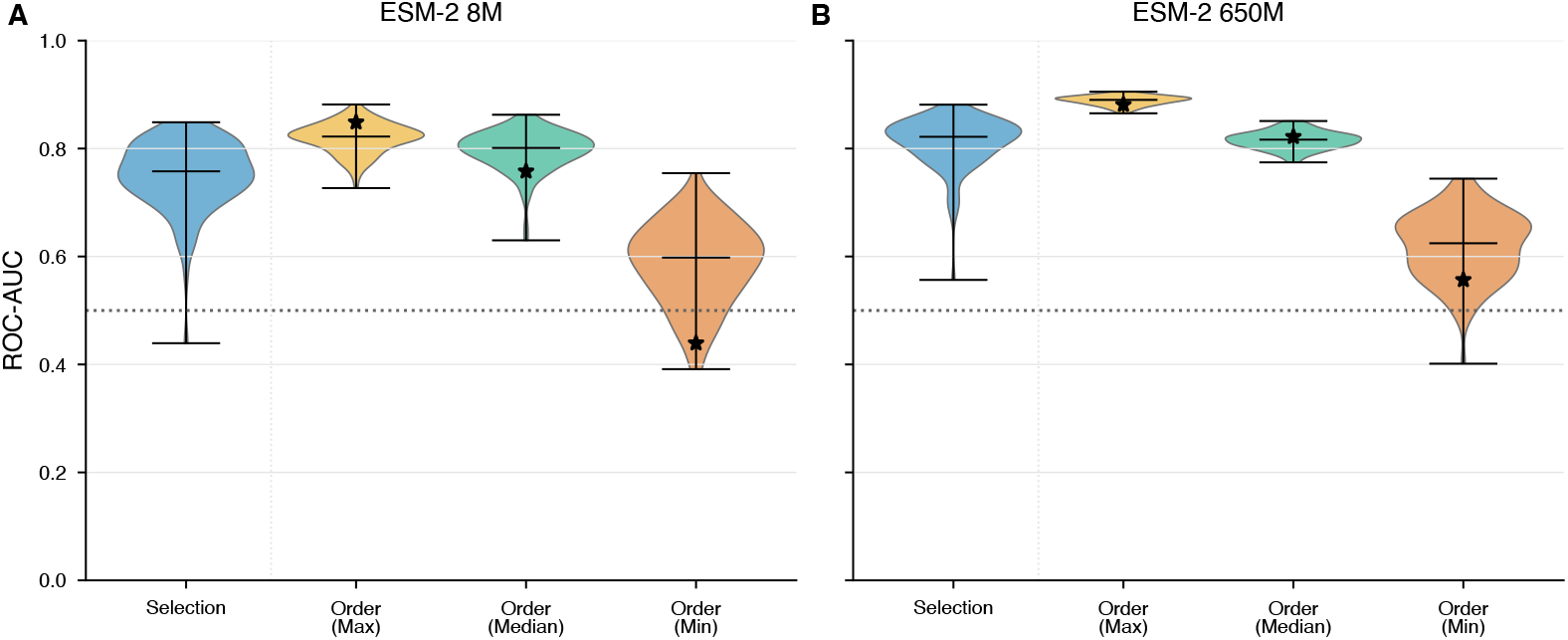
Violin plots comparing ROC-AUC across example-selection and example-ordering strategies on the MHC-II binder prediction task, for (**A**) ESM-2 8M and (**B**) ESM-2 650M. The “Selection” distribution shows ROC-AUC across 100 random selections of 16 positive and 16 negative examples from the MHC-II HLA-DRB1*0101 dataset. The “Order (Max)”, “Order (Median)”, and “Order (Min)” distributions each show 100 random shuffles of the examples selected from the top-, median-, and worst-performing selection, respectively. Black stars indicate the ROC-AUC of the specific example set used to generate each ordering distribution. Dotted line indicates chance ROC-AUC.

To test the importance of induction heads for in-context learning in PLMs, we performed an ablation test where we remove the top-*m* induction heads, and compare with the performance of the model with *m* random heads removed. We then test whether the log-probability of an MHC-II binder increases with an increasing the number of other MHC-II binders. As shown in Figure 6B, removing induction heads nearly eliminates the gain in log-probability with increasing example count, suggesting that removing the induction heads limits the model’s capability for in-context learning.

### 3.5 Additional experiments and ablations

We provide additional experiments and ablations in the supplementary information. To quantify how sensitive MPEP-based classification is to the particular examples chosen and to their order, we resample and permute examples from the MHC-II task in Section B of the Supplement. We also apply our approach to the PepBenchmark dataset (Zhang et al., 2026). We show generalizability over many tasks. We also perform several ablation studies. Notably, we show that the difference score benefits more from the amino acid positional information than the positive score.

## 4 Discussion

Our results provide strong evidence that single-sequence protein language models (PLMs) are capable of in-context peptide learning. Without task-specific training, parameter updates, or architectural modifications, PLMs can classify peptides using only information provided in the prompt. While performance does not increase strictly with size (ESM-2-3B is generally outperformed by the smaller 150M and 650M models), there is a general trend showing that performance increased performance model size. Performance also increases with the number of examples used. This is consistent with the scaling behavior observed for incontext learning in natural language models (Brown et al., 2020). Moreover, we observe this for both encoder-only (ESM-2) and decoder-only (ProGen2) architectures, suggesting that this capability is innate to PLMs, rather than the result of a specific architecture or training procedure.

On biologically relevant peptide classification tasks, MPEP-based classification is competitive with standard few-shot classifiers trained on frozen PLM embeddings, while requiring no explicit training and being considerably faster to run. Key to this strong performance is the use of a difference prompt, which helps correct for the model’s intrinsic preference for particular sequence compositions. Positive-only prompts exhibit a zero-shot preference for one class if the model believes its sequences are intrinsically more likely. The difference score addresses this with a matched negative prompt that provides a simple, model-agnostic correction. This gives a stable zero-shot baseline and improves performance at larger values of *k*. We recommend the difference score as the default scoring strategy, although positive-only prompts may be useful when negative examples are unavailable.

We believe our work gives new insight into how both in-context learning and PLMs function. Our work shows that in-context learning is not specific to NLP-based LLMs. Rather, large pretrained transformers are capable of in-context learning when trained without language data, or even data with a direct concept of examples or multiple sequences. This dissociates the phenomenon of incontext learning from text and language data, instead suggesting that in-context learning arises from model architecture and pretraining objectives, independently of the data.

Prior work has proposed that PLMs implicitly learn physical interactions (Vig et al., 2020; Rao et al., 2020) or evolutionary relationships between amino acids (Zhang et al., 2024). While our results do not contradict these claims, they suggest that PLMs also leverage statistical correlations in the input sequence even if they have no underlying biological basis, as also observed by Kantroo et al. (2025b). This is highlighted by our results on the pattern completion task. The sequences we construct are highly artificial: repeating blocks of tryptophan and tyrosine are unlikely to appear in natural proteins. A model that was attempting to either match the physics or evolutionary history of natural proteins would likely attempt to balance this highly hydrophobic sequence with hydrophilic residues. In contrast, we see in Figure 2C that the model prioritizes completing the pattern over producing a biophysically or evolutionary “realistic” protein.

Future directions include assessing whether in-context learning can be used as components in larger peptide screening and design pipelines. As our focus has been on establishing incontext learning as a phenomenon in PLMs, we have focused exclusively on “in-class” benchmarks, that compare with alternate learning approaches using the same PLM. This was done specifically to critically assess the power of in-context learning in these models in comparison with other use-strategies. Similarly, we have presented in-context learning as an alternative to fine-tuning approaches. In practice these approaches may instead be synergistic. Assessing the relative capability of MPEPs *vis-a-vis* models that explicitly operate on multiple sequences such as PoET (Truong & Bepler, 2025) or Metalic (Beck et al., 2025) would also help contextualize their importance to practical peptide-design pipelines.

### 4.1 Limitations

Several limitations are worth noting. First, MPEP performance is sensitive to which examples are chosen and (at small *k*) to how they are ordered. While large variance is a generic property of low-data classifiers, developing methods to identify representative demonstrations may improve robustness. At the same time, this sensitivity may present an opportunity: because MPEP classifiers can be instantiated without training, many distinct classifiers can be generated simply by permuting and sampling from the same set of context examples. Future work could explore bagging-style ensembles that aggregate predictions across multiple prompt orderings, potentially reducing variance due to context ordering while exploiting the diversity induced by different example arrangements. Second, the number of examples is bounded by the model’s context length. ESM-2 and ProGen2-xl both have a maximum context length of 1024 tokens. Quantized variants of ESM-2 with extended context length offer one path beyond this bound (Oliveira et al., 2024). Third, we have focused our experiments of peptides of fixed length. Extension and analysis of MPEP-based strategies on varying length peptides remains a direction for future work. Finally, we note that the “all-at-once” decoding we employ for masked language models decodes each peptide independently and consequently cannot capture higher-order correlations between amino acid sequences. While we have not observed this leading to a decrease in model performance, this nevertheless poses a limit on the performance of MPEP-based classifiers that use this strategy.

## 5 Conclusion

We have shown that single-sequence PLMs can classify peptides from a small number of in-context examples without any gradient updates. In our approach, we construct a multi-peptide example prompt by concatenating demonstration peptides with glycine spacers. Despite the fact that neither model has been trained on multiple protein sequences, both models are able to deduce shared properties between the concatenated prompts and use them to infer shared peptide properties. Across a synthetic pattern-completion task, secondary-structure classification, and MHC-II binding prediction, performance improves consistently with the number of examples and with model size, mirroring scaling trends for in-context learning in large language models. Moreover, MPEP-based classification outperforms PSSM approaches and achieves competitive performance with explicit learning on PLM embeddings while being three orders of magnitude faster. These results indicate that single-sequence PLMs implicitly support in-context inference and that MPEP conditioning represents an underexplored, lightweight approach to peptide classification in low-data settings.

## 6 AI Usage

LLMs were used to write parts of the code for experiments performed in this work. Text for this work was written by humans with feedback from LLMs. LLMs were used to help search for references and related work.

LLMs were used to assist with literature review, code development and debugging, analysis and interpretation, figure preparation, and manuscript revisions. The authors developed the MPEP framework, directed the research, and critically evaluated the findings. The authors take full responsibility for the methods, results, and final manuscript.

## A Secondary Structure classification

## B Effect of Example Choice and Order

Under the positive-only score, zero-shot performance varies substantially with model size: ESM-2 3B achieves ROC-AUC ≈ 0.71 while ESM-2 650M scores below 0.5. This variance reflects whether each model assigns higher unconditional probability to binder or non-binder compositions. To quantify this, we computed the Kullback-Leibler (KL) divergence between the amino-acid frequency distributions of the binder (*B*) and non-binder (*N*) test sets relative to the model’s unconditioned distribution (*M*, estimated from 15 uncontextualized mask tokens). The signed difference Δ = KL(*B* ∥*M*) − KL(*N* ∥ *M*) predicts the direction of the zero-shot bias (Table 2): models with Δ *<* 0 (8M, 150M, 3B) score zero-shot above 0.5 (0.67, 0.64, 0.71), while models with Δ *>* 0 (35M, 650M) score below or near 0.5 (0.56, 0.46), and the magnitude of Δ tracks the magnitude of the deviation. The same pattern explains the asymmetric zero-shot behavior observed on the secondary-structure task and clarifies that any peptide-scoring approach that uses raw PLM probabilities directly is implicitly conditioned on the model’s compositional prior.

**Table 2:**
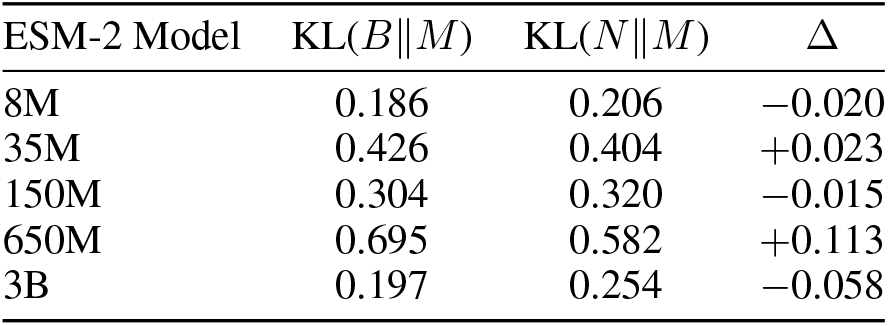
KL divergence from each ESM-2 model’s unconditioned amino-acid distribution (*M*) to the binder (*B*) and non-binder (*N*) test sets. Δ = KL(*B*∥ *M*) −KL(*N*∥ *M*). The sign of Δ predicts the direction of the zero-shot bias under the positive-only score; the difference score eliminates this dependence by construction.

**Table 3:** Comparative effect of removing attention heads; induction-head rows in bold. Control rows are the mean over three random draws of non-induction heads matched in count and layer. Copy NLL is the loss-per-residue on repeats 2-4 of a random sequence (random chance equates to ln 20 ≈ 3.0 approximately. ROC-AUC is for the difference score with *k* = 32 in-context examples, averaged over five random sequence draws.

| Model (params) | Heads ablated (ablated total) | copy NLL | ROC-AUC ($k = 32$) |
| --- | --- | --- | --- |
| small (151M) | none | 0.20 | 0.832 |
|  | **induction (5)** | **3.11** | **0.663** |
|  | layer-matched control (5) | 0.20 | 0.837 |
| medium (765M) | none | 0.19 | 0.862 |
|  | **induction (5)** | **3.05** | **0.661** |
|  | layer-matched control (5) | 0.20 | 0.856 |
| base (765M) | none | 0.20 | 0.839 |
|  | **induction (3)** | **3.06** | **0.736** |
|  | layer-matched control (3) | 0.19 | 0.841 |
| large (2.78B) | none | 0.21 | 0.846 |
|  | **induction (3)** | **2.97** | **0.780** |
|  | layer-matched control (3) | 0.21 | 0.848 |
| xlarge (6.44B) | none | 0.17 | 0.866 |
|  | **induction (4)** | **2.72** | **0.799** |
|  | layer-matched control (4) | 0.17 | 0.866 |

## C MHC-II binder classification

## D PepBenchmark generalization and ablation studies

We sought to assess the generalizability of our MPEP method on a wide range of tasks. However, labeled peptide datasets are sparse due to experimental time, cost, and throughput constraints. PepBenchmark is a recently curated benchmarking dataset intended to address this problem Zhang et al. (2026). However, our approach does not allow for direct utilization of PepBenchmark due to the heterogeneous peptide sequence lengths in each binary classification task. To assess MPEP for generalizability, we bin the sequences in PepBenchmark by task and length and report performance metrics on each of those bins.

We evaluated positive-only and difference scoring with intact examples, residue-scrambled examples, and label-mixed prompts on a fixed collection of PepBenchmark task–length bins. This section describes the data preparation, scoring, and results for these six experiments.

### D.1 Shared experimental methods

#### Data preparation and partitions

We started from 26 PepBenchmark binary clssification tasks, excluding the peptide–protein interaction task. The peptide-protein interaction dataset is out of scope for these experiments because each entry is a unique peptide-protein pair with corresponding binary label indicating yes/no binding. Within each eligible task, duplicate sequences retained their first occurrence and label, including conflicting duplicates. Sequences that were identical across tasks were retained only in the smallest task set. Retained sequences were binned by length. Bins required at least 500 sequences and at least 100 of each label. This resulted in 41 task-length bins covering eight tasks.

Each sequence-length bin was partitioned once, stratified by class. From each class, max(32, ⌈0.2, *n*_*c*_⌉) sequences (where *n*_*c*_ is the class size) were randomly selected to form a fixed training pool; all remaining sequences formed a fixed test set. For each *k* and repeat, *k* positive and *k* negative in-context examples were drawn without replacement from the training pool. Test sequences and labels were held fixed across prompt sizes, repeats, and conditions.

#### Prompts and scores

All PepBenchmark experiments used the ESM-2 650M model. MPEPs were made with *k* examples and *L* mask tokens appended to the prompt end of the prompt where *L* was the bin’s peptide length. The model’s log-softmax probabilities at those positions formed a lookup table. For a test peptide *x*, the positive-only score summed its residue log probabilities under the first prompt; the difference score subtracted the corresponding sum under the second prompt. Each prompt was checked separately against the 1,024-token limit using *k*(*L* + 5) + *L* + 2, including beginning- and end-of-sequence tokens. This excluded the *k* = 32 cells for antimicrobial lengths 29, 30, and 36.

#### Sampling and reporting

All sampling was seeded for reproducibility, with a separate seed for each task–length split and each example draw. We drew three independent sets of examples per cell, sampling without replacement within each class pool; the same example could appear in more than one draw, and the sets at larger *k* were not supersets of those at smaller *k*. For each draw we computed ROC-AUC on the fixed test set and then averaged the three values, rather than pooling predictions first. Only cells with all three repeats were reported. The heatmaps show these means to three decimals without uncertainty intervals. The descriptive averages below are computed from the displayed cell values, weighting the same 38 bins equally — the bins with *k* = 32 results — including when summarizing smaller *k*

### D.2 MPEP-ESM2 generalization and ablations on PepBenchmark

We began our investigation by assessing the generalizability of our MPEP method on the 41 PepBenchmark task–length bins. As previously observed, the difference scoring method (Fig. 17) outperforms the positive-only scoring method (Fig. 10). This we knew from previous experiments, and it has now been validated more thoroughly across eight different classification tasks. These baseline figures allow us to compare each of the ablation experiments that follow.

**Figure 10:**
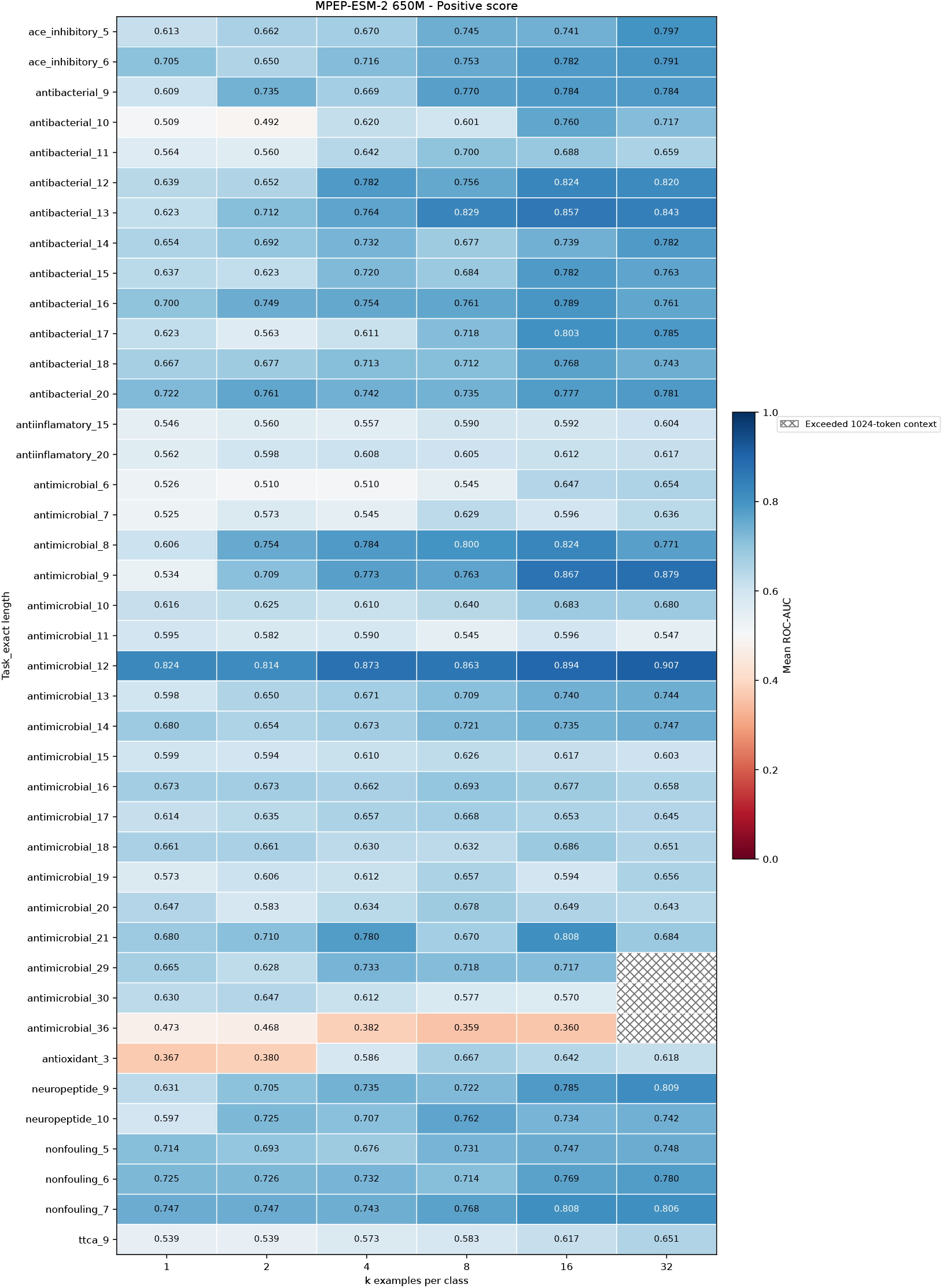
Positive-score MPEP-ESM2 650M PepBenchmark baseline. The heatmap shows classification performance using the positive-only scoring method as a function of number of examples in each prompt (columns) on 41 task–length bins from PepBenchmark binary classification tasks (rows). Each cell is the mean test ROC-AUC over three prompt draws. Hatched cells exceed the 1,024-token context limit.

#### Ablation 1: random examples drawn from UniProtKB/Swiss-Prot amino-acid frequencies

Each MPEP has a token length of *k*(*L* + 5) + *L* + 2. To rule out performance improvements due to increasing prompt length alone, we made MPEPs whose examples were randomly sampled from the amino-acid distribution of UniProtKB/Swiss-Prot (2026-03 release) and assessed the resulting classifiers on our binned PepBenchmark dataset. For the positive-only scoring method, absolute classification performance remains above random chance, with *δ*ROC-AUC = |ROC-AUC − 0.5 | ≈0.1–0.2, indicating that some class-specific statistical signal is present for some examples. However, context-length-dependent classification performance remains qualitatively flat (Fig. 11). This behavior starkly contrasts with the results seen in the baseline MPEP–PepBenchmark experiment, and it also contrasts with the behavior seen in our other experiments. This indicates that performance gains are not due to prompt length alone, and that MPEPs provide class-specific signal to the ESM protein language model.

**Figure 11:**
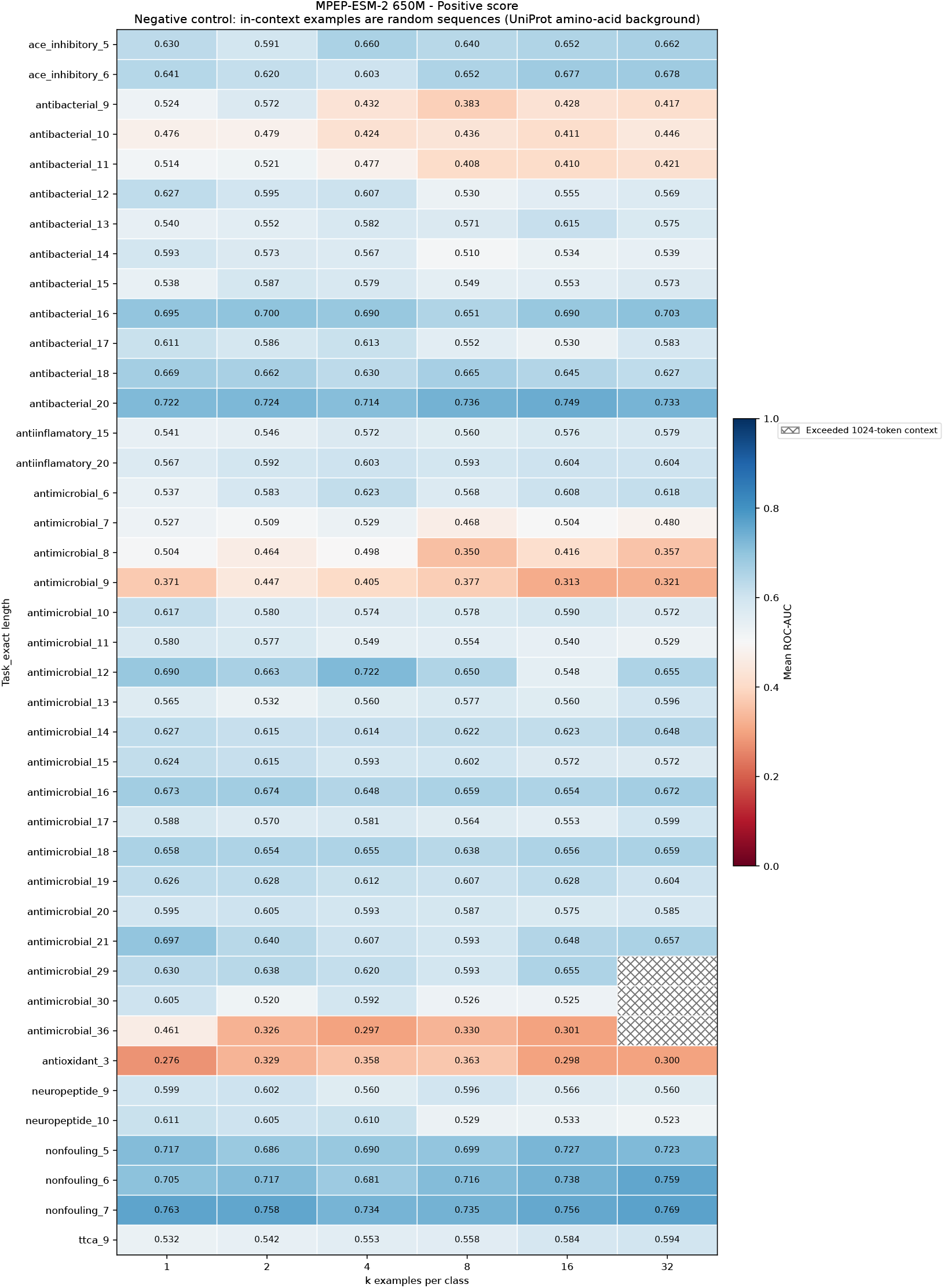
Positive-score MPEP-ESM2 650M PepBenchmark UniProtKB/Swiss-Prot randomized example ablation. Each peptide example is a random sequence whose residues are drawn i.i.d. from the amino-acid background frequency of the UniProtKB/Swiss-Prot database (2026-03 release).

**Figure 12:**
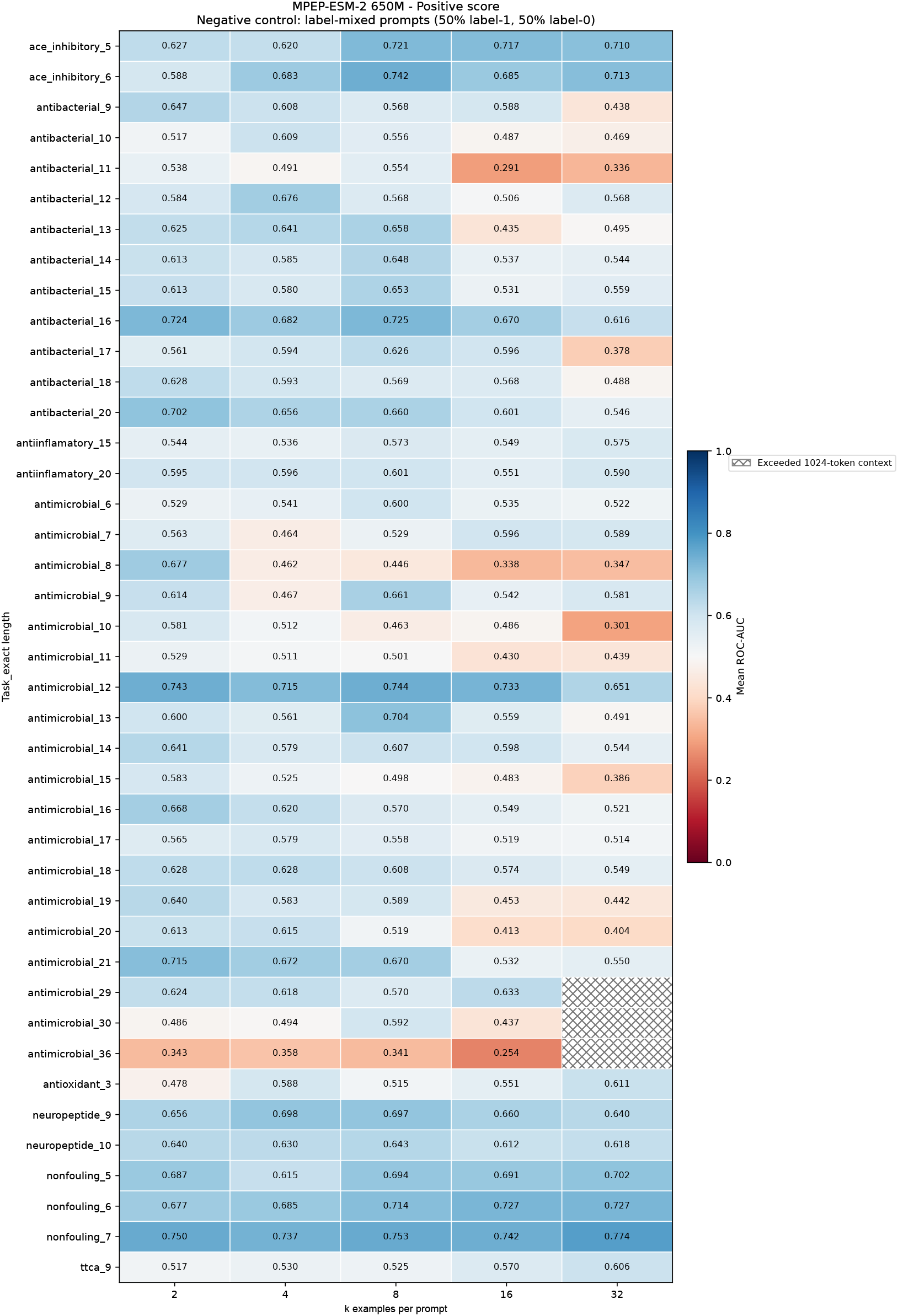
Positive-score MPEP-ESM2 650M PepBenchmark label mixing ablation. Each size-*k* prompt contains *k/*2 true positives and *k/*2 true negatives in randomized order (*k* = 1 was omitted for this reason). Each cell averages three prompt draws evaluated with unchanged test sequences. Label-mixing decreases classification performance relative to baseline MPEP 10.

Additionally, the greater-than-chance classification performance of the positive-only scoring method indicates that the background amino-acid distribution of UniProtKB/Swiss-Prot may overlap with the positively or negatively labeled examples in the PepBenchmark dataset, though further investigation into the background similarity of the reference dataset is out of scope for this work.

For the difference scoring method, both absolute and context-length-dependent performance fall to near chance across all tasks and all numbers of prompted examples (Fig. 18). This provides empirical evidence that the difference score cancels out redundant statistical signal from the prompt.

Overall, these data show that MPEPs provide class-specific signal to the model, and that the improvement in classification performance due to the use of MPEPs is attributable to more than the length of the prompt alone.

#### Ablation 2: label mixing

Our method depends on the assumption that building an MPEP with examples of peptides that share a common activity confers an implicit class labeling. To test that assumption, we generated MPEPs in which half of the examples are positively labeled and half are negatively labeled. We omit *k* = 1 in these experiments, as such prompts represent either a positive or a negative prompt, but not both. Difference scoring performance falls to near random chance for most tasks (Fig. 19). This result is expected, since two MPEPs with an equal number of positive and negative examples carry roughly the same statistical information; thus their difference is expected to be near zero by construction. Label mixing reduces positive-only scoring performance compared to baseline on most tasks, though classification performance remains above random chance for several tasks (Fig. 12).

#### Ablation 3: example permutation

Peptides are sequence-defined biopolymers whose function depends on interactions between specific residues and the interacting substrate Cascarina & Ross (2025). To assess the sensitivity of our method to the arrangement of residues within the prompt— as opposed to their composition alone—we permute each example sequence independently and then generate the MPEPs as before. Permutation preserves the length and amino-acid composition of every example exactly, so any change in performance is attributable to the loss of position-specific information. Positive-only scoring is largely unaffected by this manipulation (Fig. 13), whereas difference scoring loses performance in 35 of 38 matched bins, lowering the mean ROC-AUC at *k* = 32 by 0.070 (Fig. 20). Substantial discrimination nevertheless survives composition-preserving scrambling under both scoring methods.

**Figure 13:**
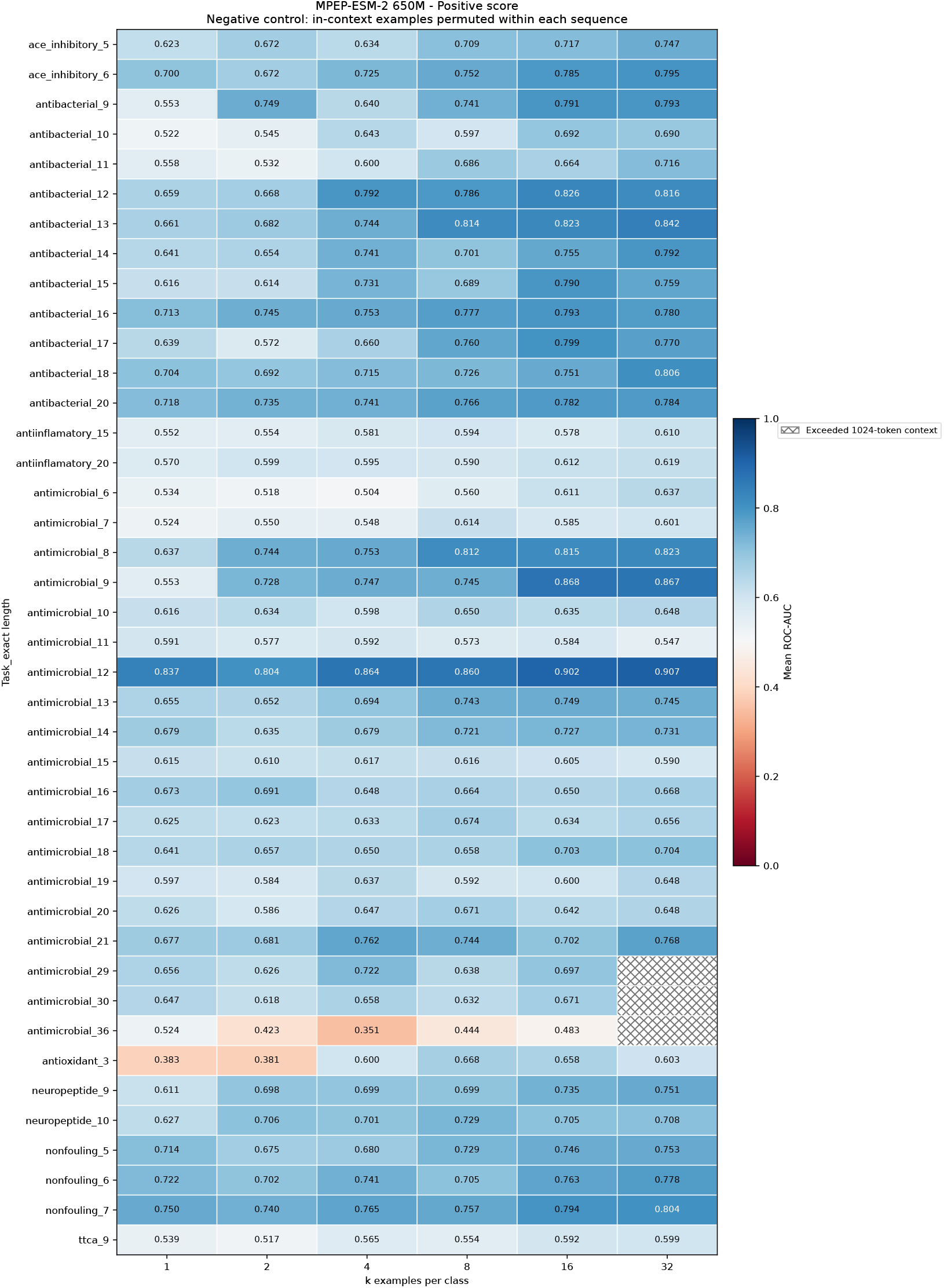
Positive-score MPEP-ESM2 650M PepBenchmark example permutation ablation. The heatmap shows classification performance as a function of number of examples in each prompt (columns) on 41 task–length bins from the PepBenchmark binary classification tasks (rows). Each example was independently permuted, preserving its length and composition. Each cell averages three prompt draws with permutation. Performance does not decrease significantly compared with baseline.

#### Ablation 4: PPM sampling, class composition sampling, and their difference

To further investigate the role of amino-acid position versus composition on MPEP classification performance, we performed the following experiment. We calculated a position probability matrix (PPM) from the training set of each class for each task. This [*L*, 20] matrix encodes position-specific statistical information about each class. For each *k*, we built 50 positive MPEPs and 50 negative MPEPs by sampling 50 \**k* positive sequences and 50 \**k* negative sequences from the PPMs generated from positive and negative examples, respectively. We then used those MPEPs to score the held-out test set for each task; those results are given in Figs. 14 and 21.

**Figure 14:**
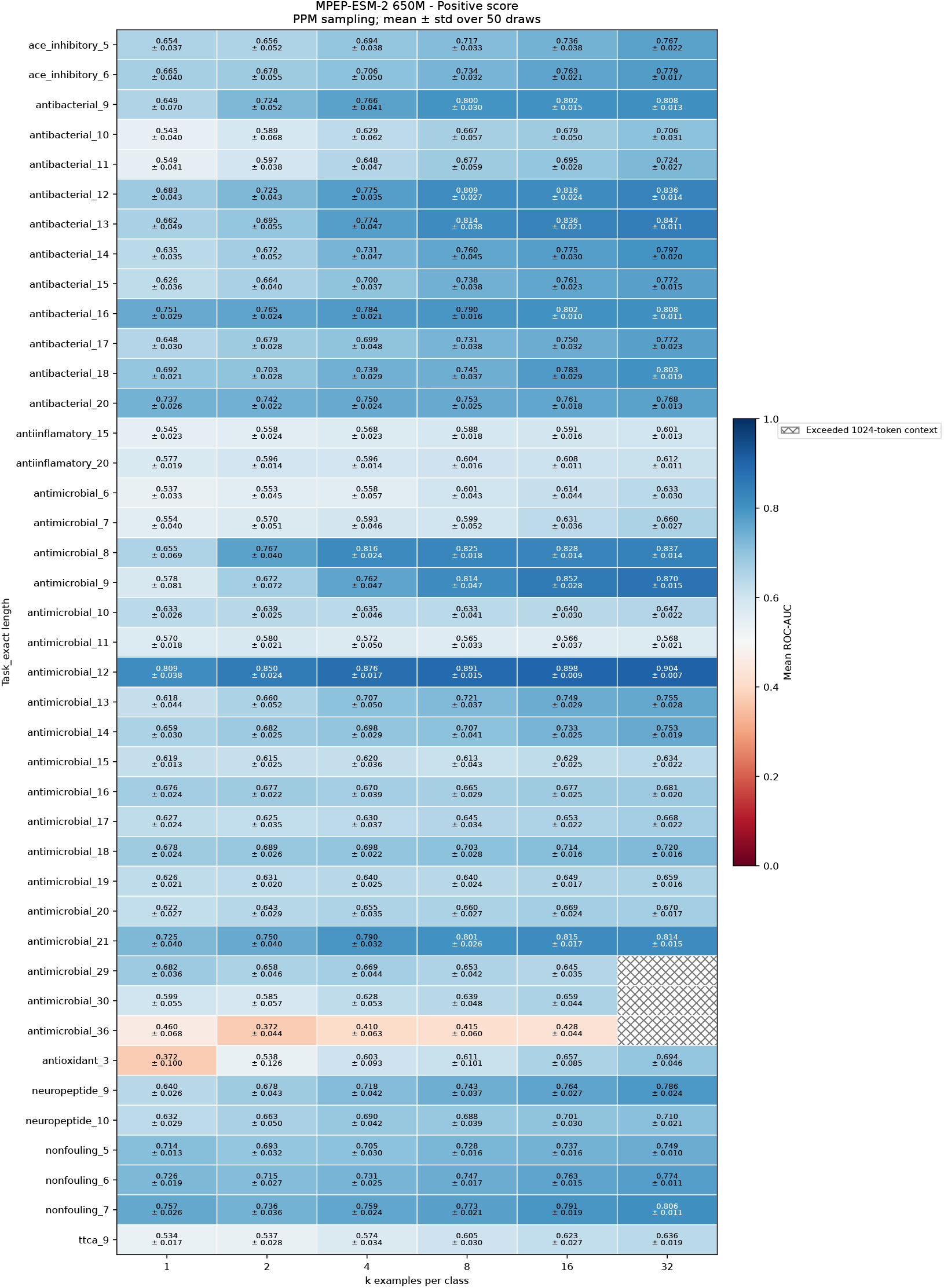
Positive-score MPEP-ESM2 650M PepBenchmark position probability matrix (PPM) sampling ablation. A PPM of size [*length*, 20] was calculated using the peptide sequences with label=1 for each task–length training set. MPEPs were built with examples sampled from each PPM. Therefore, examples maintain both class composition *and* position information. Each cell represents 50 MPEPs of length *k* whose examples were drawn from their respective PPM.

**Figure 15:**
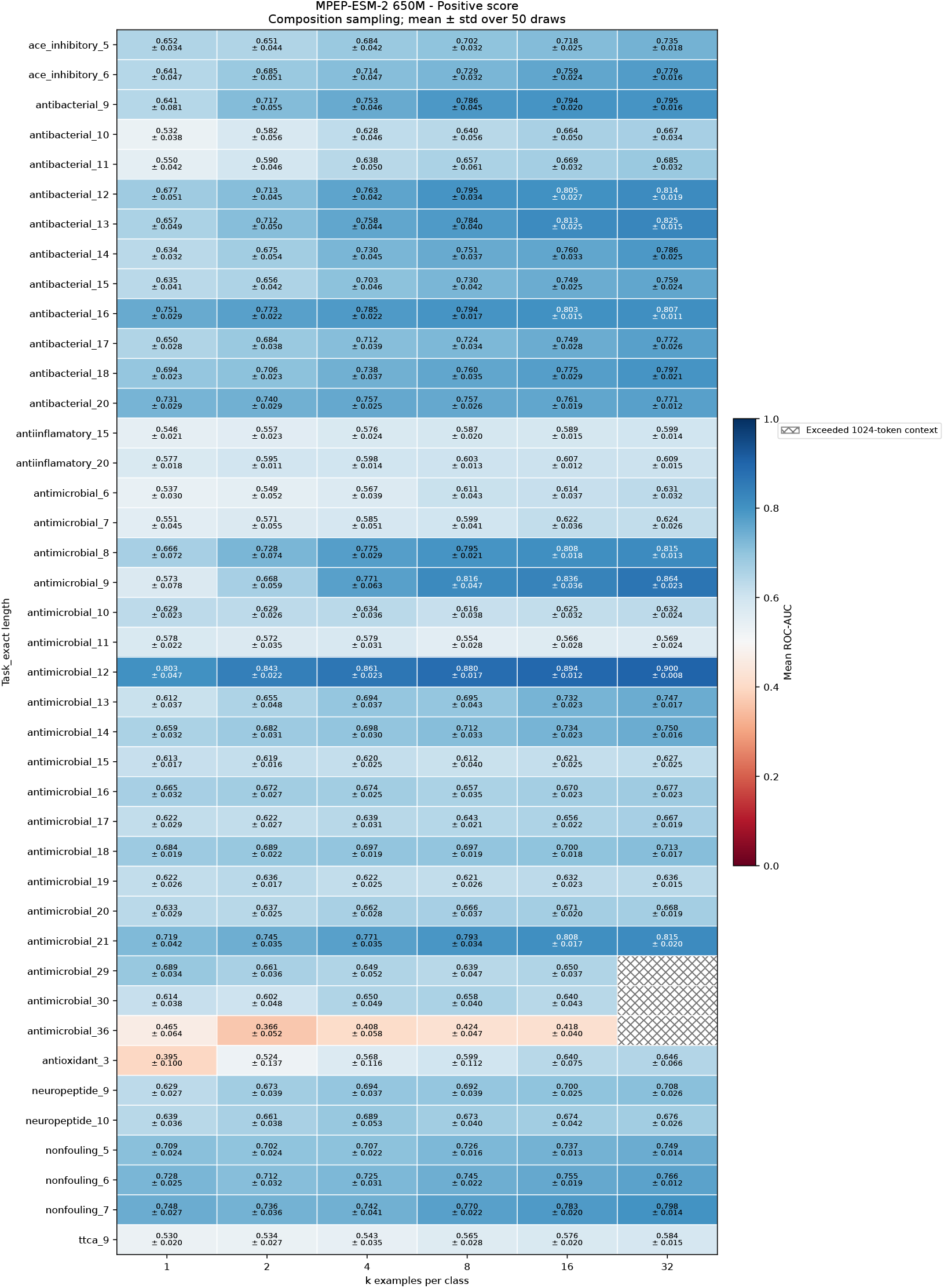
Positive-score MPEP-ESM2 650M PepBenchmark class composition ablation. A probability vector of size [1, 20] was calculated using the peptide sequences with label=1 for each task– length training set. MPEPs were built with examples sampled from each probability vector. Examples maintain overall class compositional, but not positional, information. Each cell represents 50 MPEPs of length *k* whose examples were drawn from their respective probability vector.

**Figure 16:**
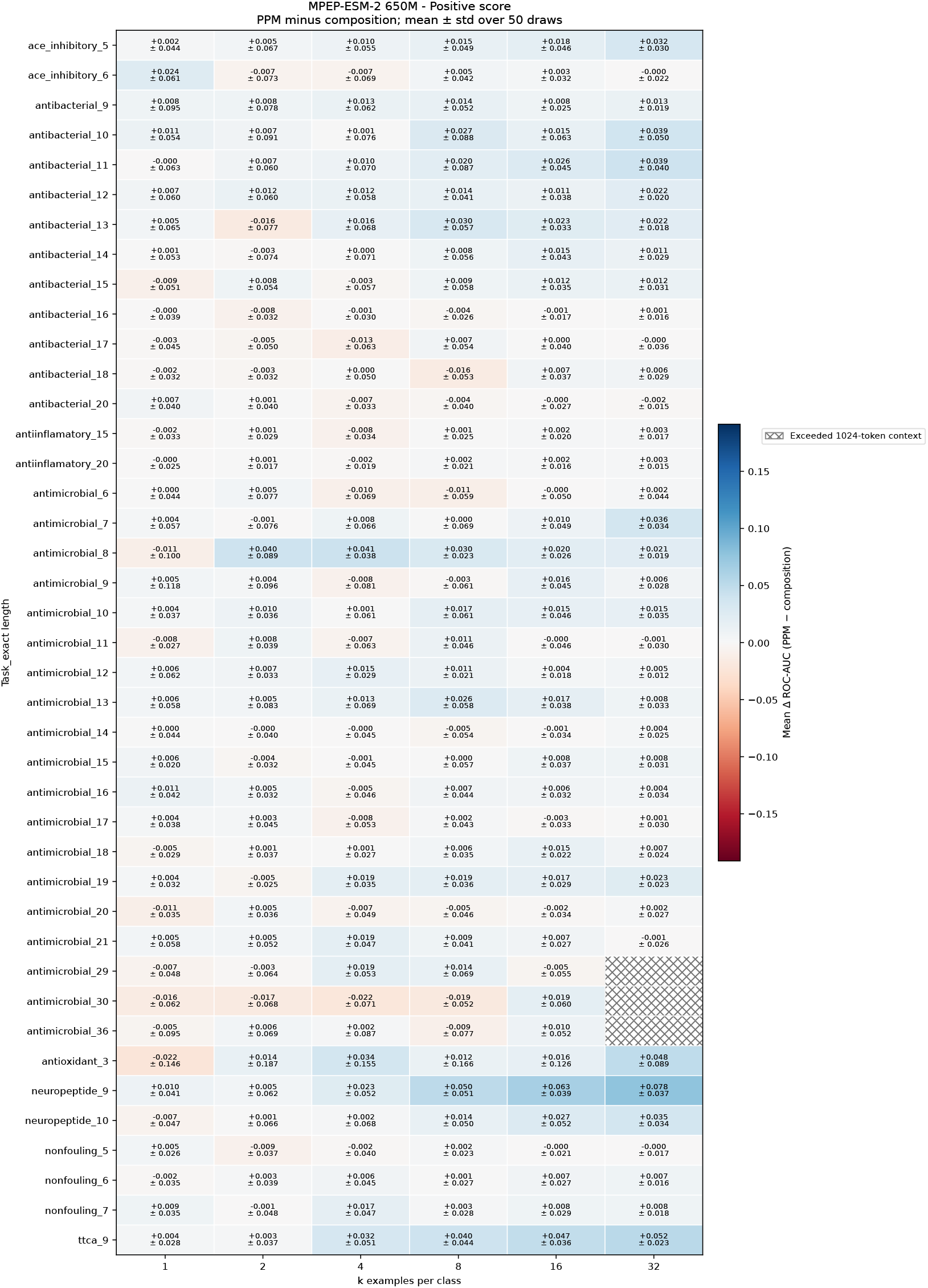
Heatmap of ROC-AUC_*Position*_ - ROC-AUC_*Composition*_ for each assessed PepBench-mark task–length bin. MPEP-ESM positive scoring does not benefit from the additional position-specific information encoded by the PPM.

**Figure 17:**
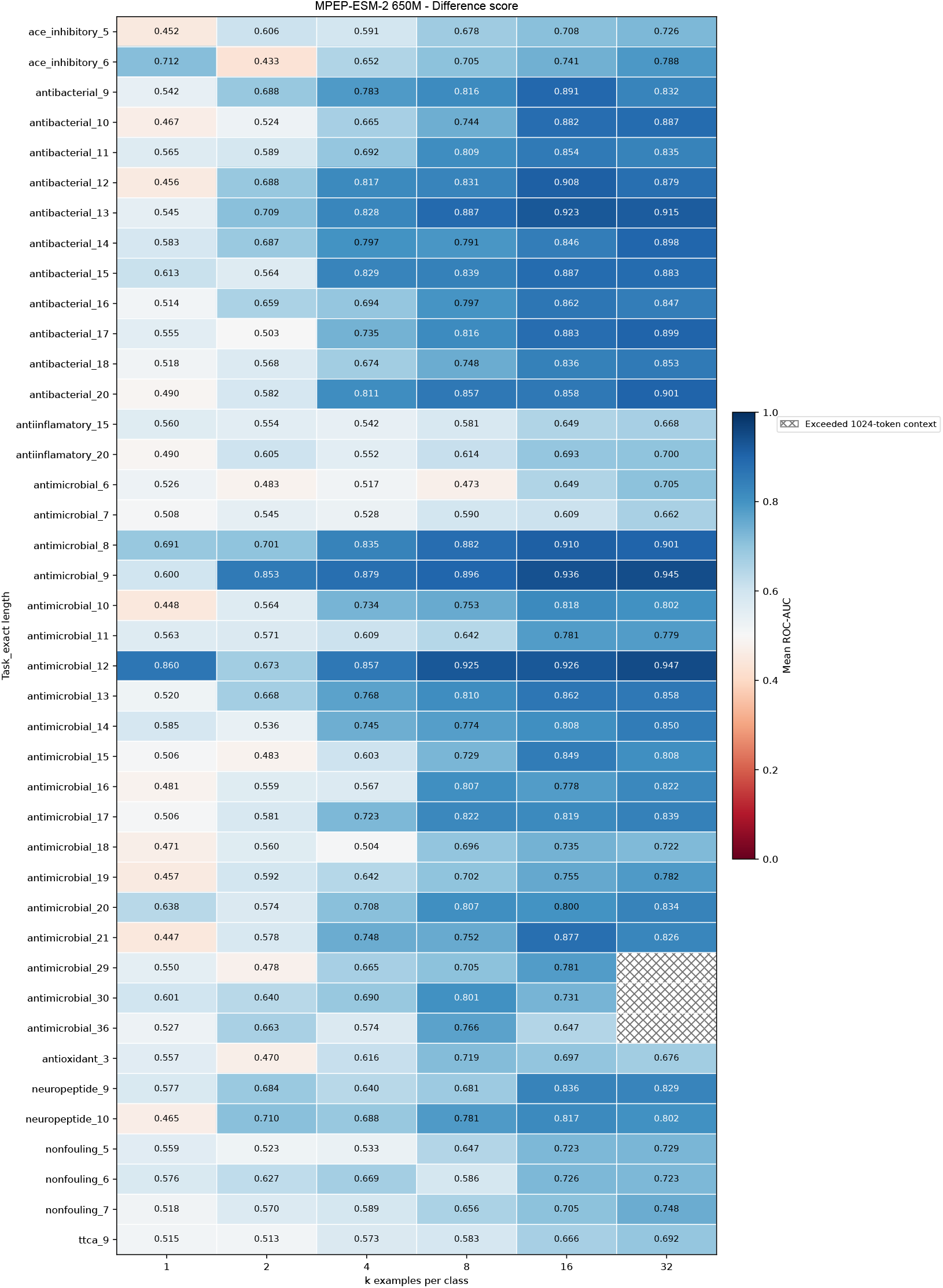
Difference-score MPEP-ESM2 650M PepBenchmark baseline. Separate prompts contain *k* positive and *k* negative examples; their log-probability sums are subtracted. Each cell is the mean test ROC-AUC over three prompt draws.

**Figure 18:**
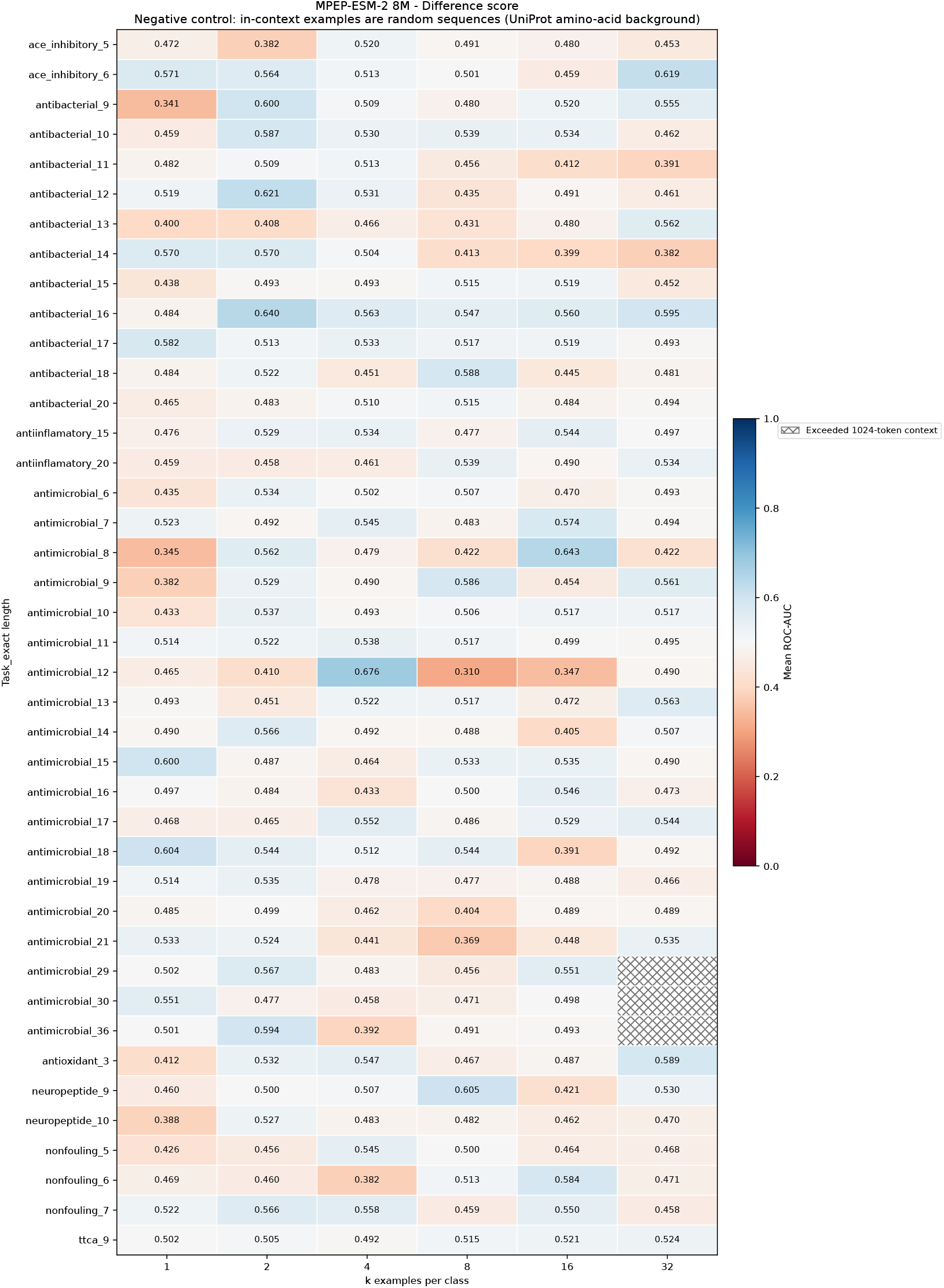
Difference score MPEP-ESM2 650M PepBenchmark UniProtKB/Swiss-Prot randomized example ablation. Each peptide example is a random sequence whose residues are drawn i.i.d. from the amino-acid background frequency of the UniProtKB/Swiss-Prot database (2026-03 release).

**Figure 19:**
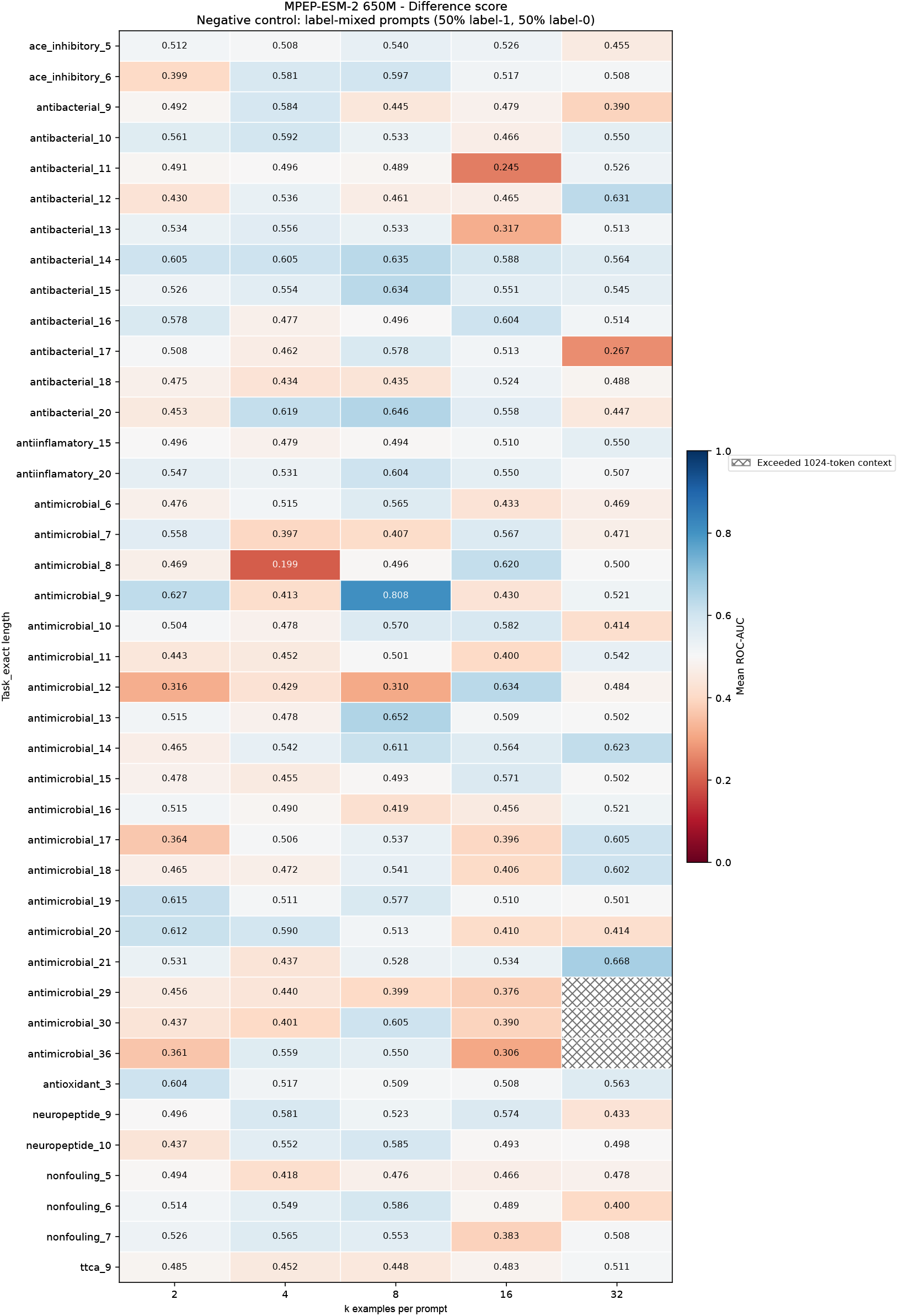
Difference score MPEP-ESM2 650M PepBenchmark label mixing ablation. Each size-*k* prompt contains *k/*2 true positives and *k/*2 true negatives in randomized order (*k* = 1 was omitted for this reason). Each cell averages three prompt draws evaluated with unchanged test labels.

**Figure 20:**
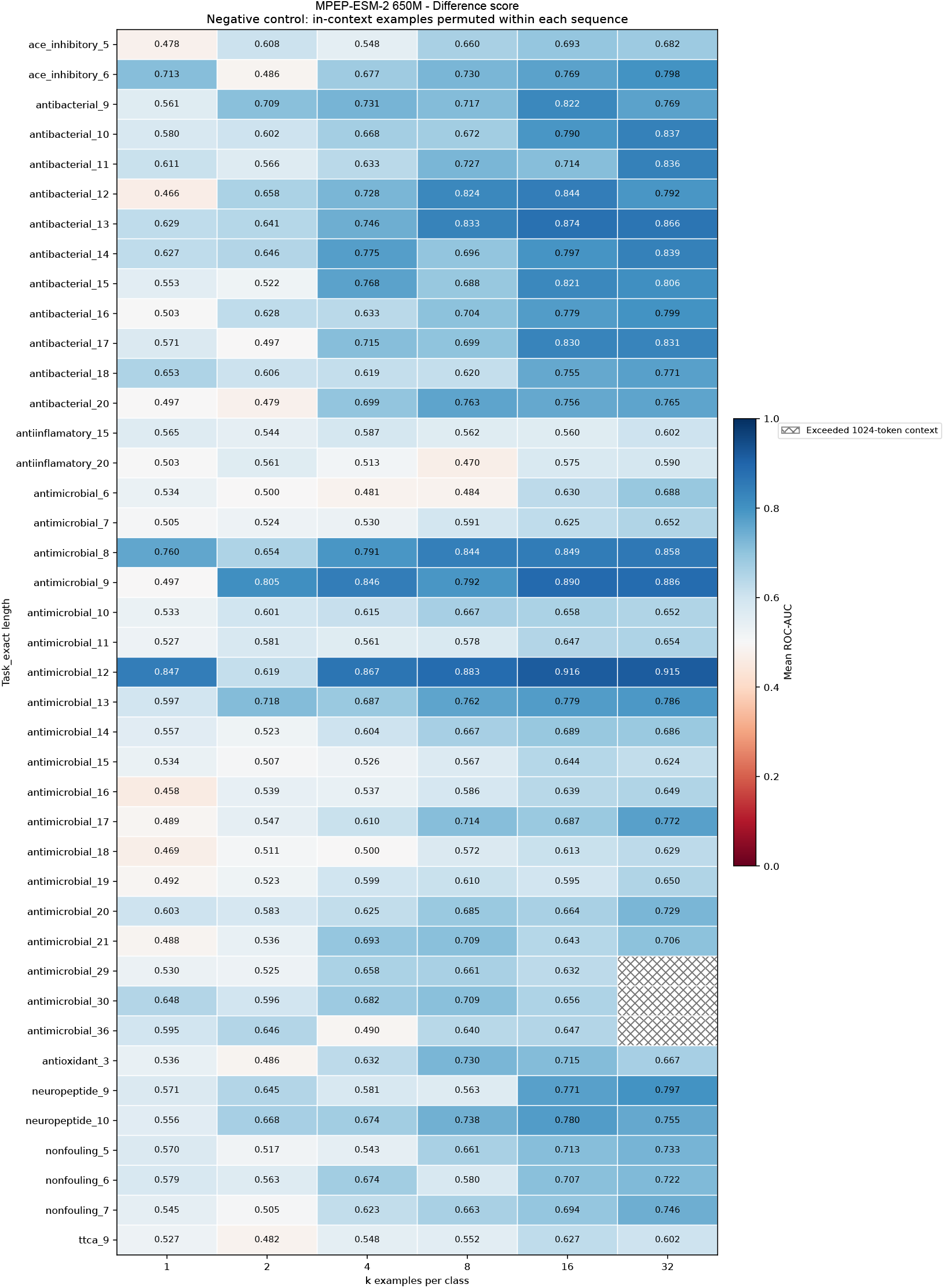
Difference-score MPEP-ESM2 650M PepBenchmark example permuation ablation. Class-specific examples were permuted prior to forming the MPEP, preserving example length and composition. Each cell averages three prompt draws with permutation; queries were unchanged.

**Figure 21:**
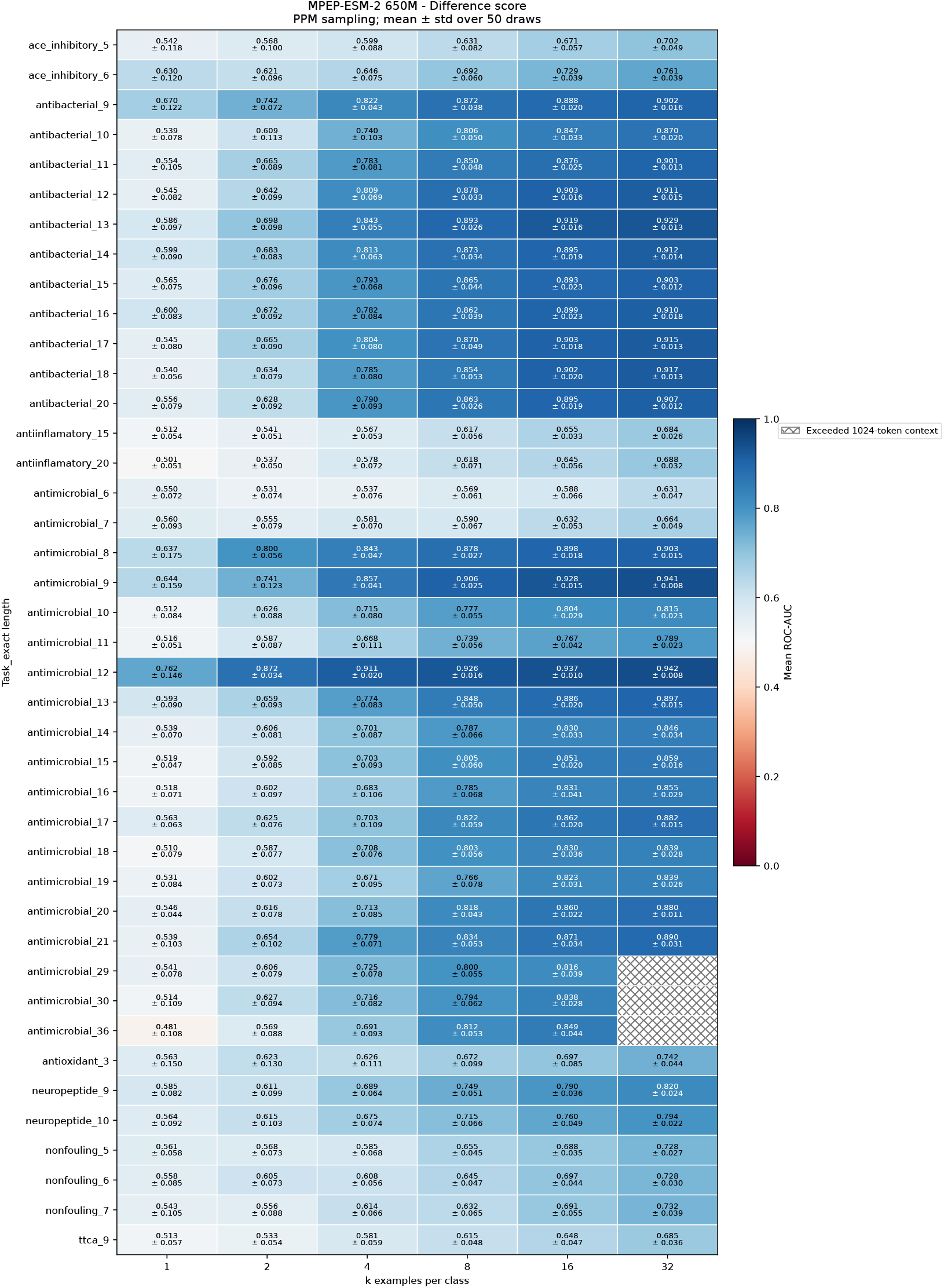
Difference score MPEP-ESM2 650M PepBenchmark position probability matrix (PPM) sampling ablation. Two PPMs of size [*length*, 20] were calculated for each task–length training set; one using the peptide sequences with label=1 and another using peptide sequences with label=0. 50 Positive and 50 negative MPEPs were built with examples sampled from each respective PPM. Therefore, examples maintain both class composition *and* position information. Each cell represents 50 MPEPs of length *k* whose examples were drawn from their respective PPM.

Next, we calculated a class composition probability vector of size [1, 20] by averaging the probabilities across the length of each PPM. This vector represents the overall amino-acid composition probabilities of the training examples in each class. Again, we built 50 positive MPEPs and 50 negative MPEPs by sampling 50 \**k* positive and 50 \**k* negative sequences from the class composition vector, and assessed classification performance (Figs. 15 and 22). Finally, we take the difference between the ROC-AUC values in each cell when prompted with PPM-generated MPEPs and with class-composition-vector MPEPs. The heatmap of ROC-AUC_Position_ − ROC-AUC_Composition_ effectively represents how well each scoring method can take advantage of amino-acid positional information, as the overall compositional information is the same for both PPM- and class-composition-sampled MPEPs. Across the 41 task–length bins, the positive-only scoring method yields little classification improvement from the additional positional information, with *δ*ROC-AUC *<* 0.08 across all tasks (Fig. 16). Remarkably, the difference scoring method shows large improvements in classification performance with the addition of positional information (Fig. 23) with *δ*ROC-AUC in the range of 0.1-0.2.

Taken together, these results indicate that the two scoring methods are not reading the same thing. Subtracting a matched negative prompt removes the signal the two prompts share—the model’s compositional prior and the prompt’s bulk amino-acid statistics—and what remains is the position-specific structure that actually separates the classes. In short: the positive-only score mostly reads *which* amino acids a peptide contains, while the difference score reads *where* they are.

**Figure 22:**
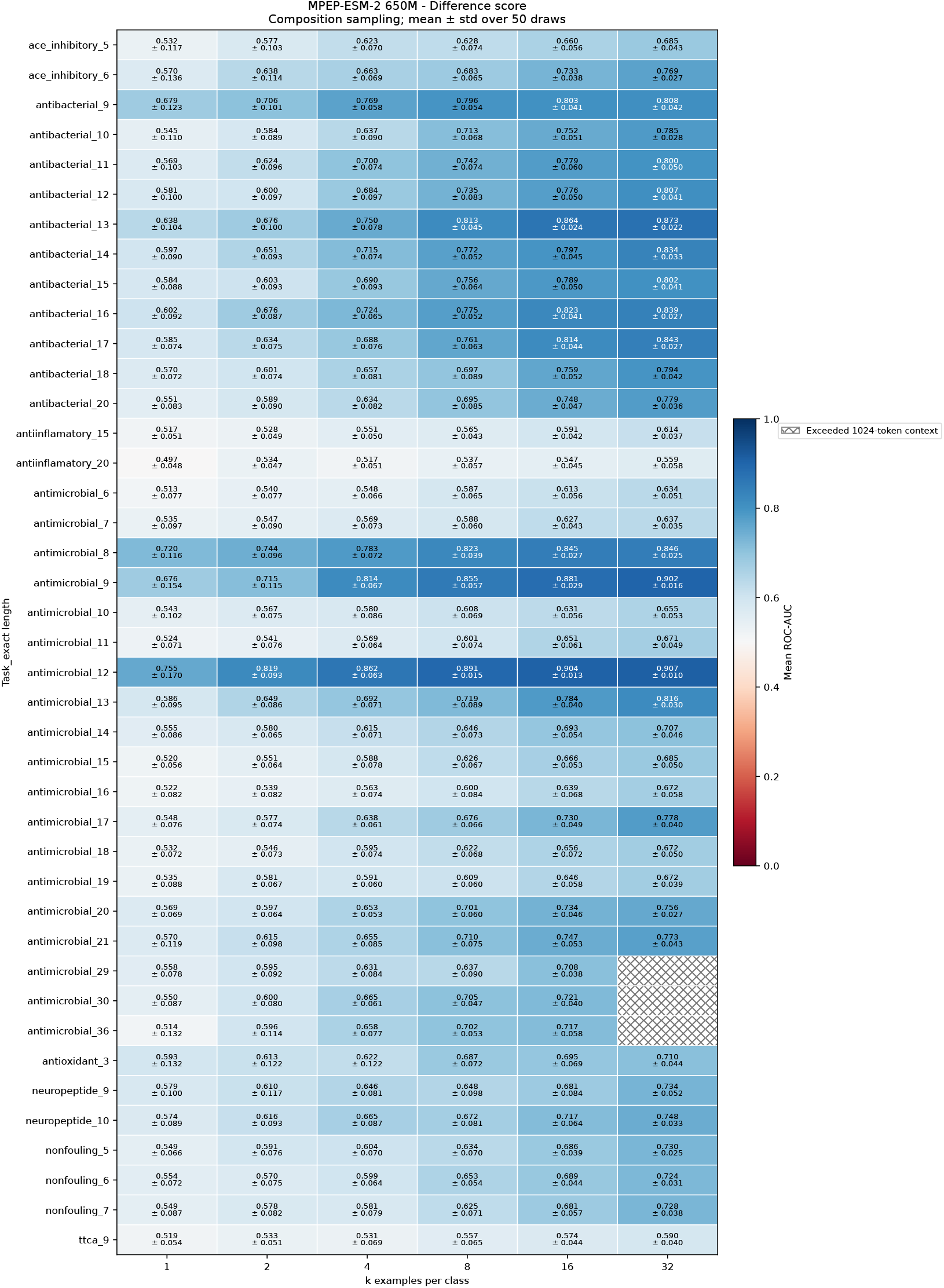
Difference score MPEP-ESM2 650M PepBenchmark class composition ablation. Two probability vectors of size [1, 20] were calculated for each task–length training set; one using the peptide sequences with label=1 and another using peptide sequences with label=0. 50 Positive and 50 negative MPEPs were built with examples sampled from each respective probability vector. Examples maintain class compositional, but not positional, information. Each cell represents the average and standard deviation of classification performance over 50 MPEPs.

**Figure 23:**
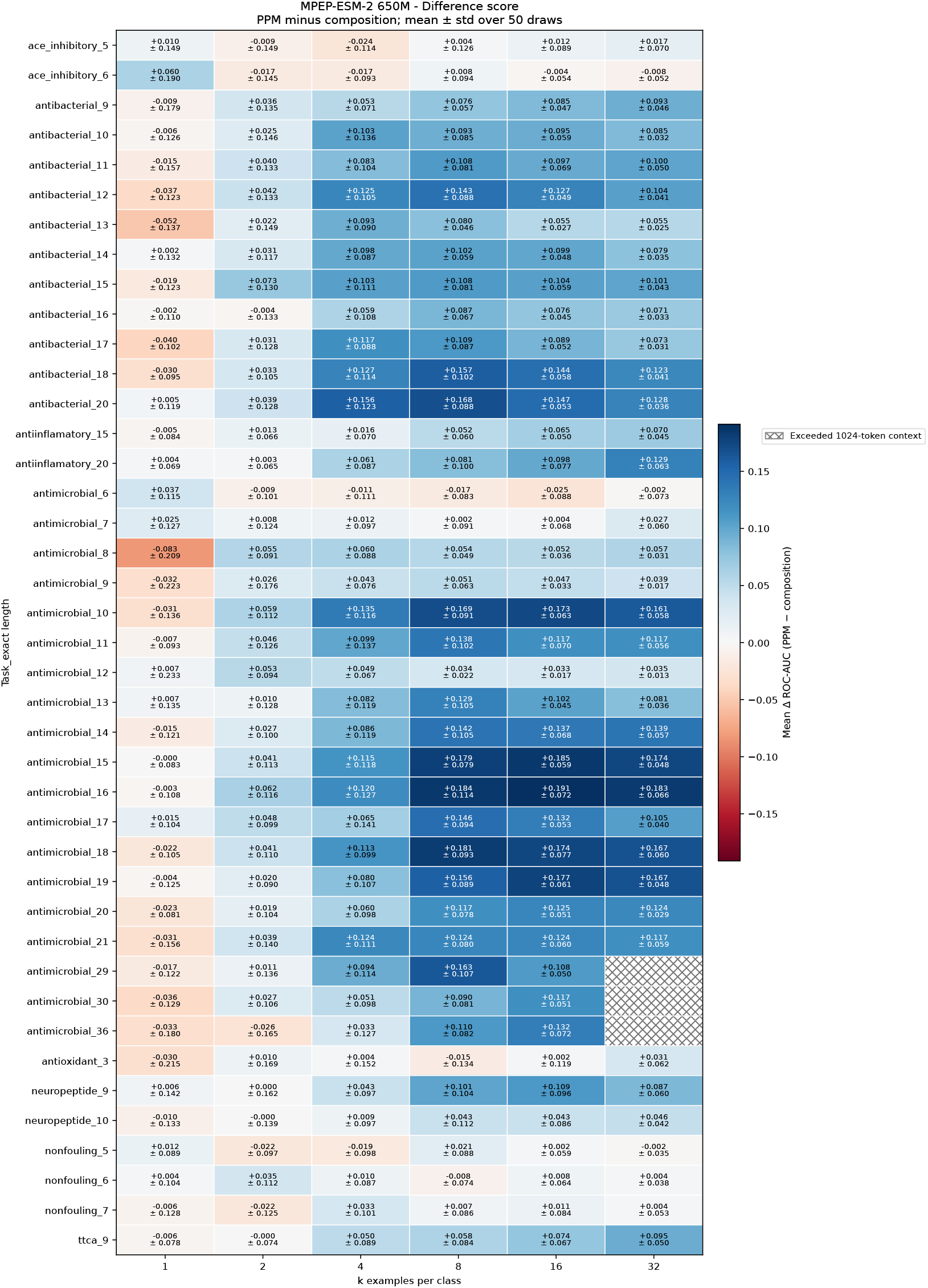
Heatmap of ROC-AUC_*Position*_ - ROC-AUC_*Composition*_ for each assessed PepBench-mark task–length bin using the difference score. MPEP-ESM difference scoring benefits from the additional position-specific information encoded by the PPM.

## E Induction heads in ProGen2

### E.1 ProGen2-small (151M)

**Figure 24:**
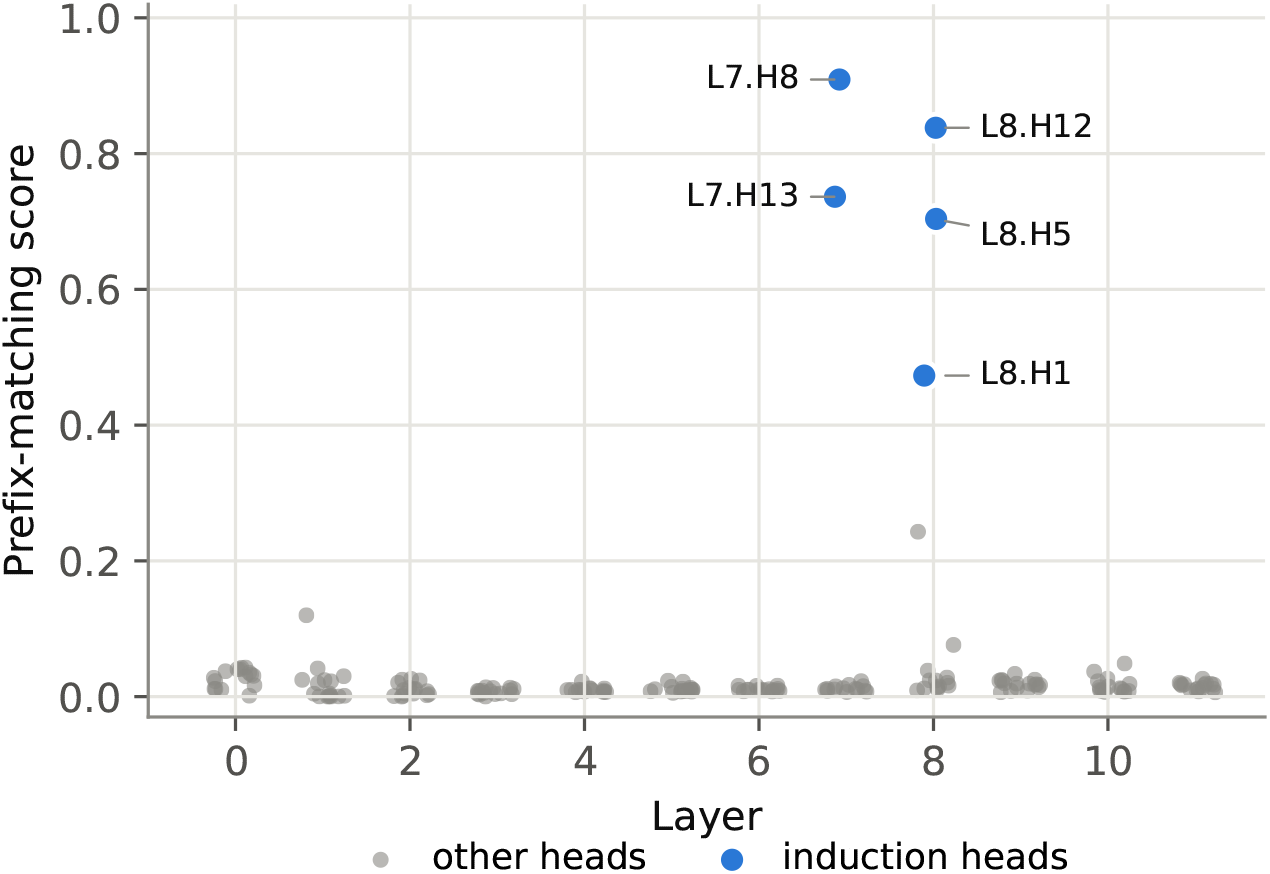
Prefix matching scatter in layers surrounding induction heads of ProGen2-small.

**Figure 25:**
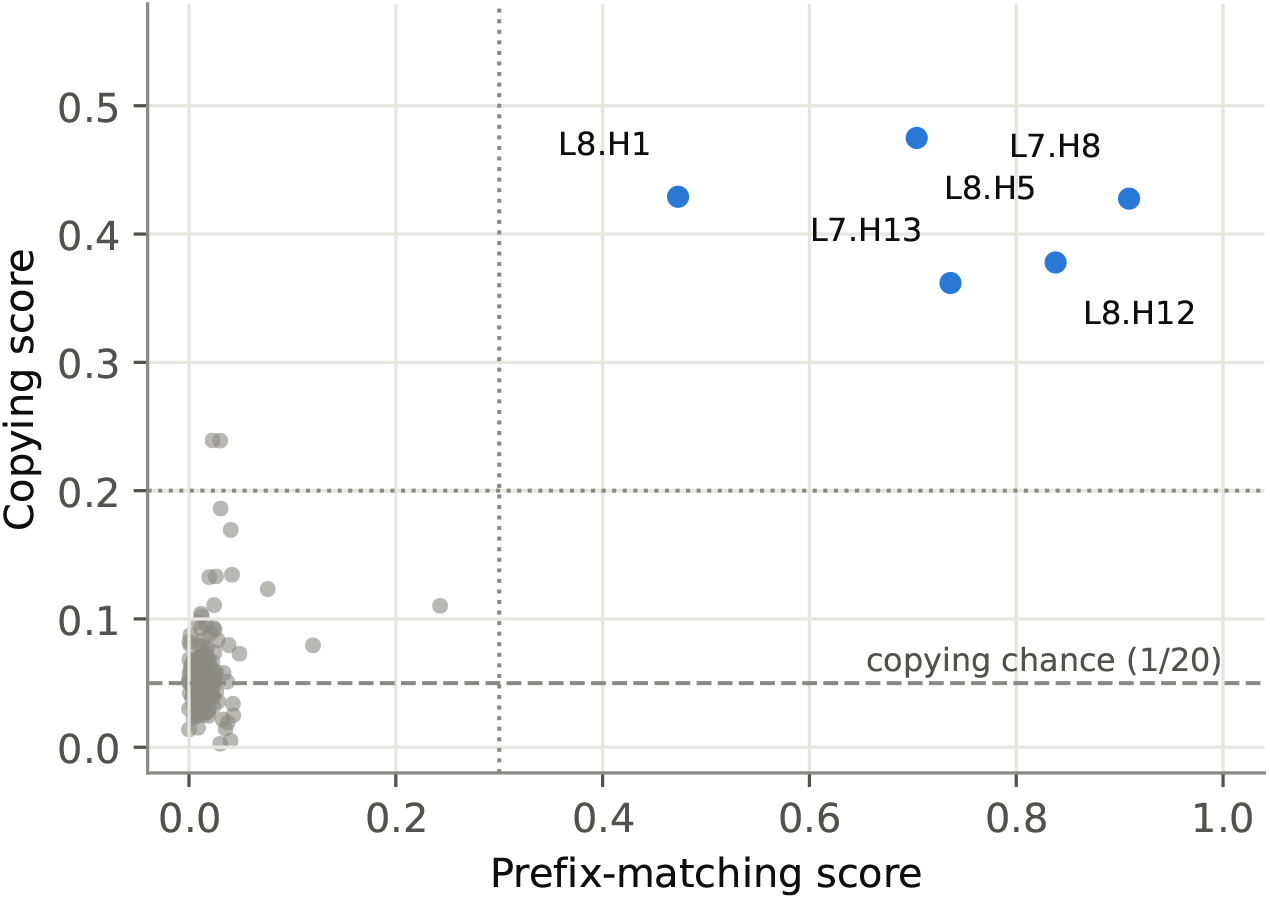
Prefix matching and copying scores of all heads in ProGen2-small.

**Figure 26:**
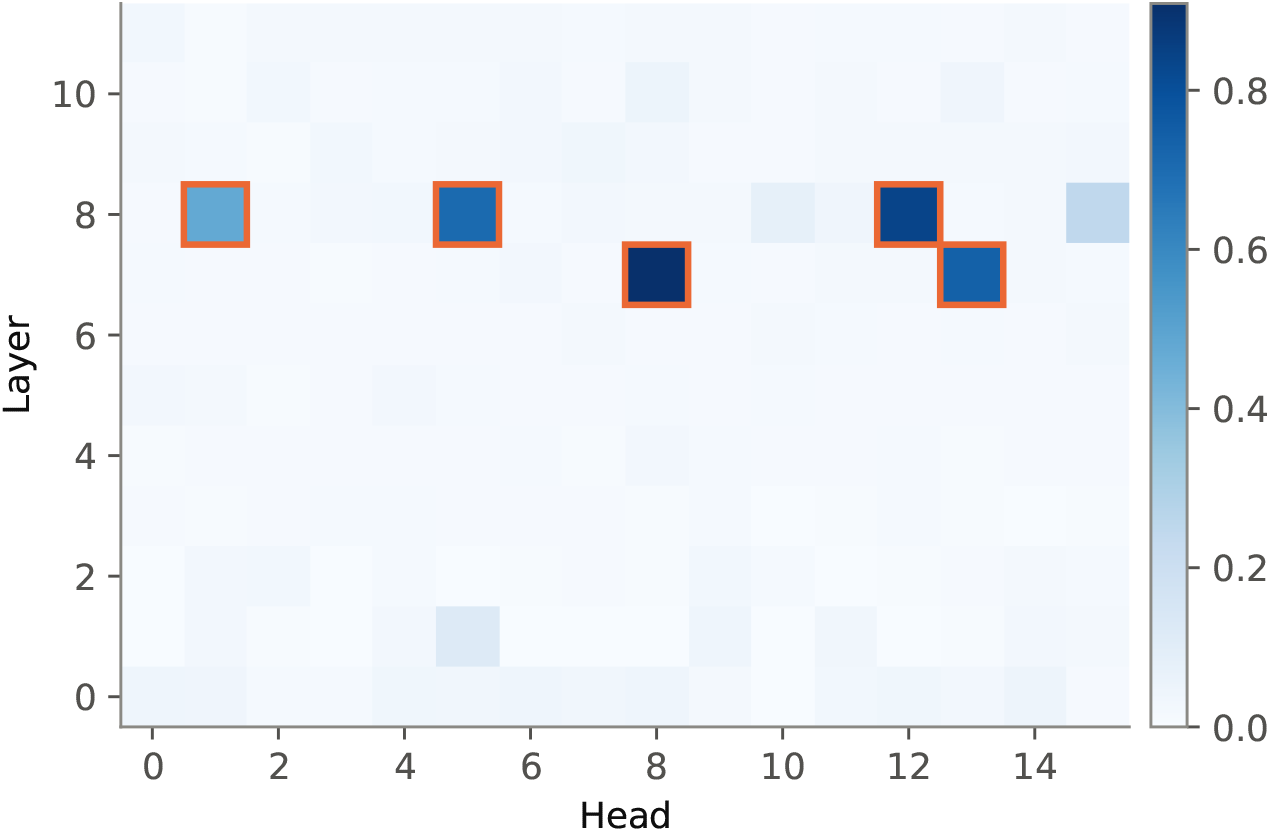
Prefix matching heatmap of induction-head and nearby layers in ProGen2-small. Induction heads are highlighted with orange border.

**Figure 27:**
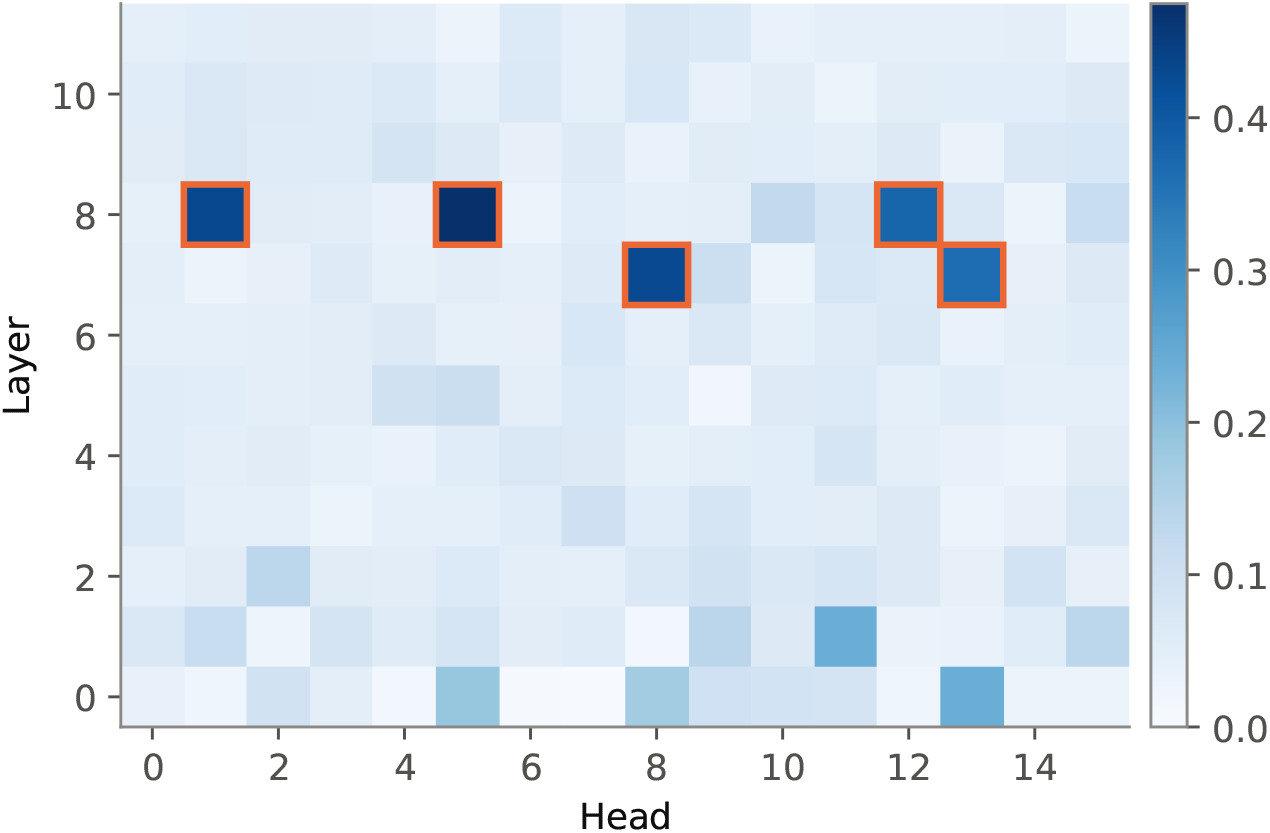
Copying heatmap of induction-head and nearby layers in ProGen2-small.

**Figure 28:**
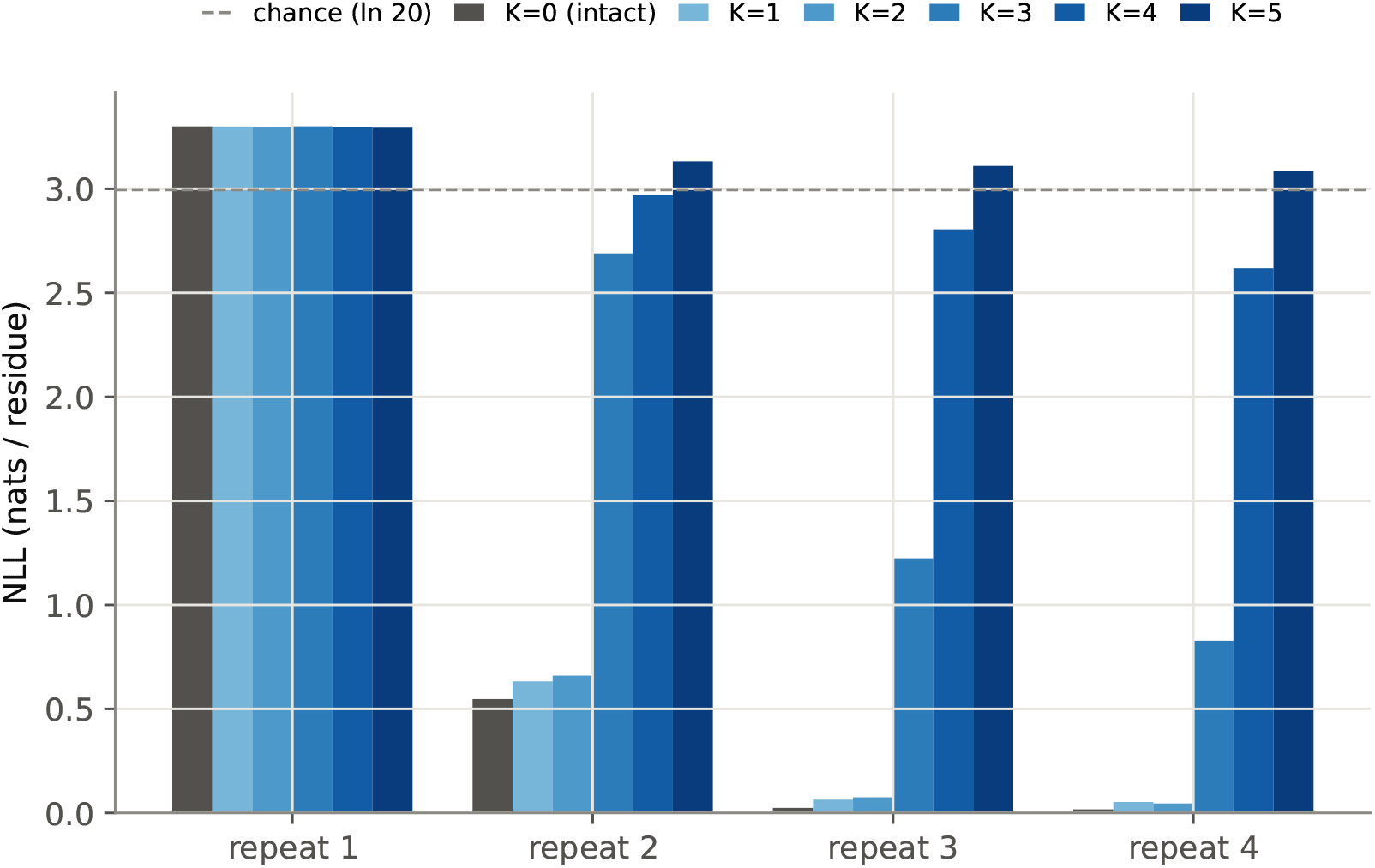
NLL loss-per-residue across all random sequence repeats during ProGen2-small inference, with *K* induction head(s) zero-ed out.

**Figure 29:**
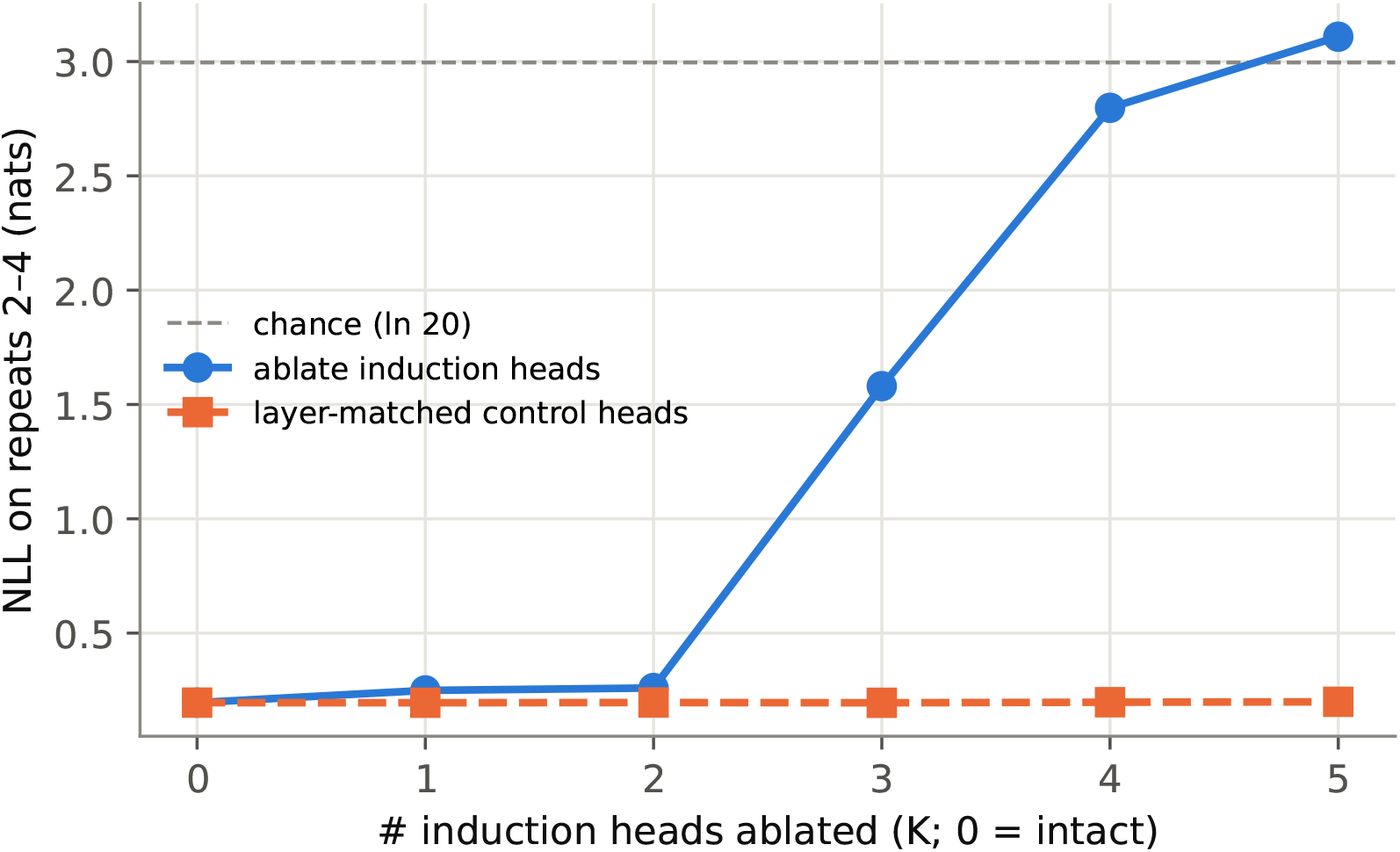
NLL loss-per-residue across random sequences repeats 2-4 between induction head ablation and control (ablating the same number of random, non-induction heads in the same induction layers) during ProGen2-small inference.

**Figure 30:**
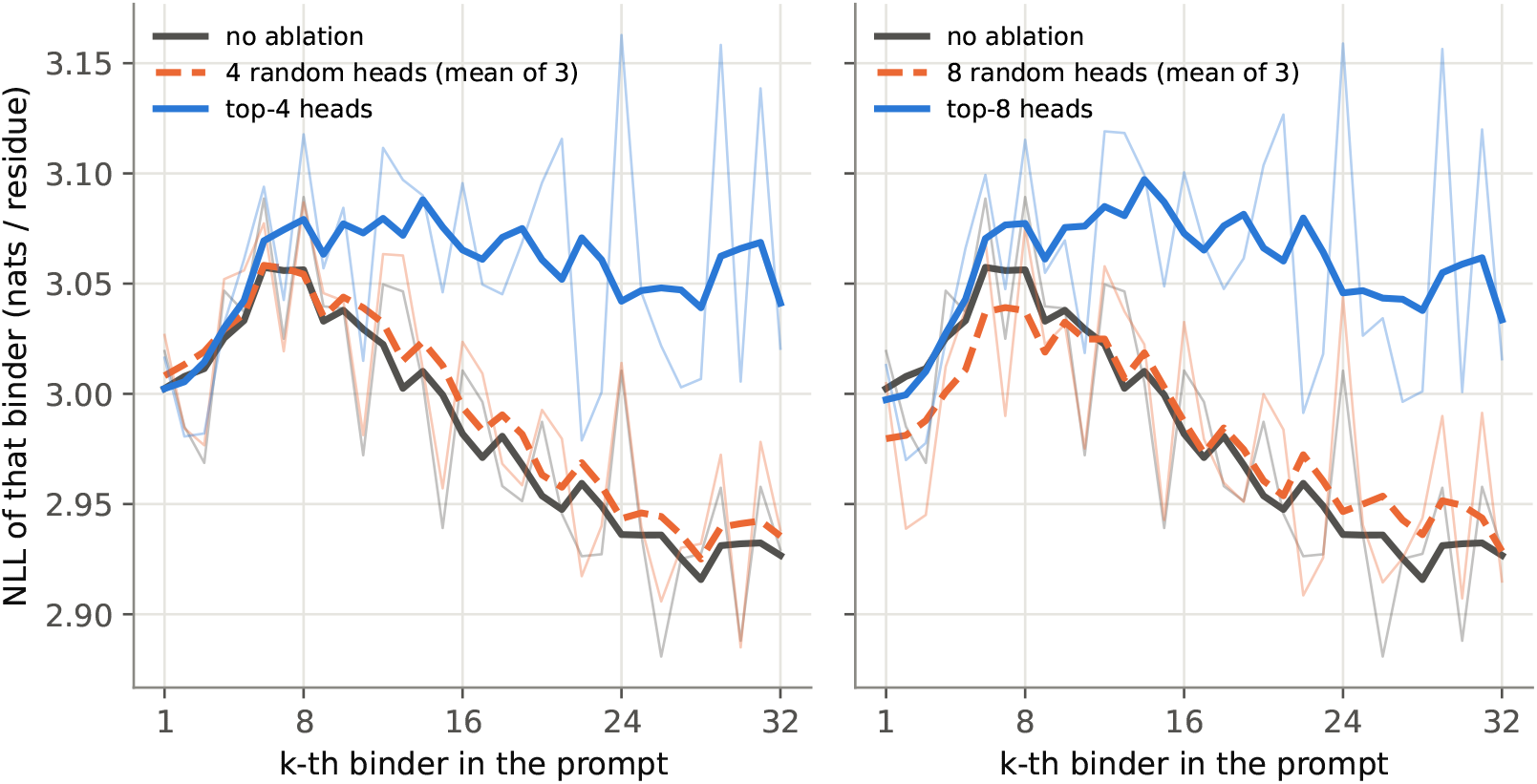
NLL loss-per-residue for each 15-mer example, averaged over 10 ProGen2-small inference using prompts constructed from 32 binder examples. Three parallel schemes are presented: no ablation, top-*K* heads and *K* random heads sampled from the same layers as top-*K* (averaged over 3 independent sampled set of heads). *\*Note*: NLL axis in this plot is positive (opposite to Fig. 6B); here, lower is better.

**Figure 31:**
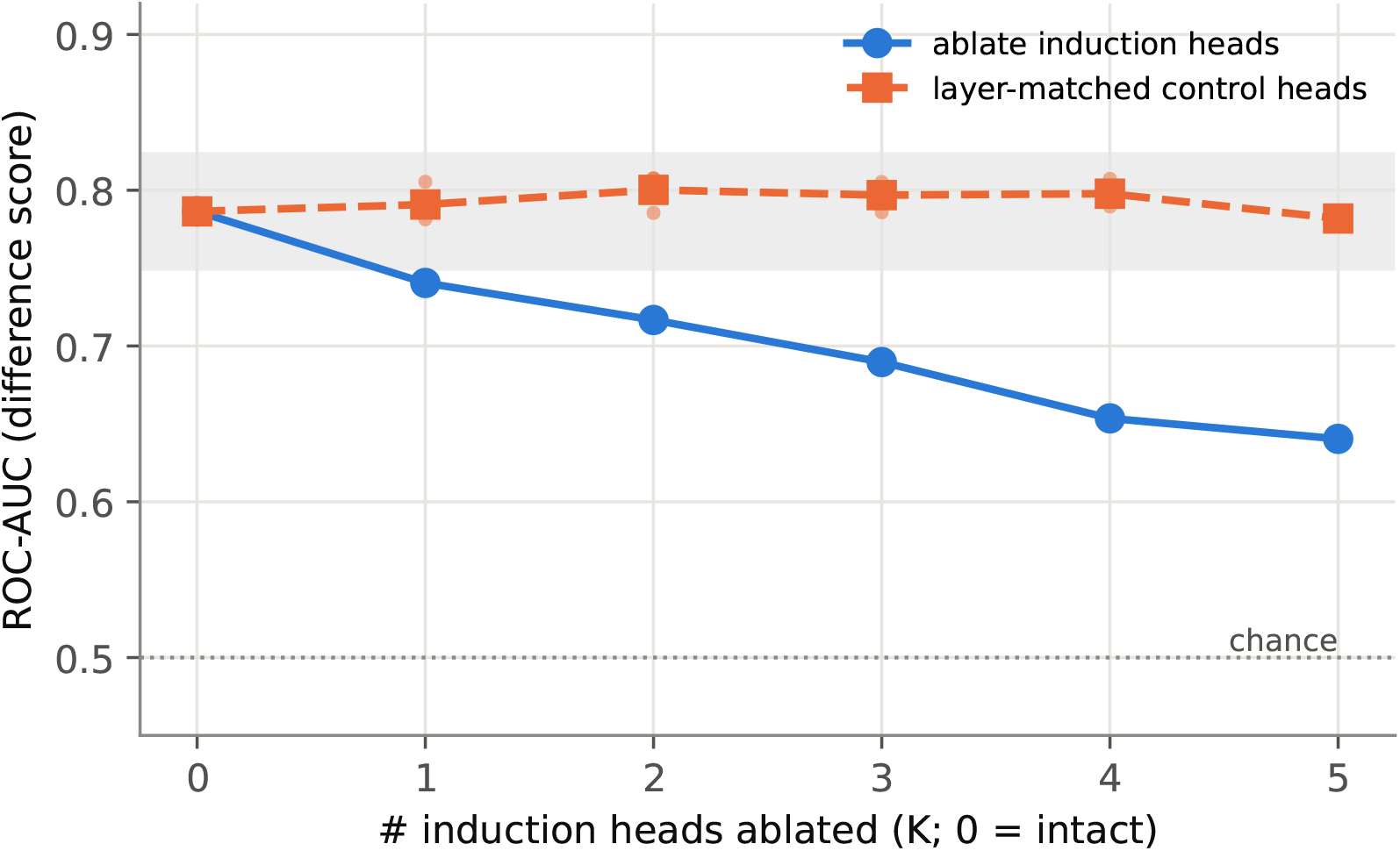
ProGen2-small binder screen after ablating *K* induction heads, using 16-example prompt. Grey area depicts noise band of the intact model.

**Figure 32:**
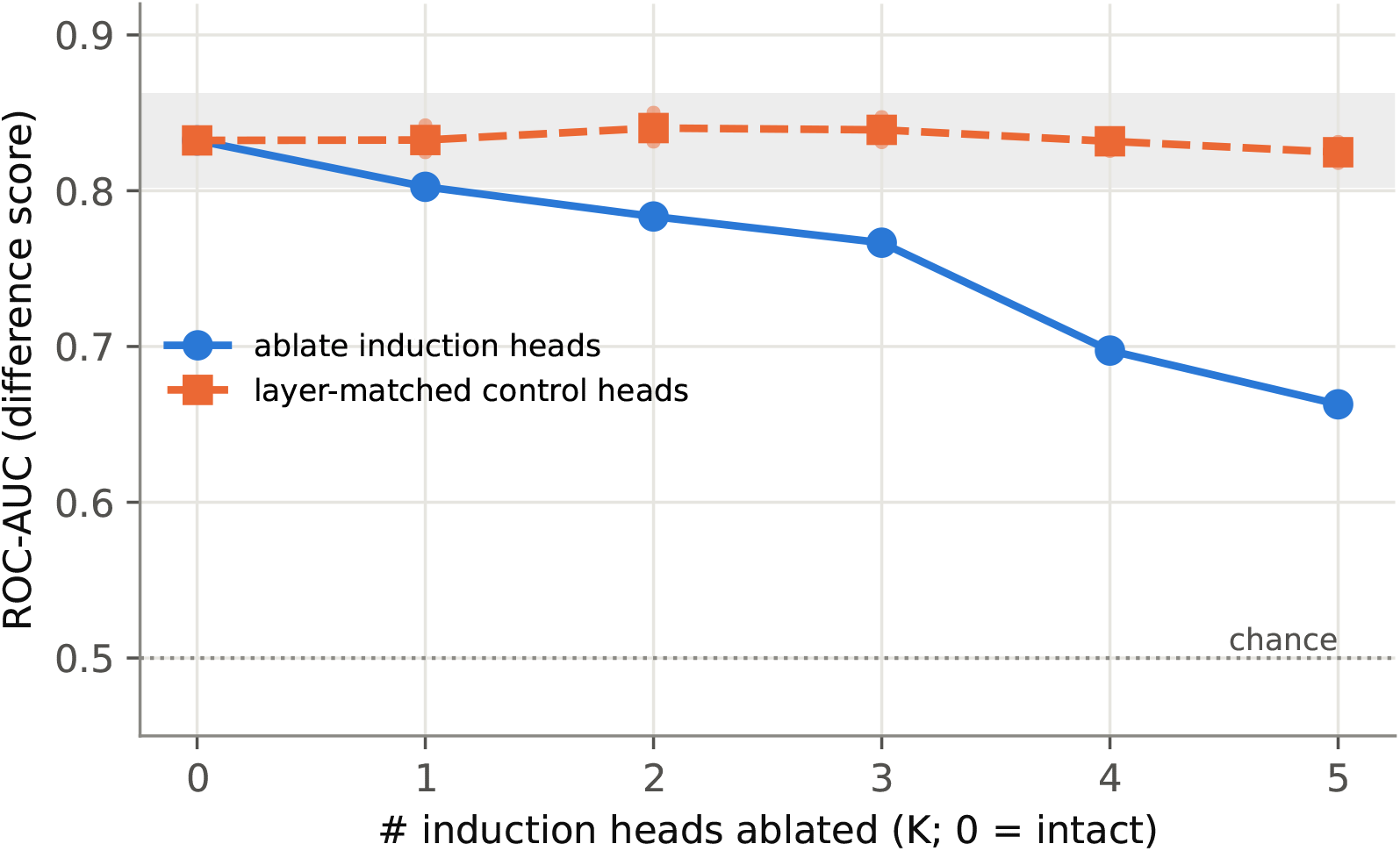
ProGen2-small binder screen after ablating *K* induction heads, using 32-example prompt. Grey area depicts noise band of the intact model

### E.2 ProGen2-base (765M)

**Figure 33:**
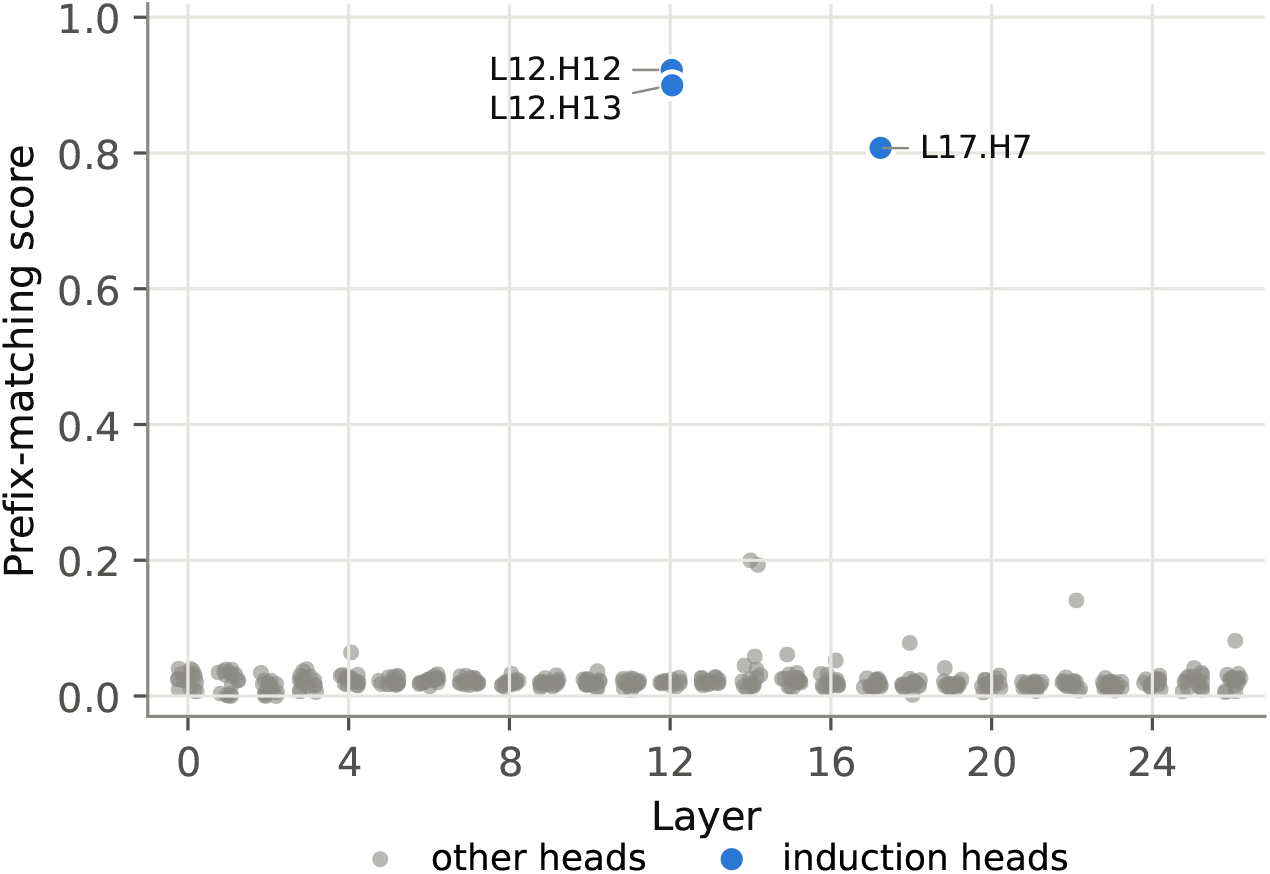
Prefix matching scatter in layers surrounding induction heads of ProGen2-base.

**Figure 34:**
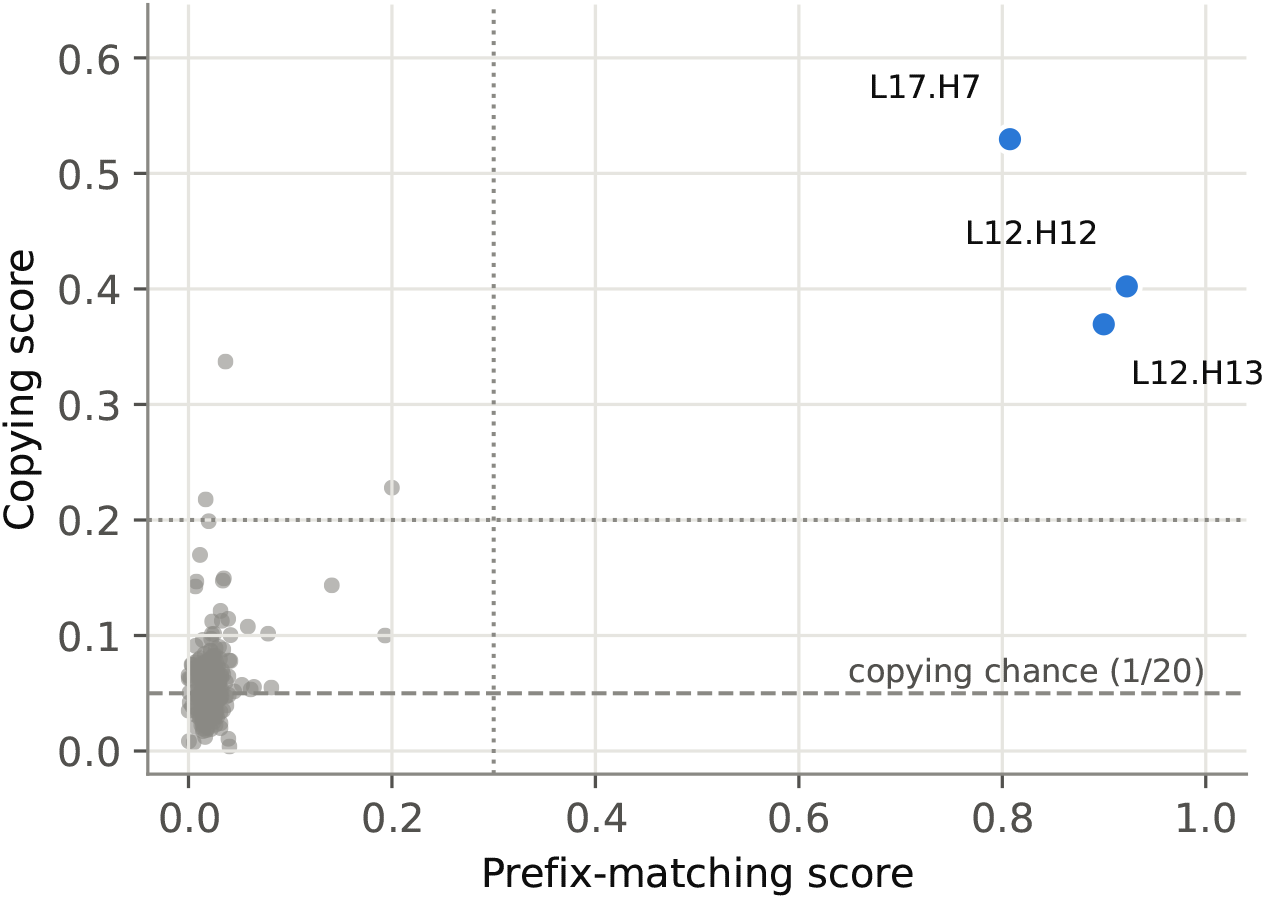
Prefix matching and copying scores of all heads in ProGen2-base.

**Figure 35:**
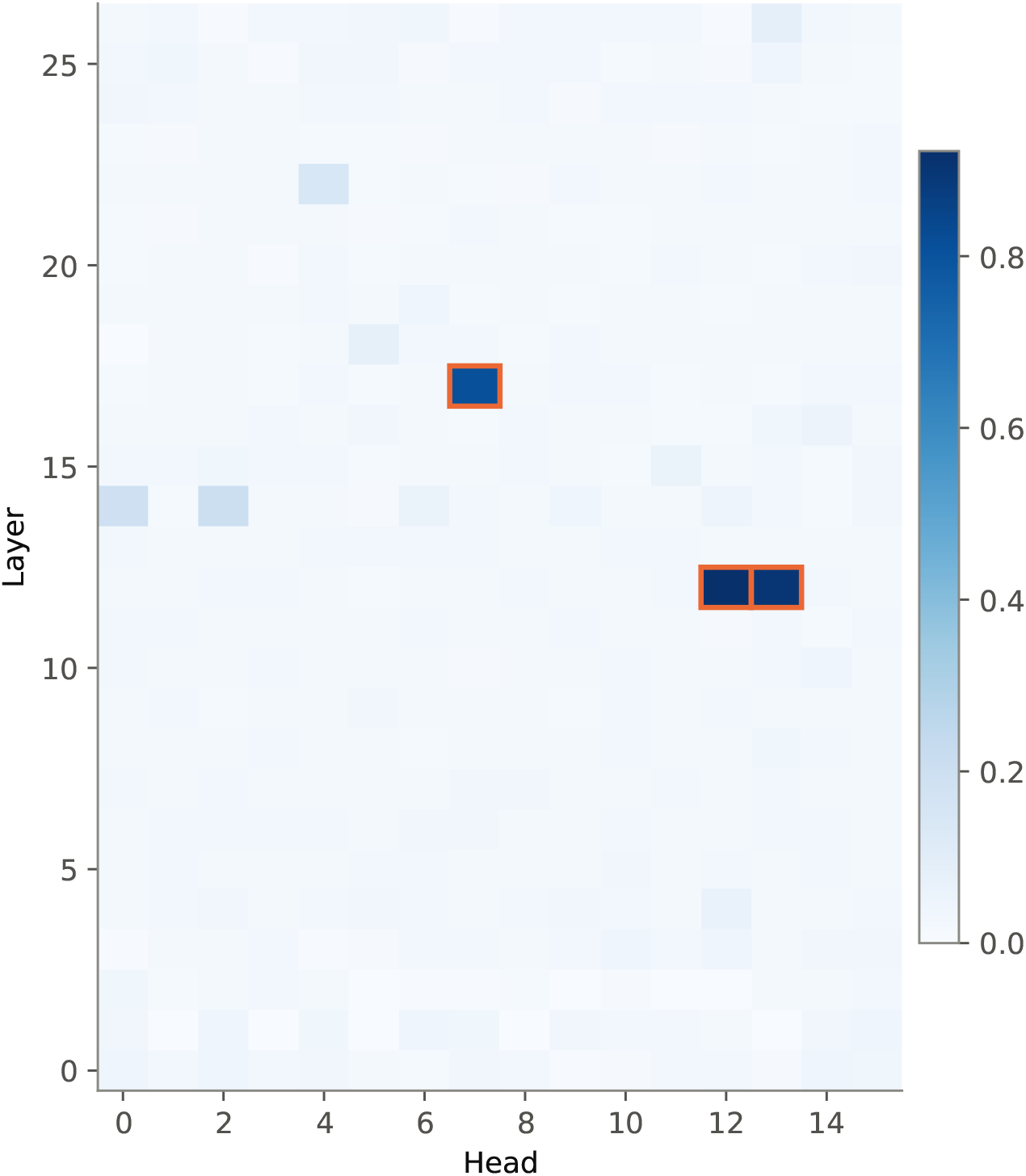
Prefix matching heatmap of induction-head and nearby layers in ProGen2-base. Induction heads are highlighted with orange border.

**Figure 36:**
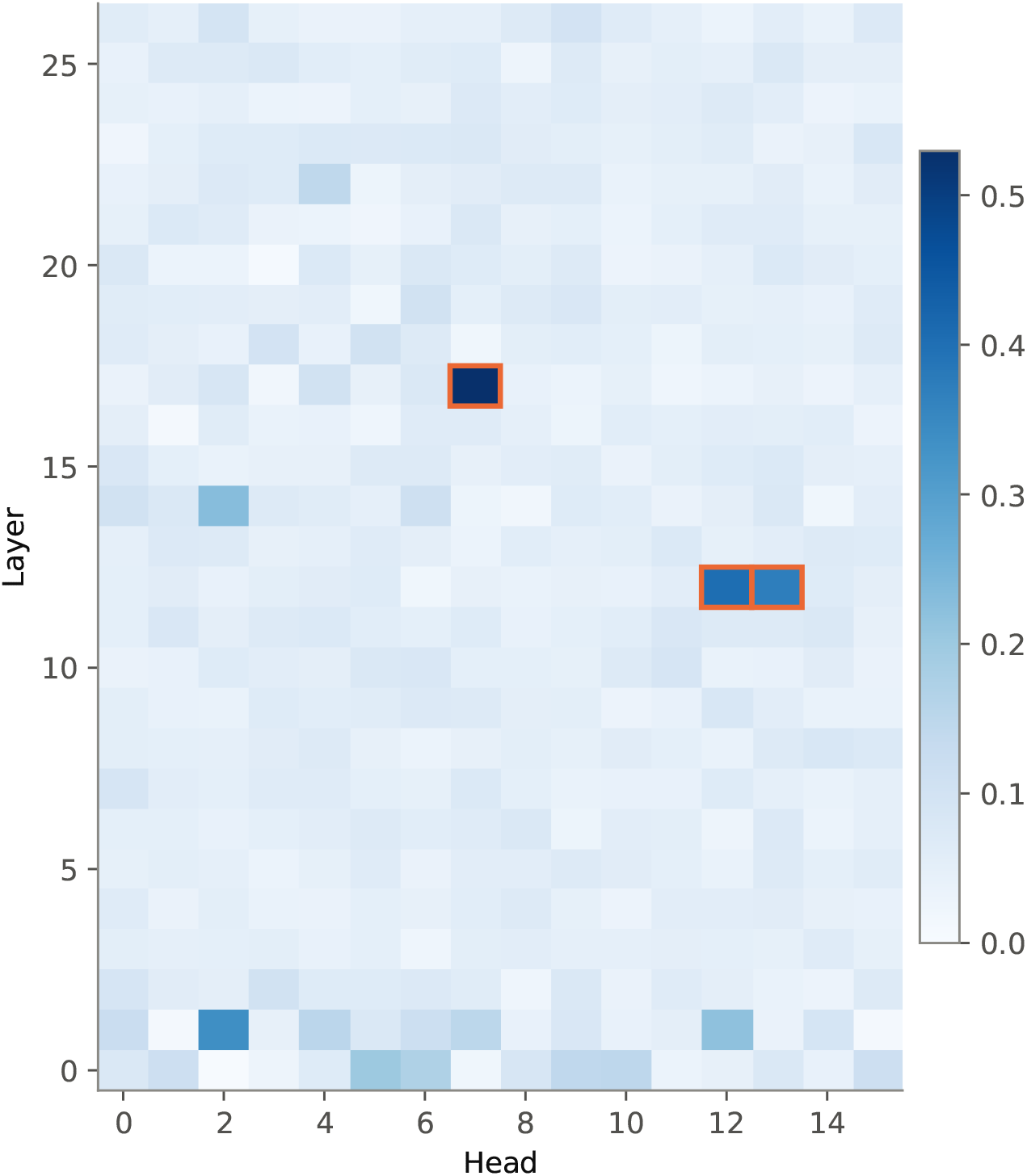
Copying heatmap of induction-head and nearby layers in ProGen2-base.

**Figure 37:**
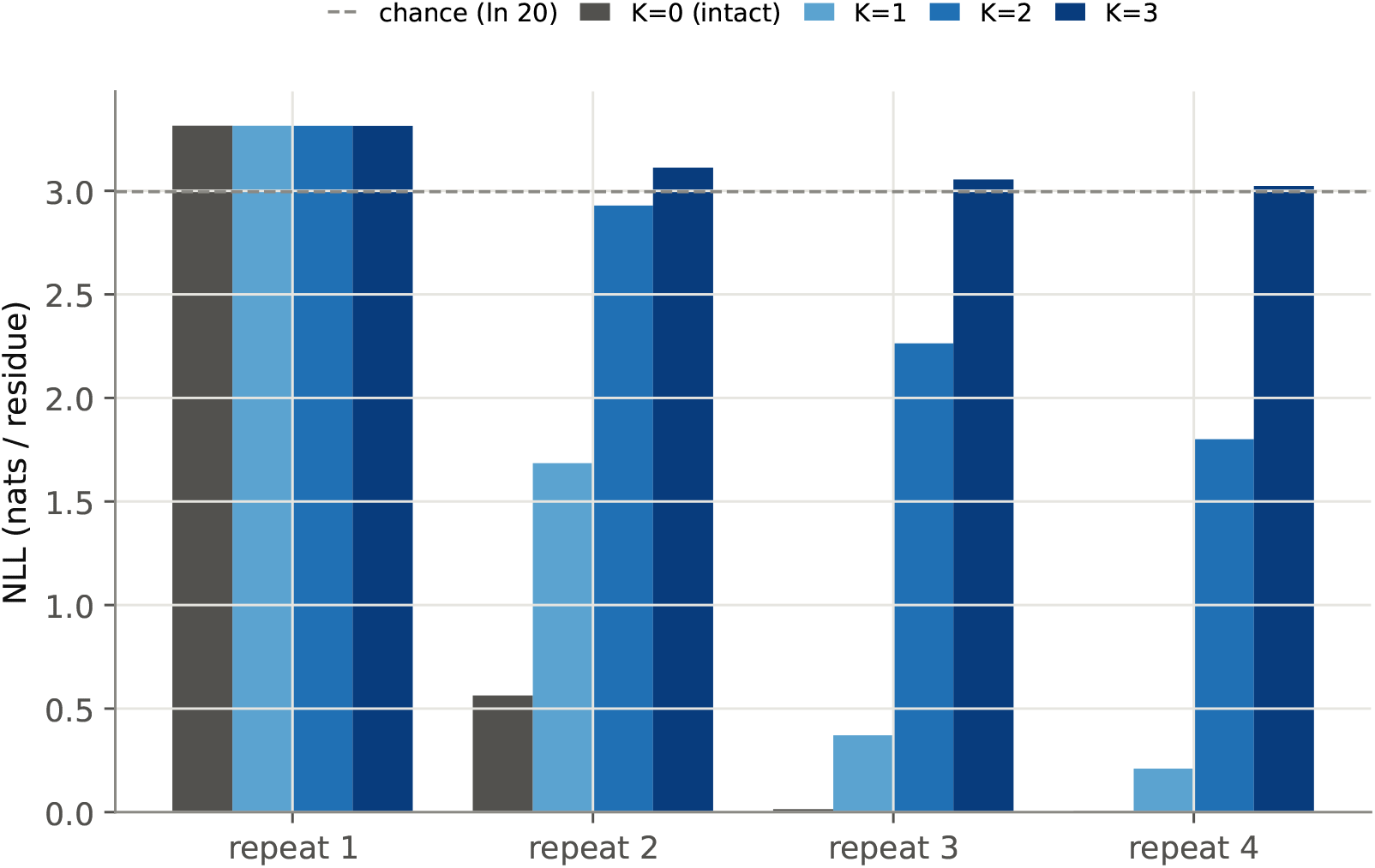
NLL loss-per-residue across all random sequence repeats during ProGen2-base inference, with *K* induction head(s) zero-ed out.

**Figure 38:**
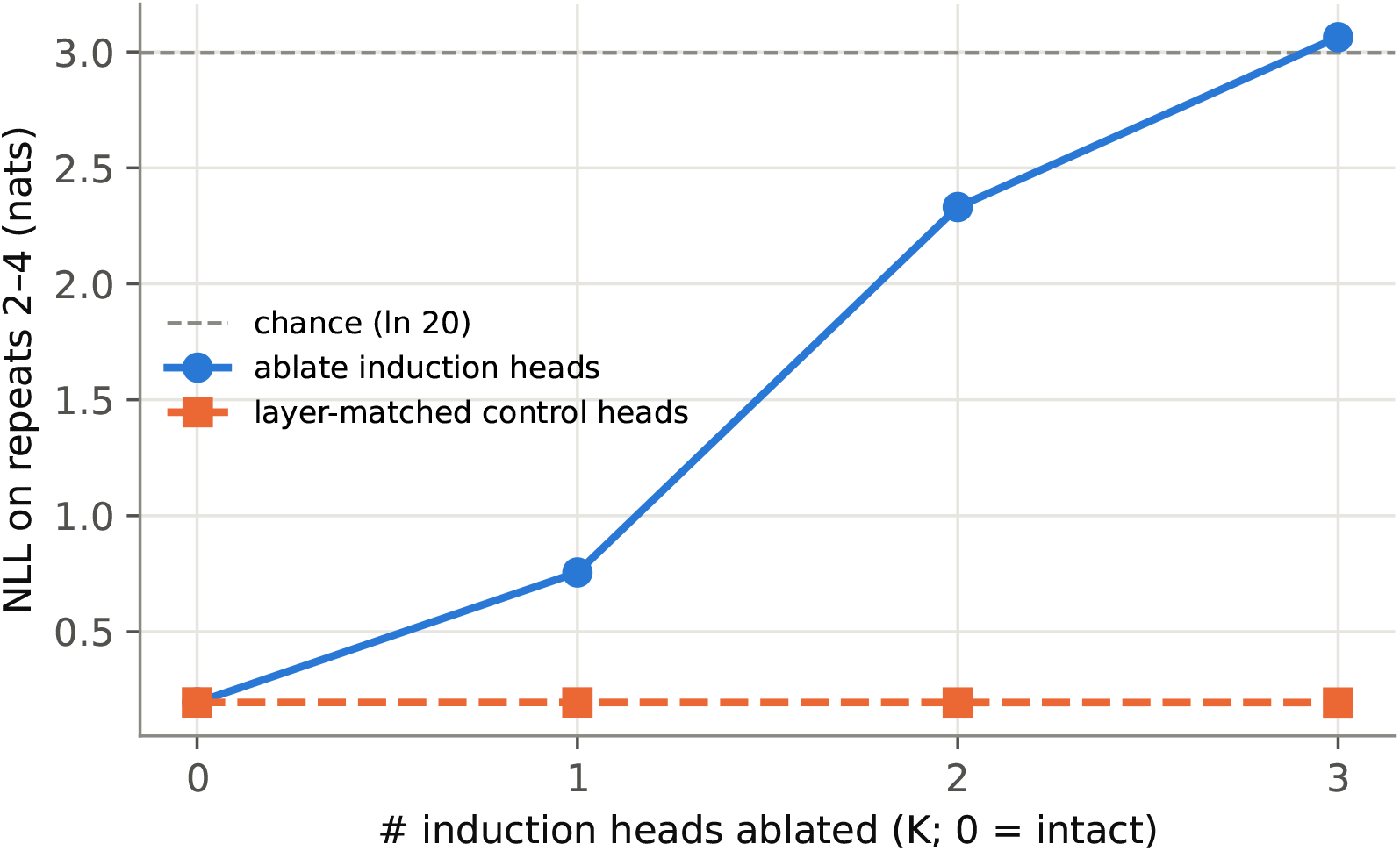
NLL loss-per-residue across random sequences repeats 2-4 between induction head ablation and control (ablating the same number of random, non-induction heads in the same induction layers) during ProGen2-base inference.

**Figure 39:**
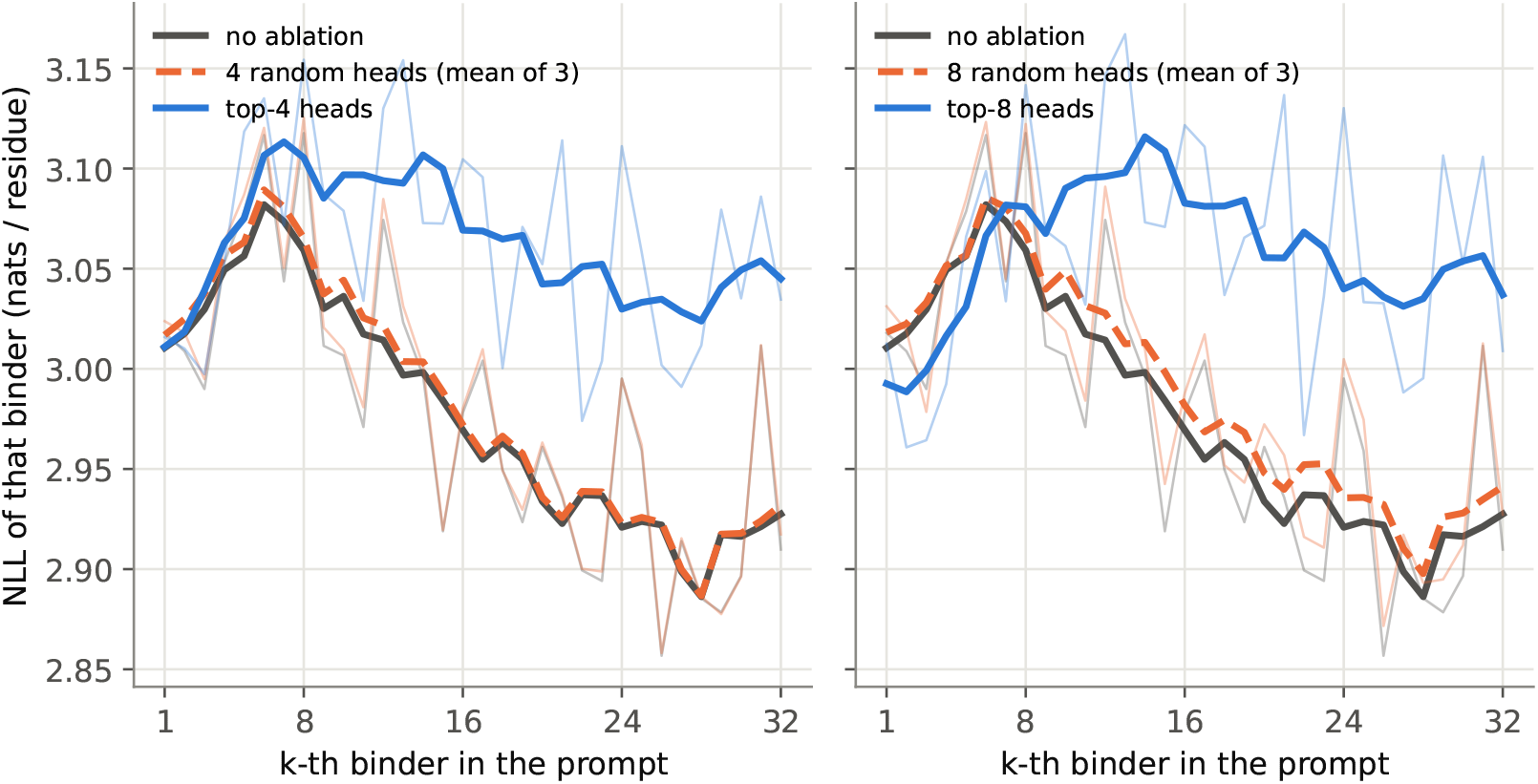
NLL loss-per-residue for each 15-mer example, averaged over 10 ProGen2-base inference using prompts constructed from 32 binder examples. Three parallel schemes are presented: no ablation, top-*K* heads and *K* random heads sampled from the same layers as top-*K* (averaged over 3 independent sampled set of heads). *\*Note*: NLL axis in this plot is positive (opposite to Fig. 6B); here, lower is better.

**Figure 40:**
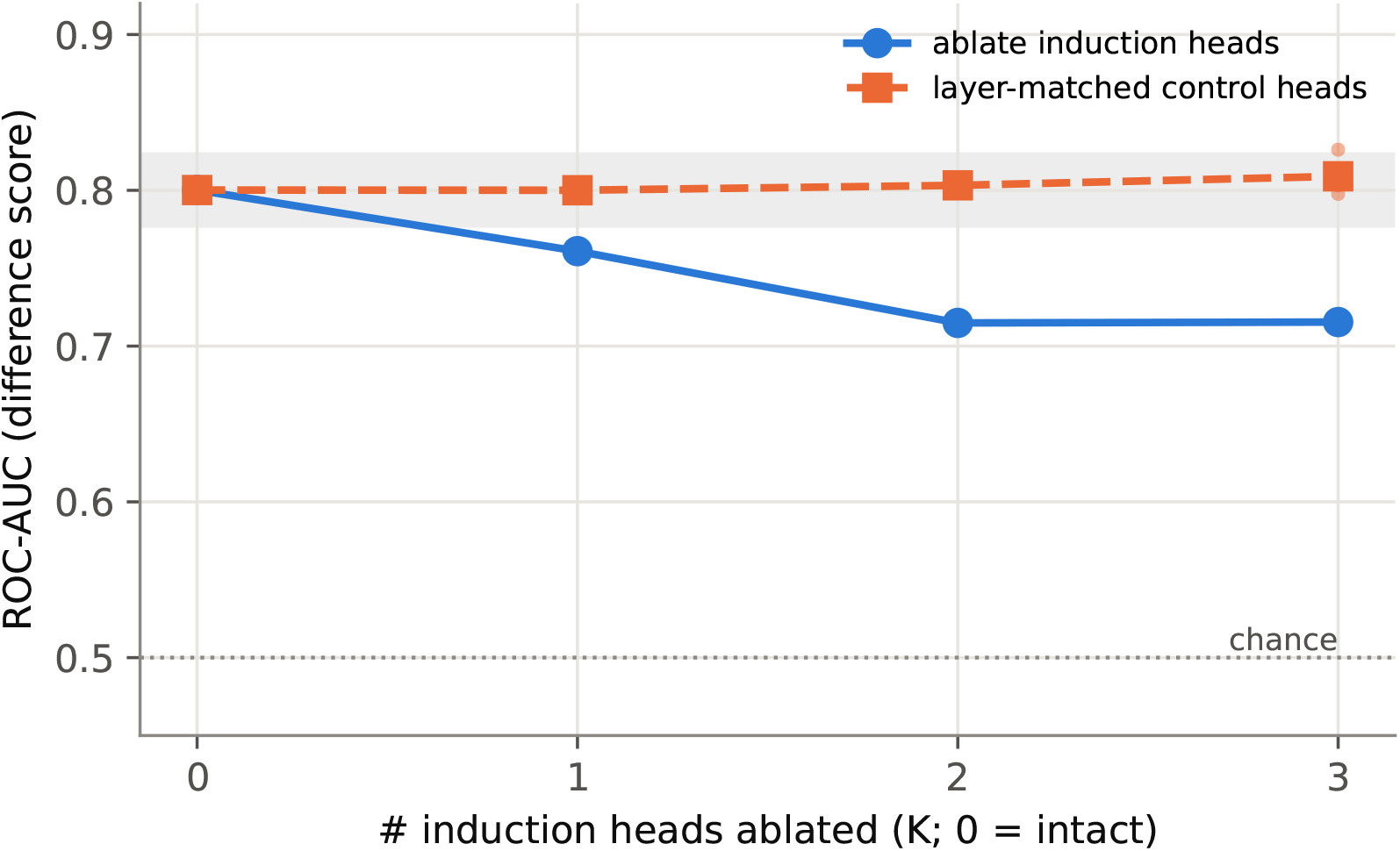
ProGen2-base binder screen after ablating *K* induction heads, using 16-example prompt. Grey area depicts noise band of the intact model

**Figure 41:**
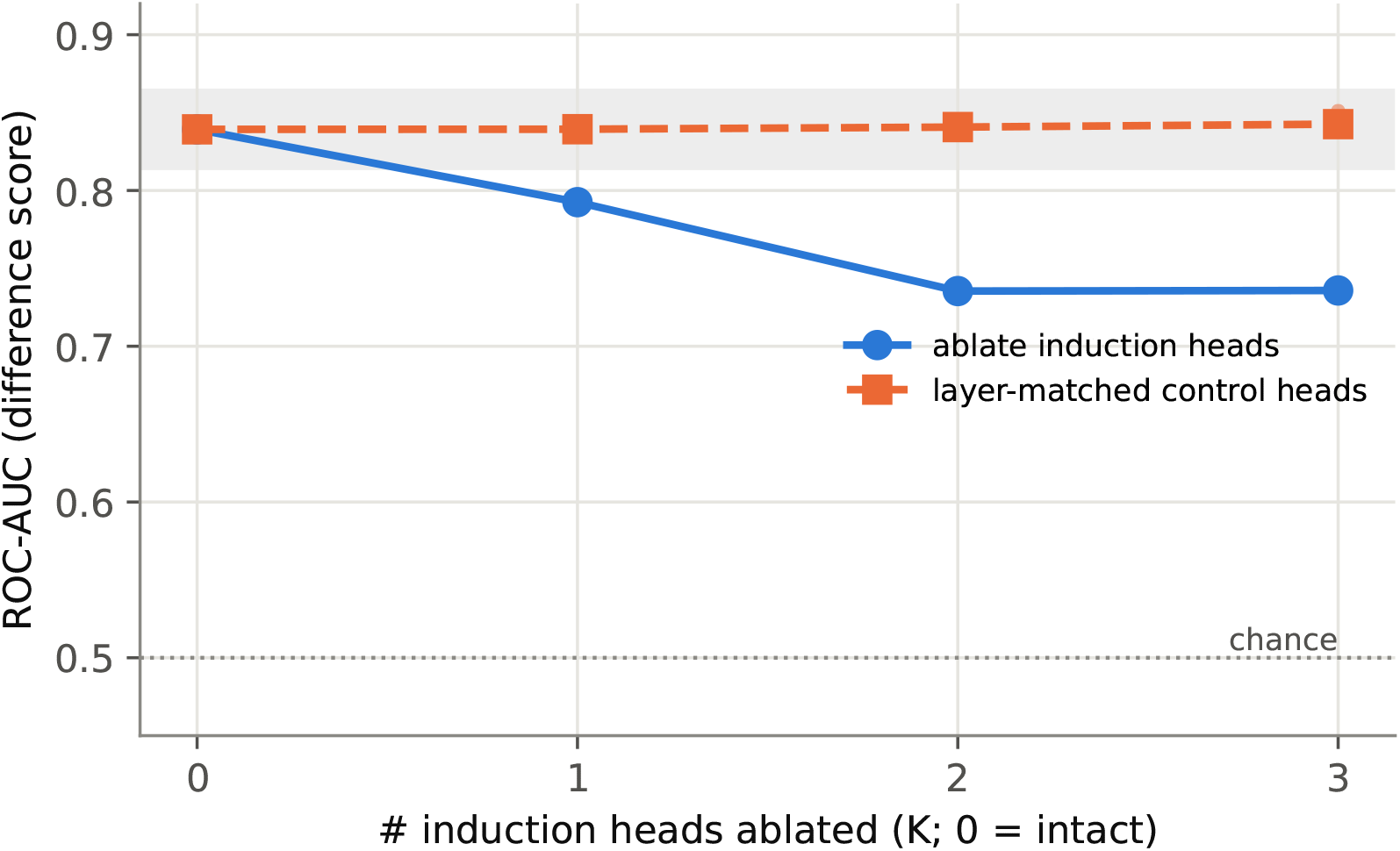
ProGen2-base binder screen after ablating *K* induction heads, using 32-example prompt. Grey area depicts noise band of the intact model.

### E.3 ProGen2-medium (765M)

**Figure 42:**
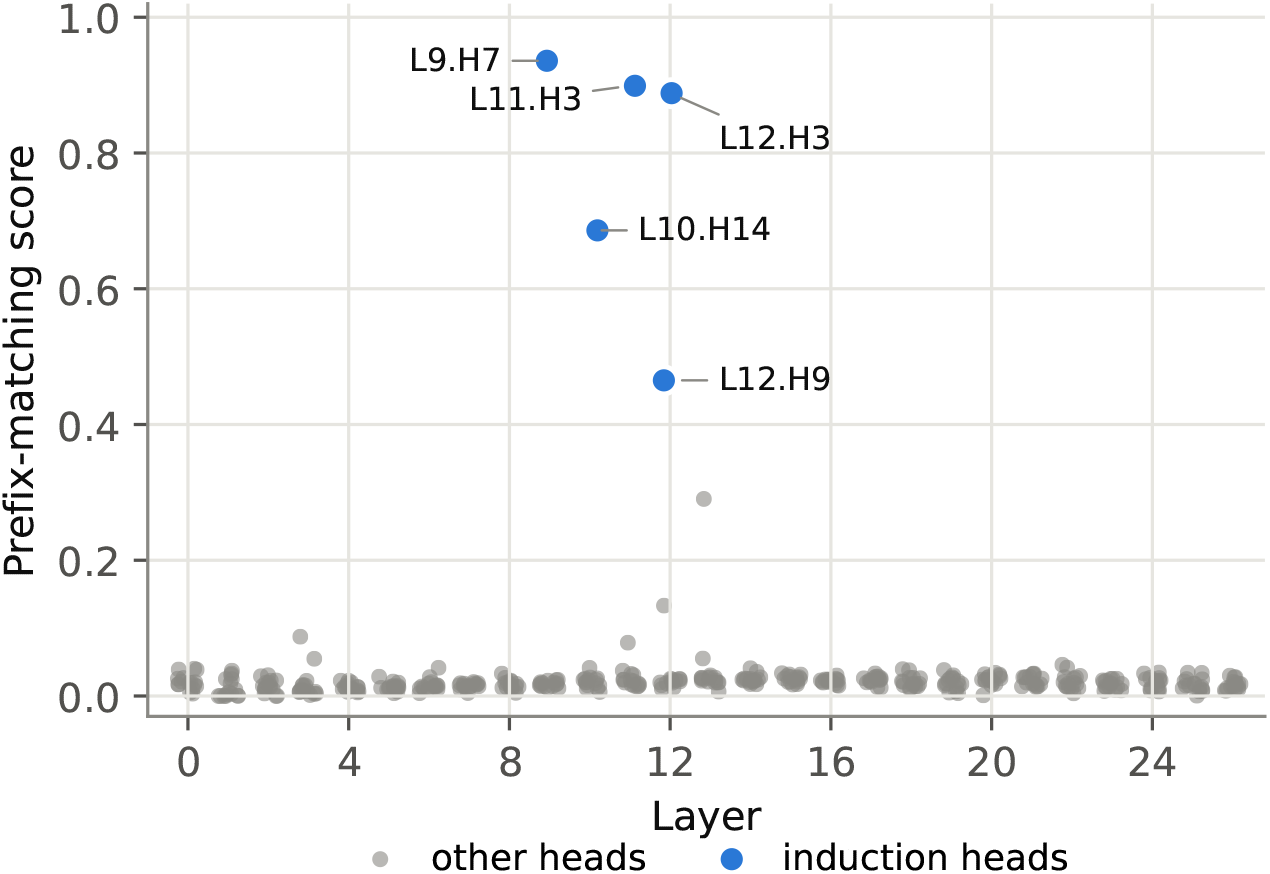
Prefix matching scatter in layers surrounding induction heads of ProGen2-medium.

**Figure 43:**
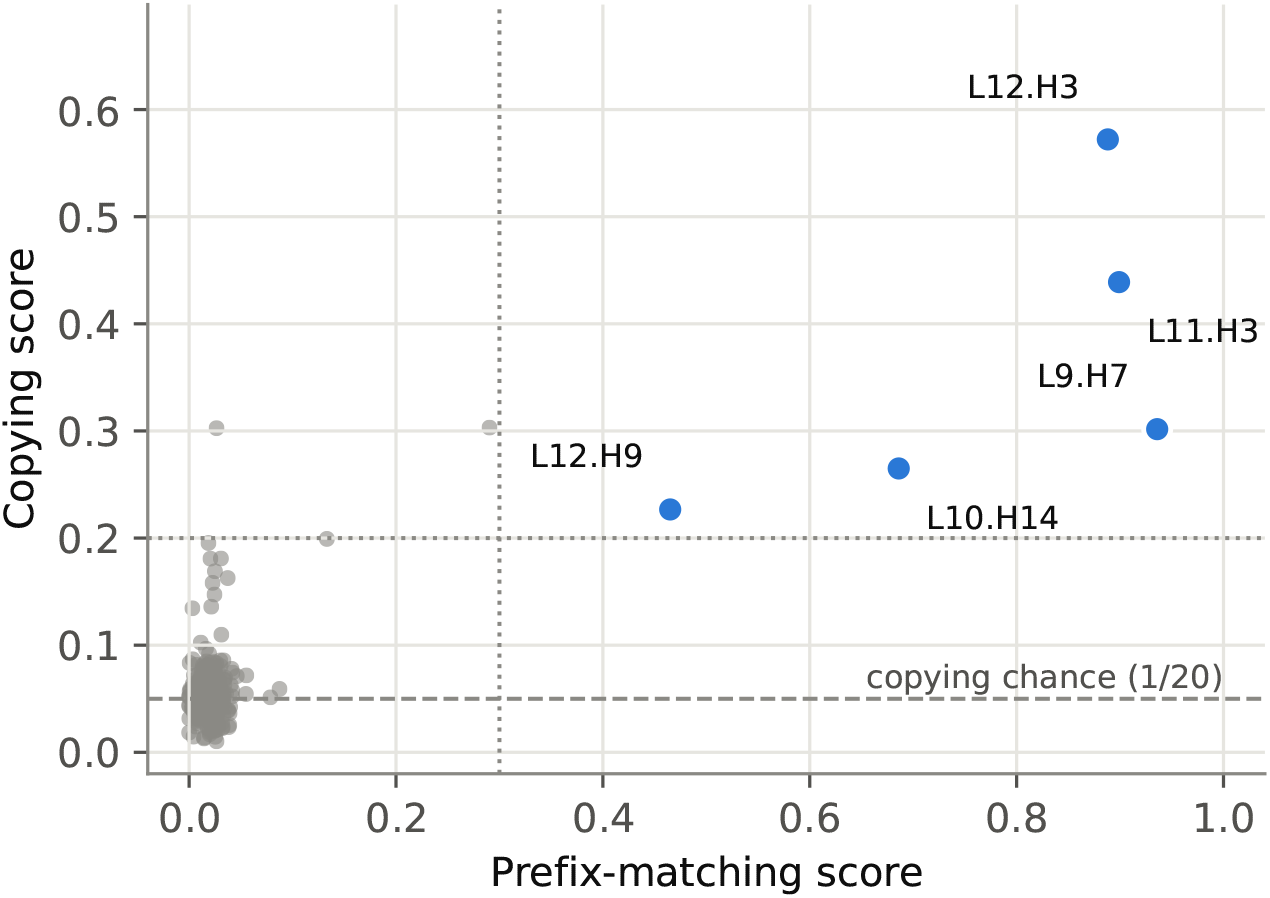
Prefix matching and copying scores of all heads in ProGen2-medium.

**Figure 44:**
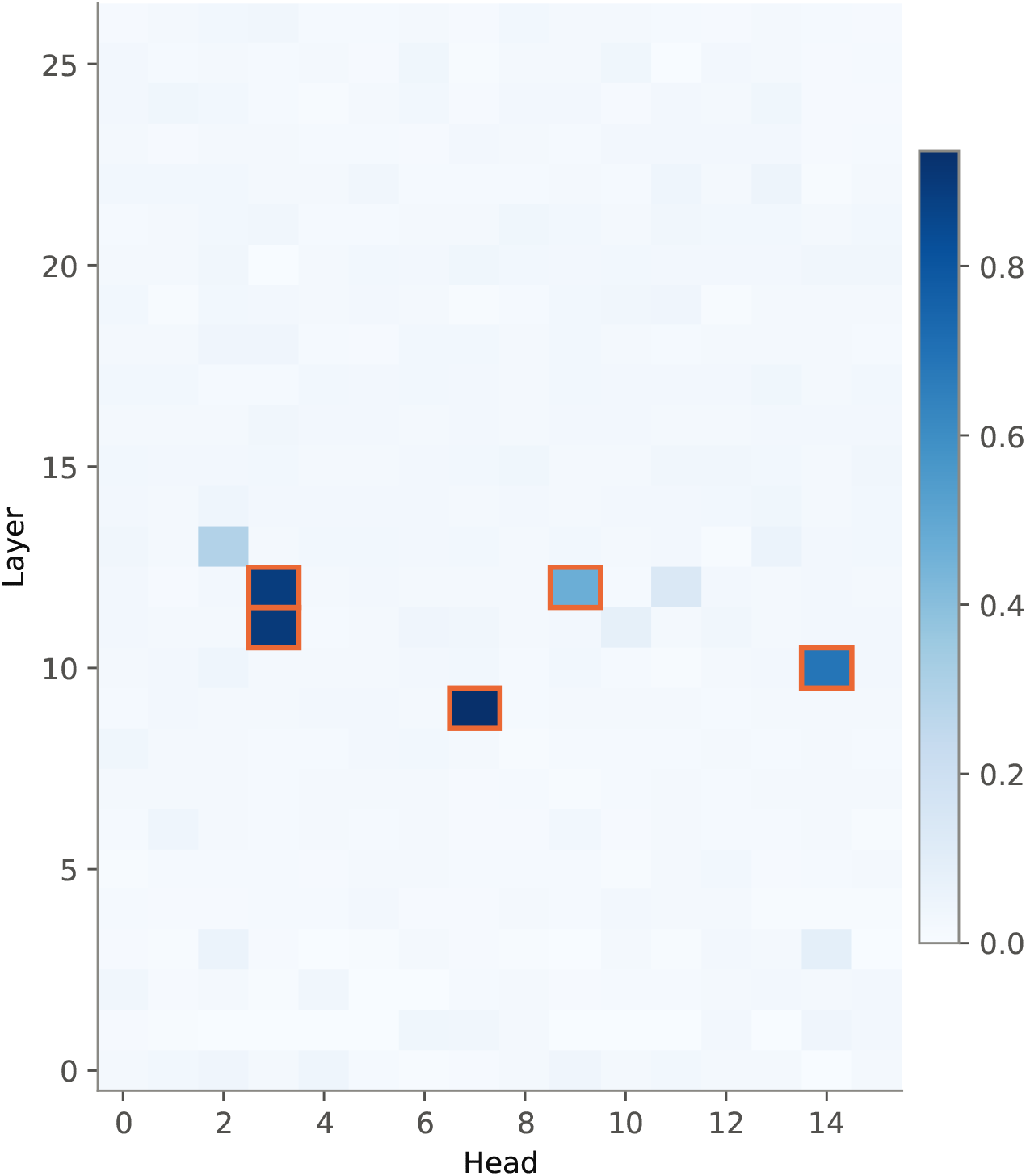
Prefix matching heatmap of induction-head and nearby layers in ProGen2-medium. Induction heads are highlighted with orange border.

**Figure 45:**
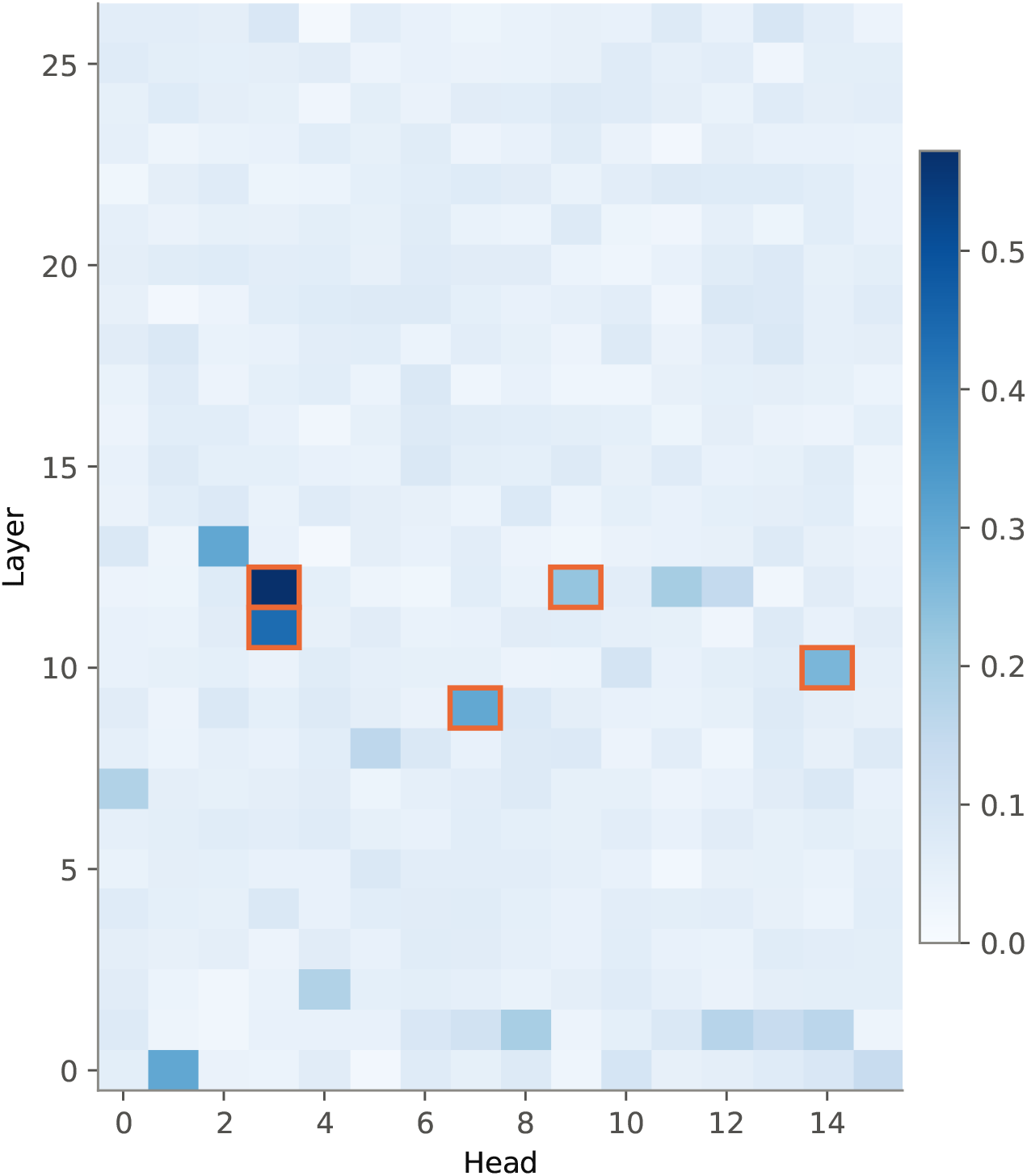
Copying heatmap of induction-head and nearby layers in ProGen2-medium.

**Figure 46:**
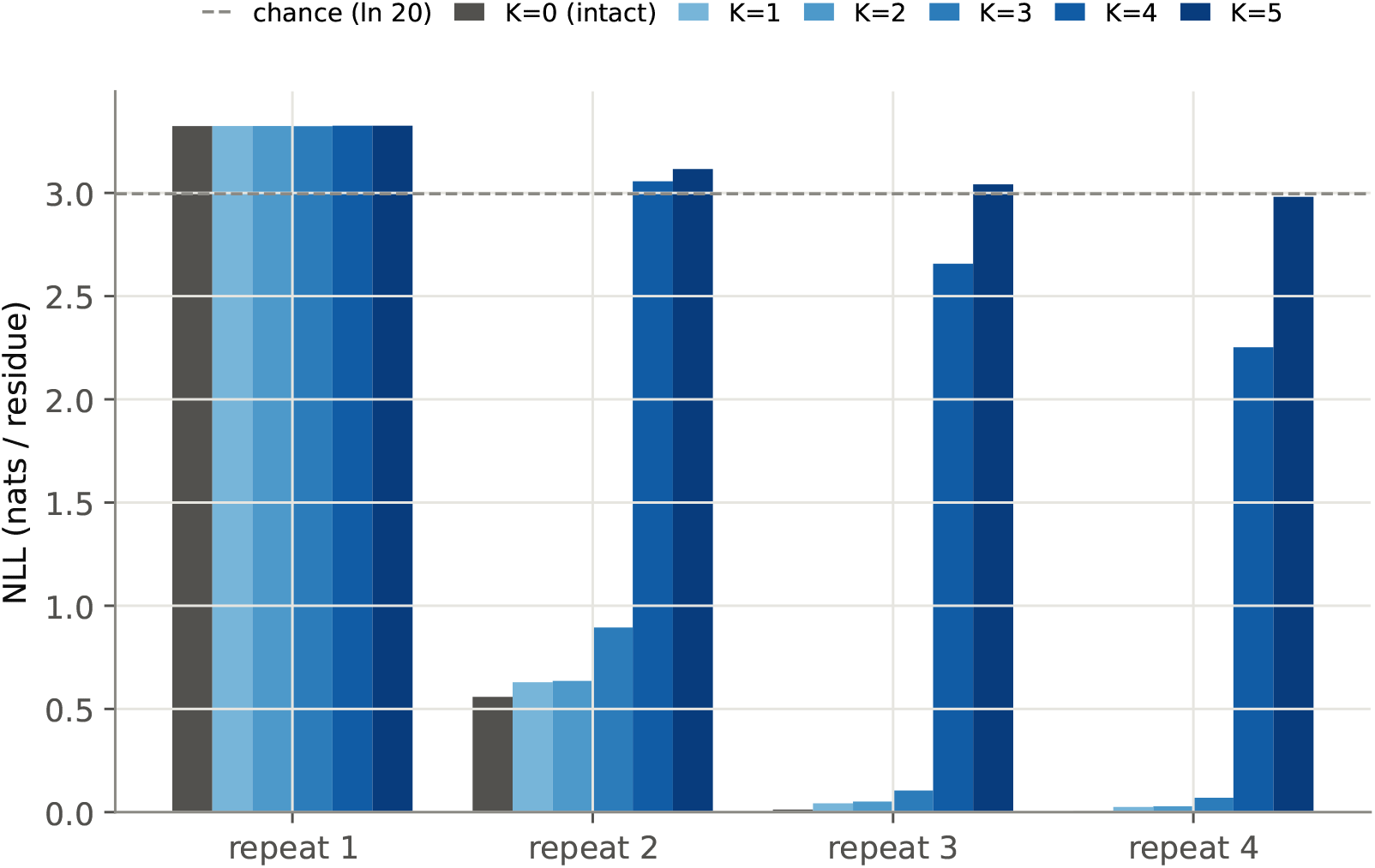
NLL loss-per-residue across all random sequence repeats during ProGen2-medium inference, with *K* induction head(s) zero-ed out.

**Figure 47:**
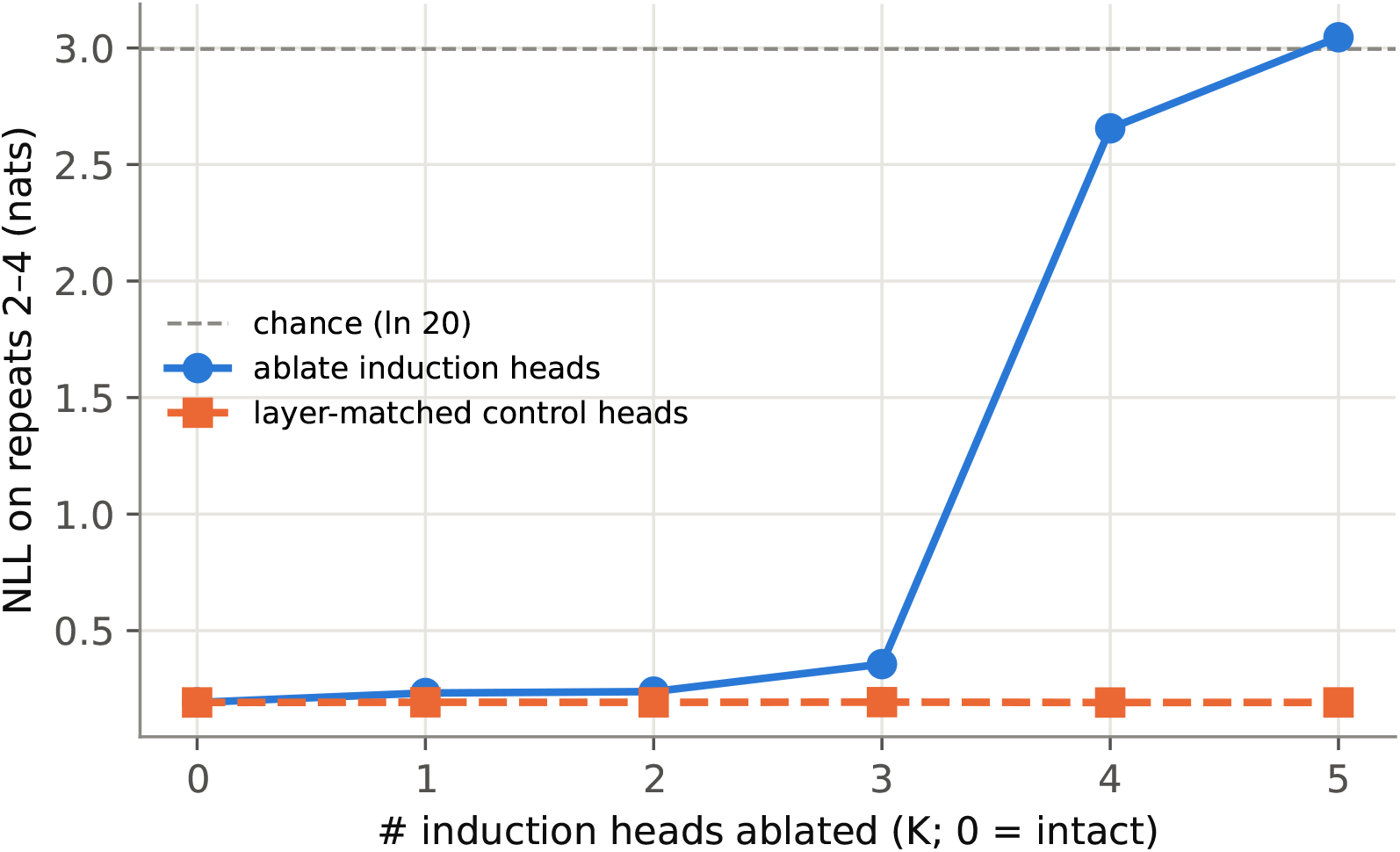
NLL loss-per-residue across random sequences repeats 2-4 between induction head ablation and control (ablating the same number of random, non-induction heads in the same induction layers) during ProGen2-medium inference.

**Figure 48:**
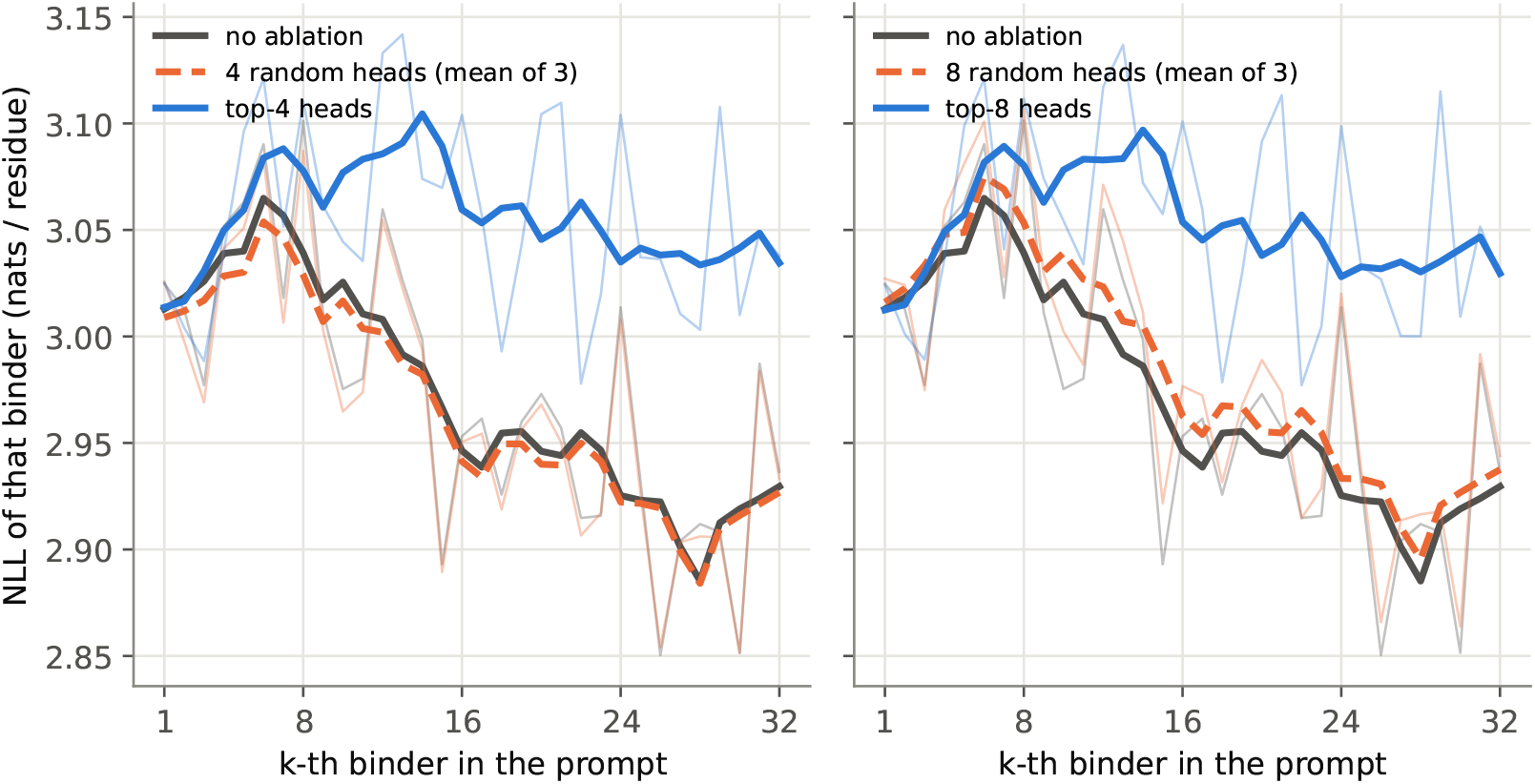
NLL loss-per-residue for each 15-mer example, averaged over 10 ProGen2-medium inference using prompts constructed from 32 binder examples. Three parallel schemes are presented: no ablation, top-*K* heads and *K* random heads sampled from the same layers as top-*K* (averaged over 3 independent sampled set of heads). *\*Note*: NLL axis in this plot is positive (opposite to Fig. 6B); here, lower is better.

**Figure 49:**
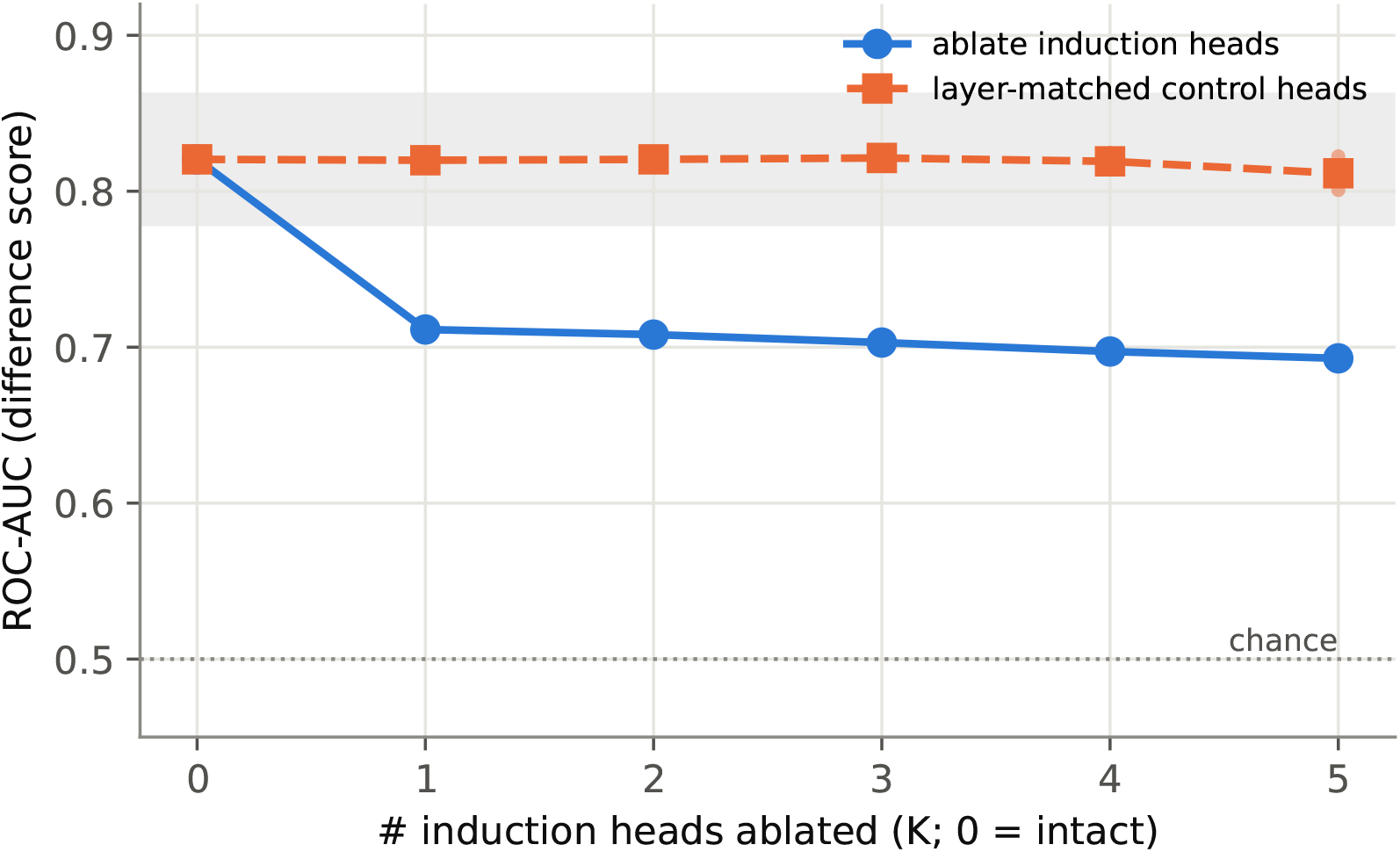
ProGen2-medium binder screen after ablating *K* induction heads, using 16-example prompt. Grey area depicts noise band of the intact model.

**Figure 50:**
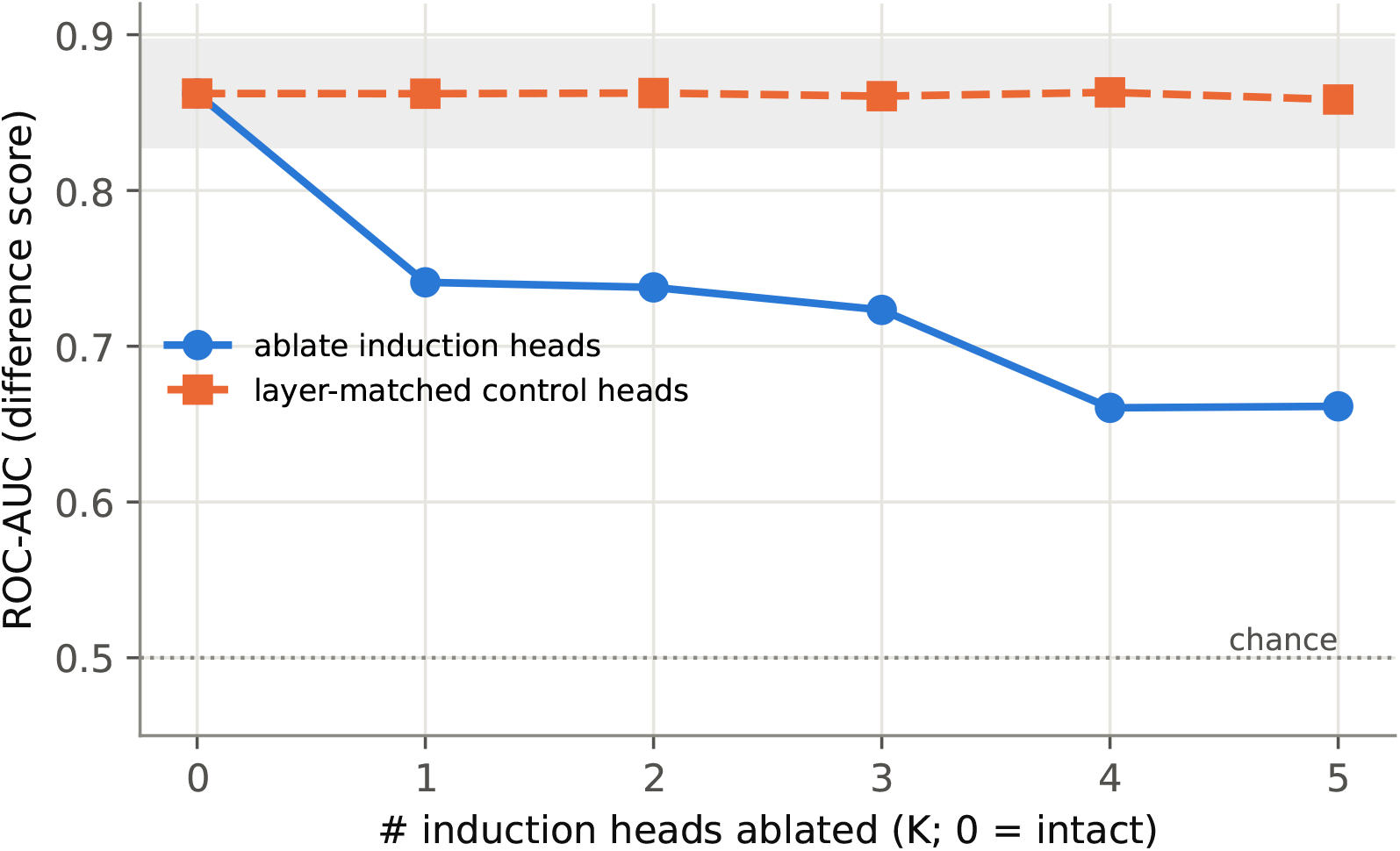
ProGen2-medium binder screen after ablating *K* induction heads, using 32-example prompt. Grey area depicts noise band of the intact model.

### E.4 ProGen2-large (2.78B)

**Figure 51:**
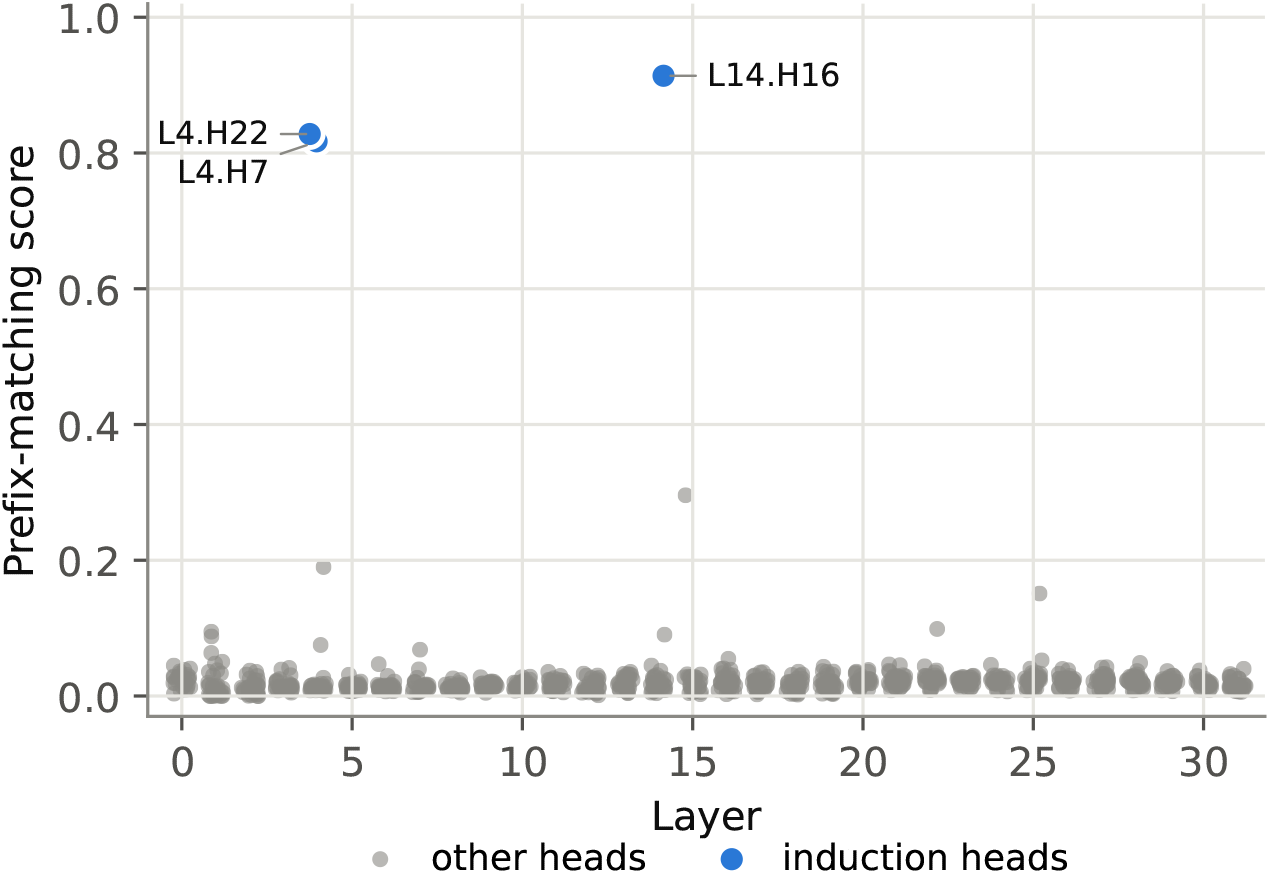
Prefix matching scatter in layers surrounding induction heads of ProGen2-large.

**Figure 52:**
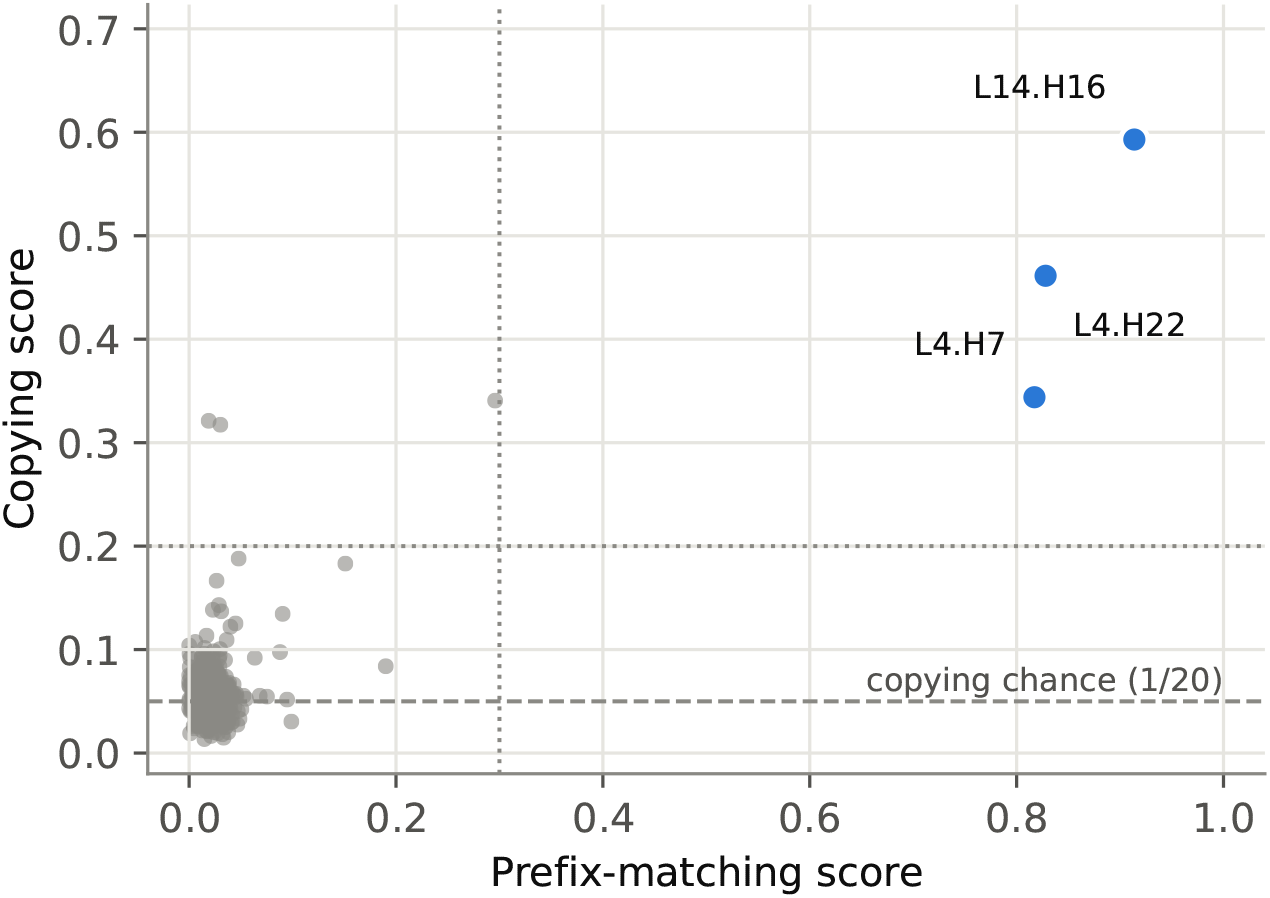
Prefix matching and copying scores of all heads in ProGen2-large.

**Figure 53:**
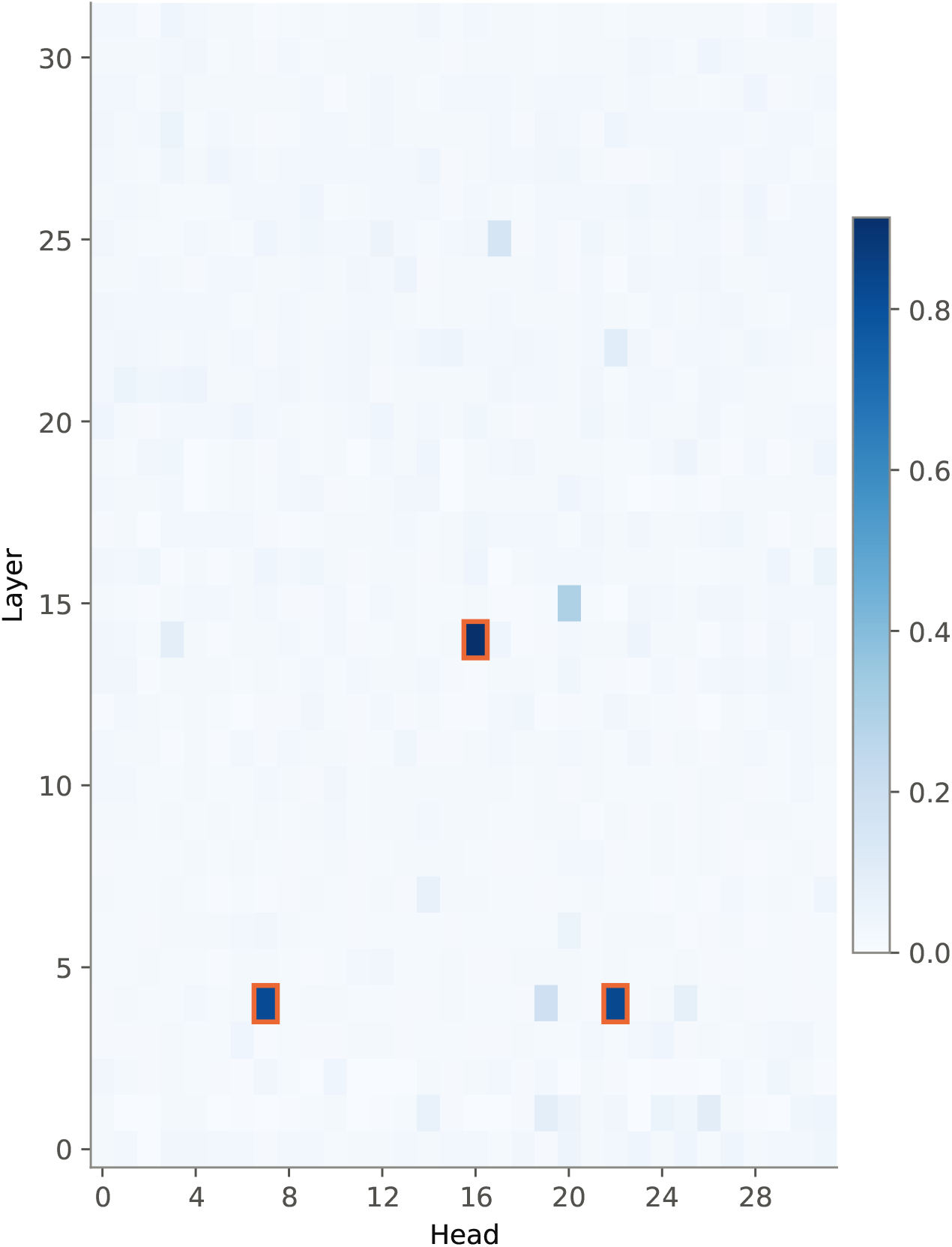
Prefix matching heatmap of induction-head and nearby layers in ProGen2-large. Induction heads are highlighted with orange border.

**Figure 54:**
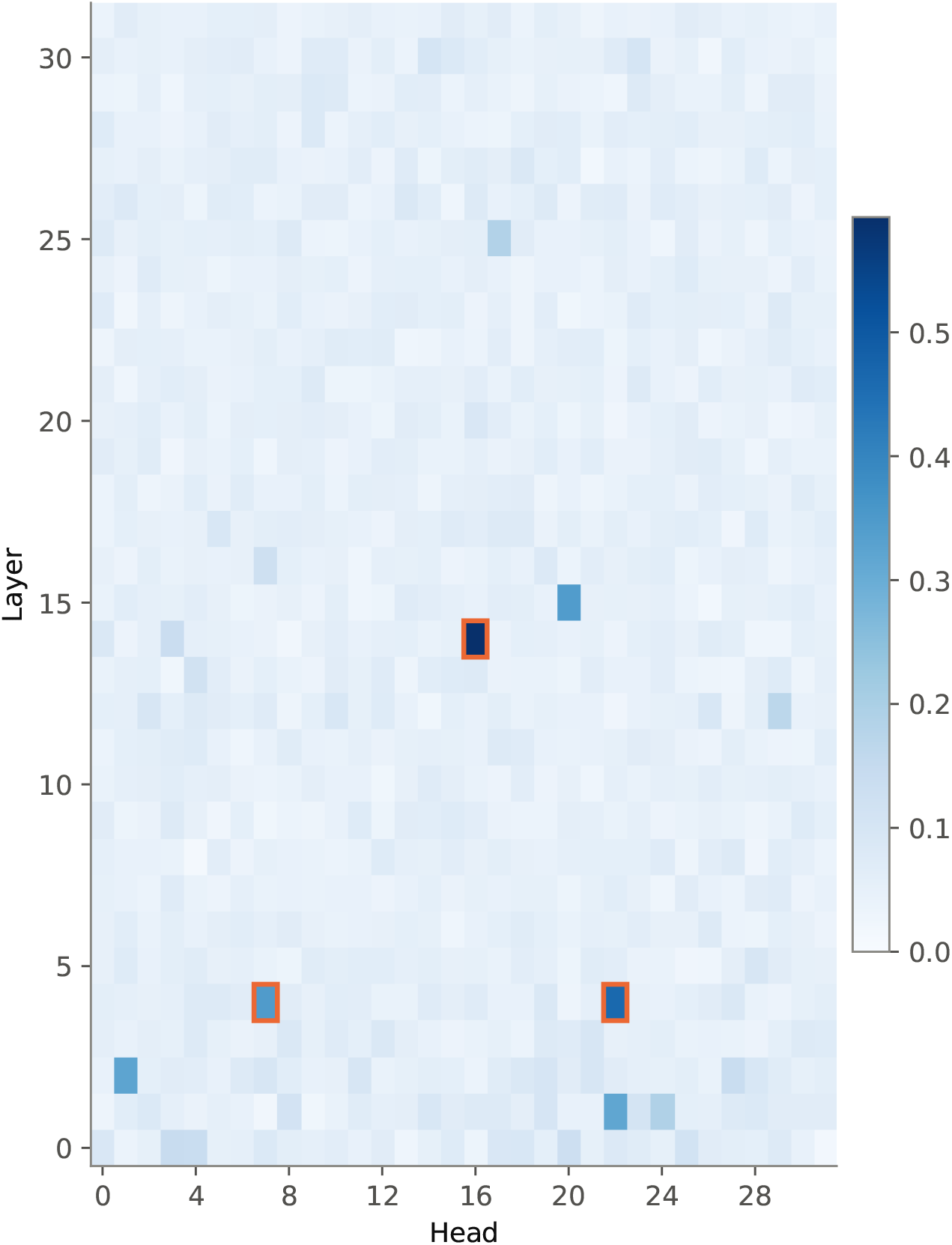
Copying heatmap of induction-head and nearby layers in ProGen2-large.

**Figure 55:**
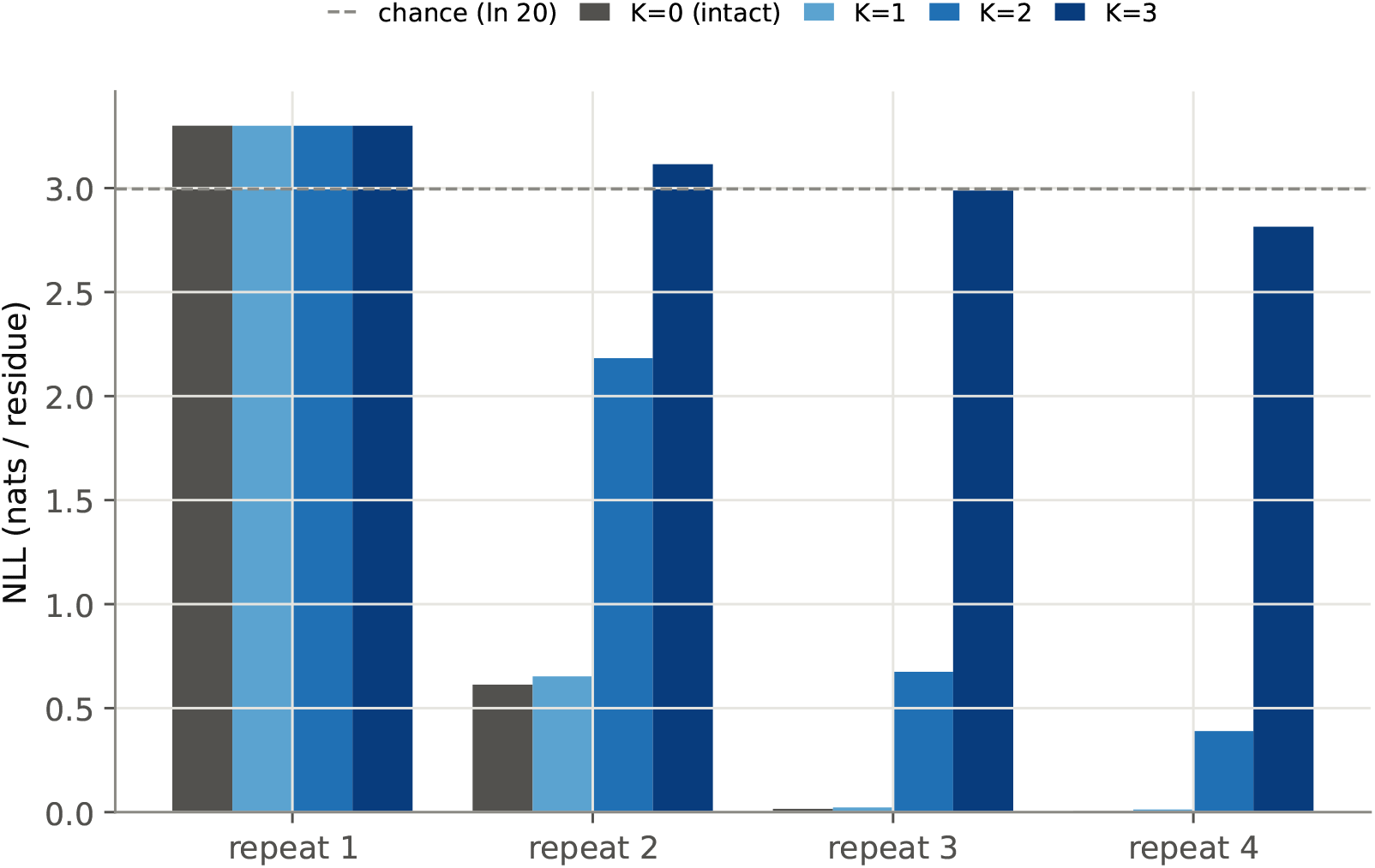
NLL loss-per-residue across all random sequence repeats during ProGen2-large inference, with *K* induction head(s) zero-ed out.

**Figure 56:**
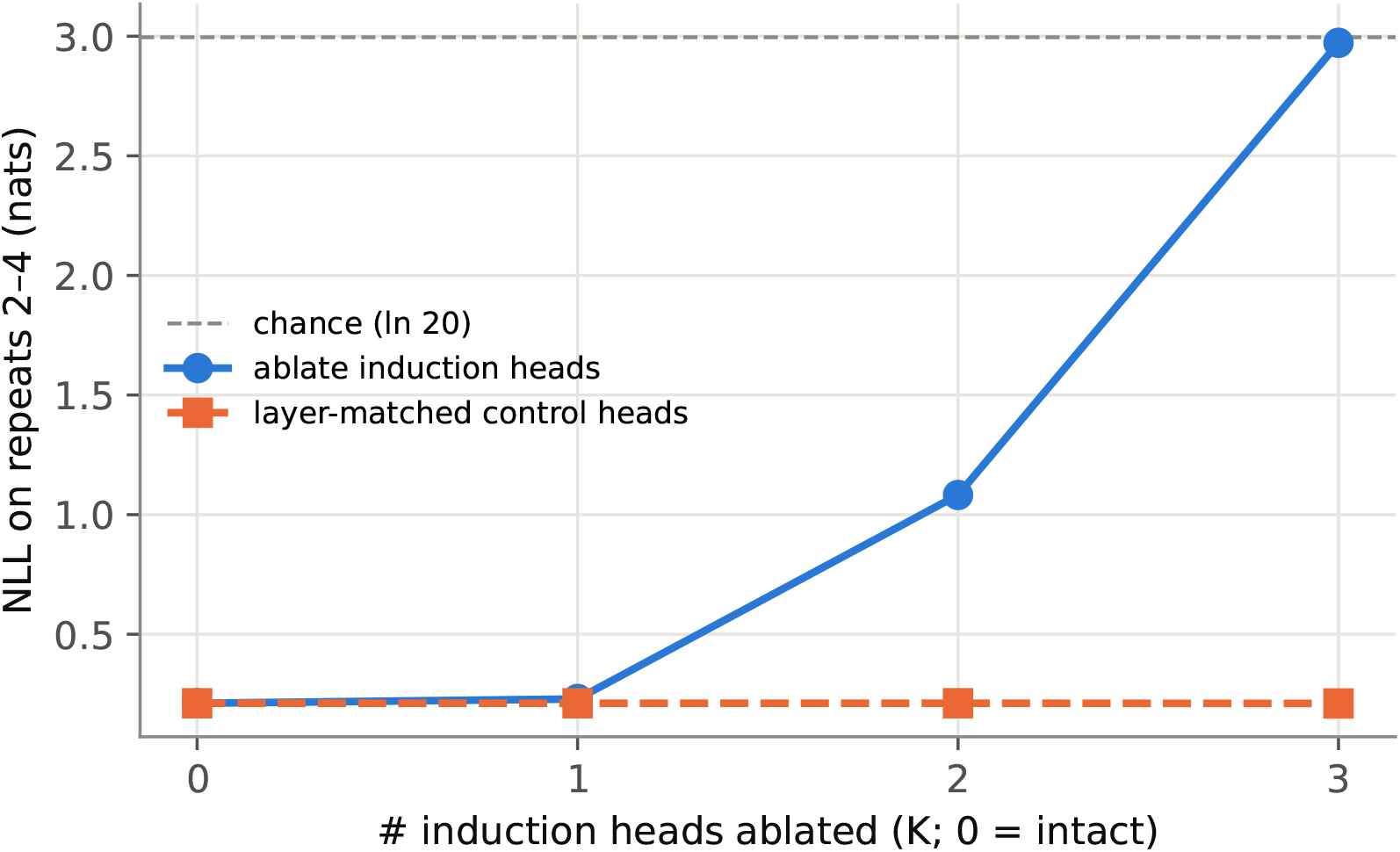
NLL loss-per-residue across random sequences repeats 2-4 between induction head ablation and control (ablating the same number of random, non-induction heads in the same induction layers) during ProGen2-large inference.

**Figure 57:**
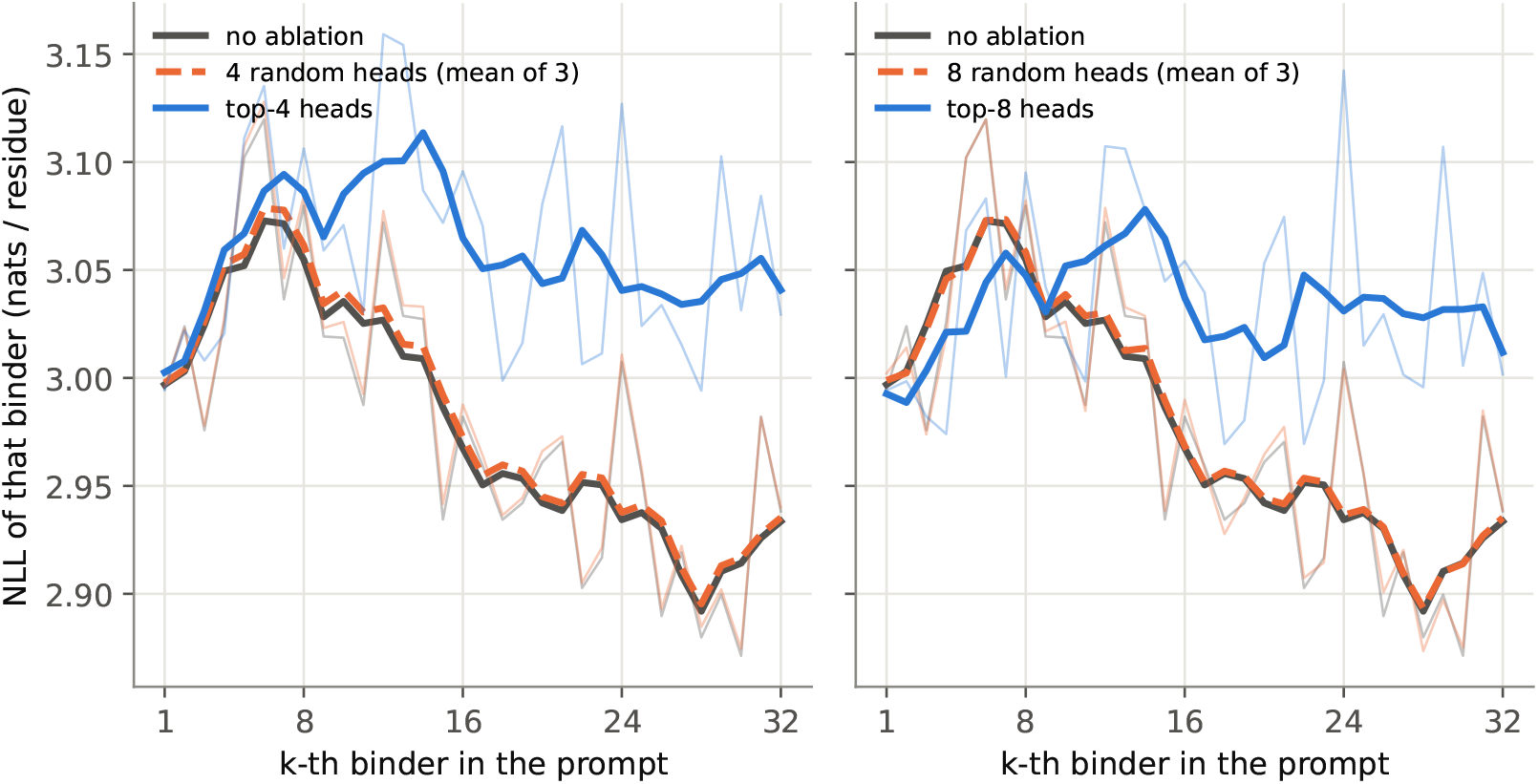
NLL loss-per-residue for each 15-mer example, averaged over 10 ProGen2-large inference using prompts constructed from 32 binder examples. Three parallel schemes are presented: no ablation, top-*K* heads and *K* random heads sampled from the same layers as top-*K* (averaged over 3 independent sampled set of heads). *\*Note*: NLL axis in this plot is positive (opposite to Fig. 6B); here, lower is better.

**Figure 58:**
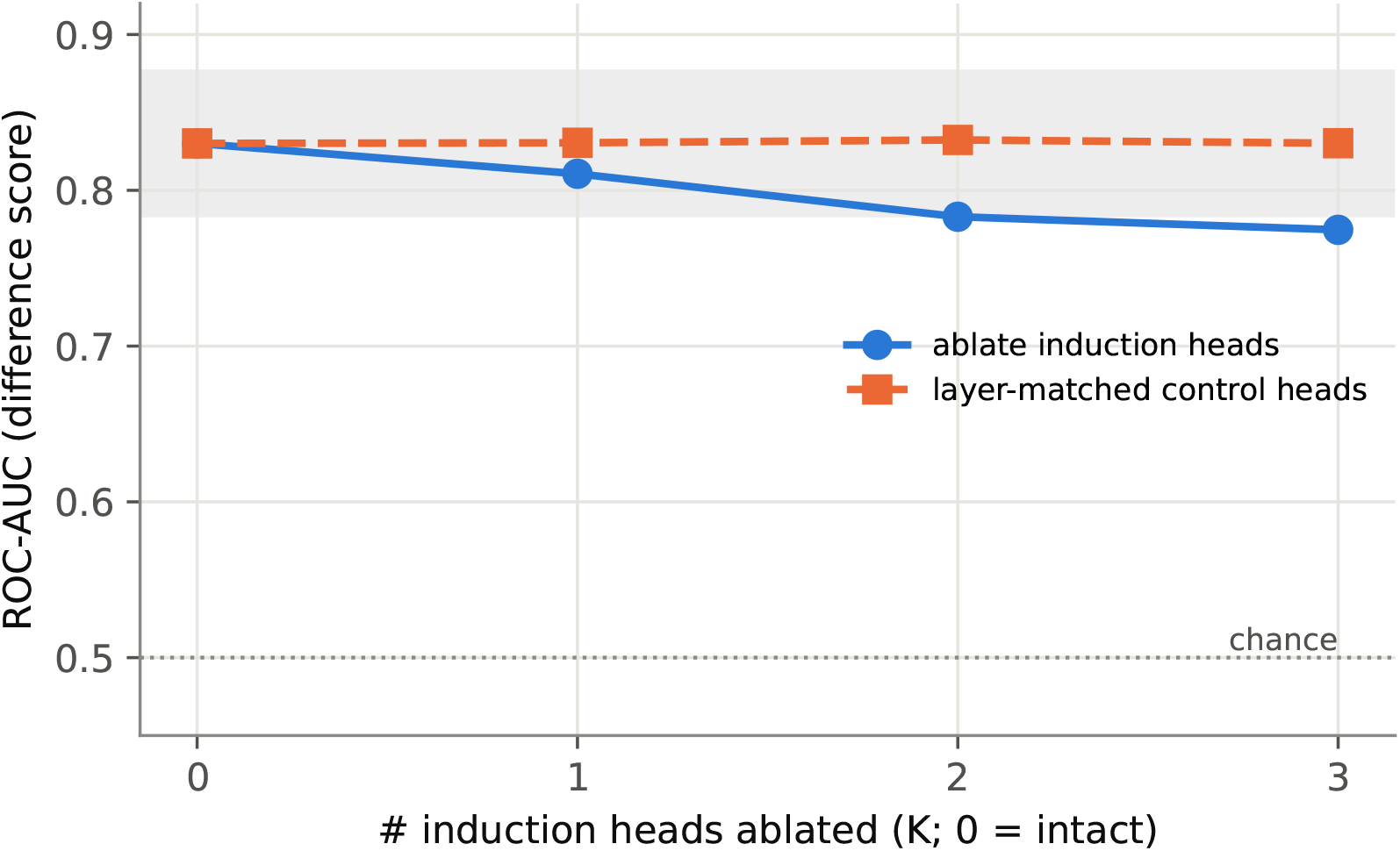
ProGen2-large binder screen after ablating *K* induction heads, using 16-example prompt. Grey area depicts noise band of the intact model.

**Figure 59:**
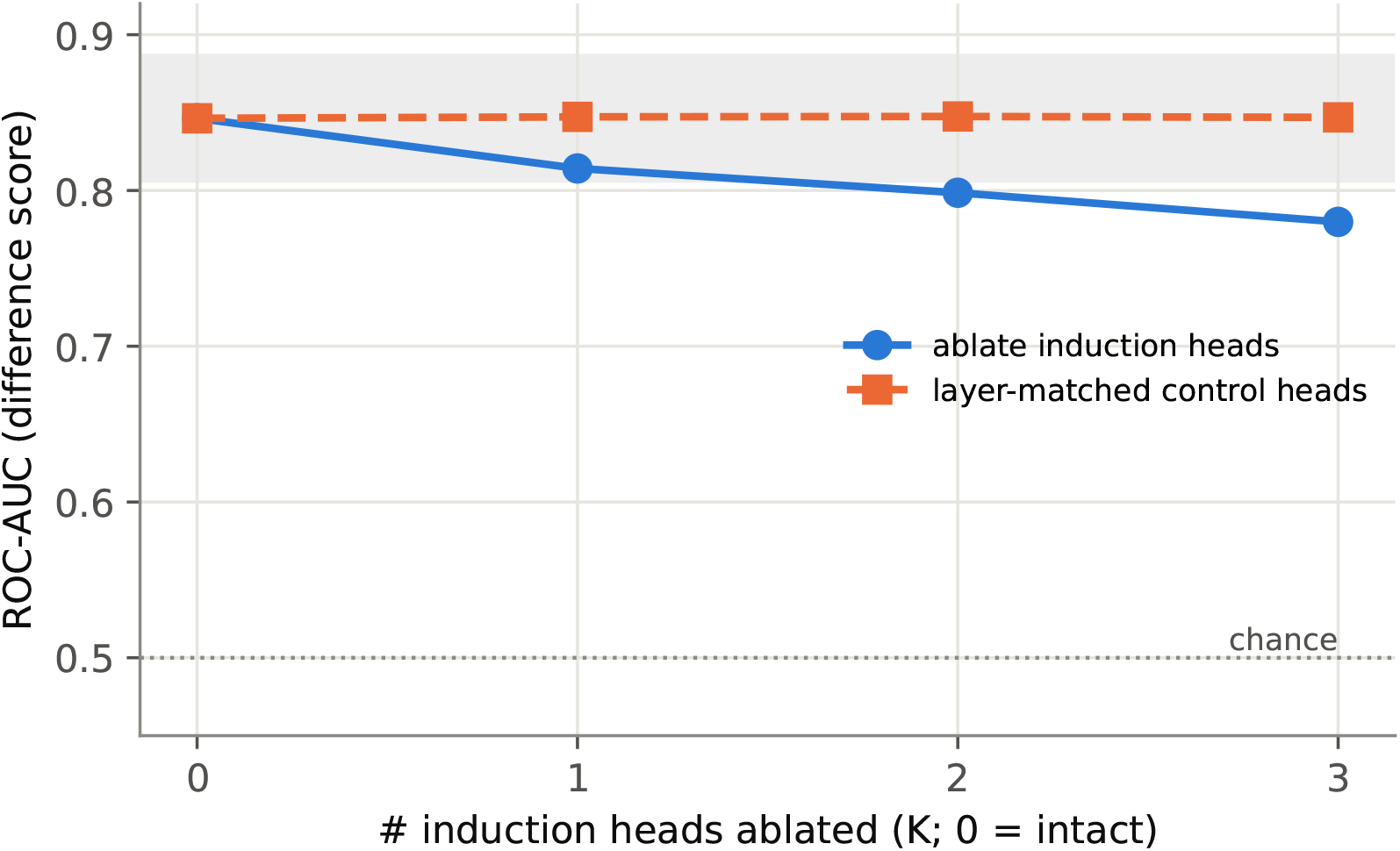
ProGen2-large binder screen after ablating *K* induction heads, using 32-example prompt. Grey area depicts noise band of the intact model.

### E.5 ProGen2-xlarge (6.44M)

**Figure 60:**
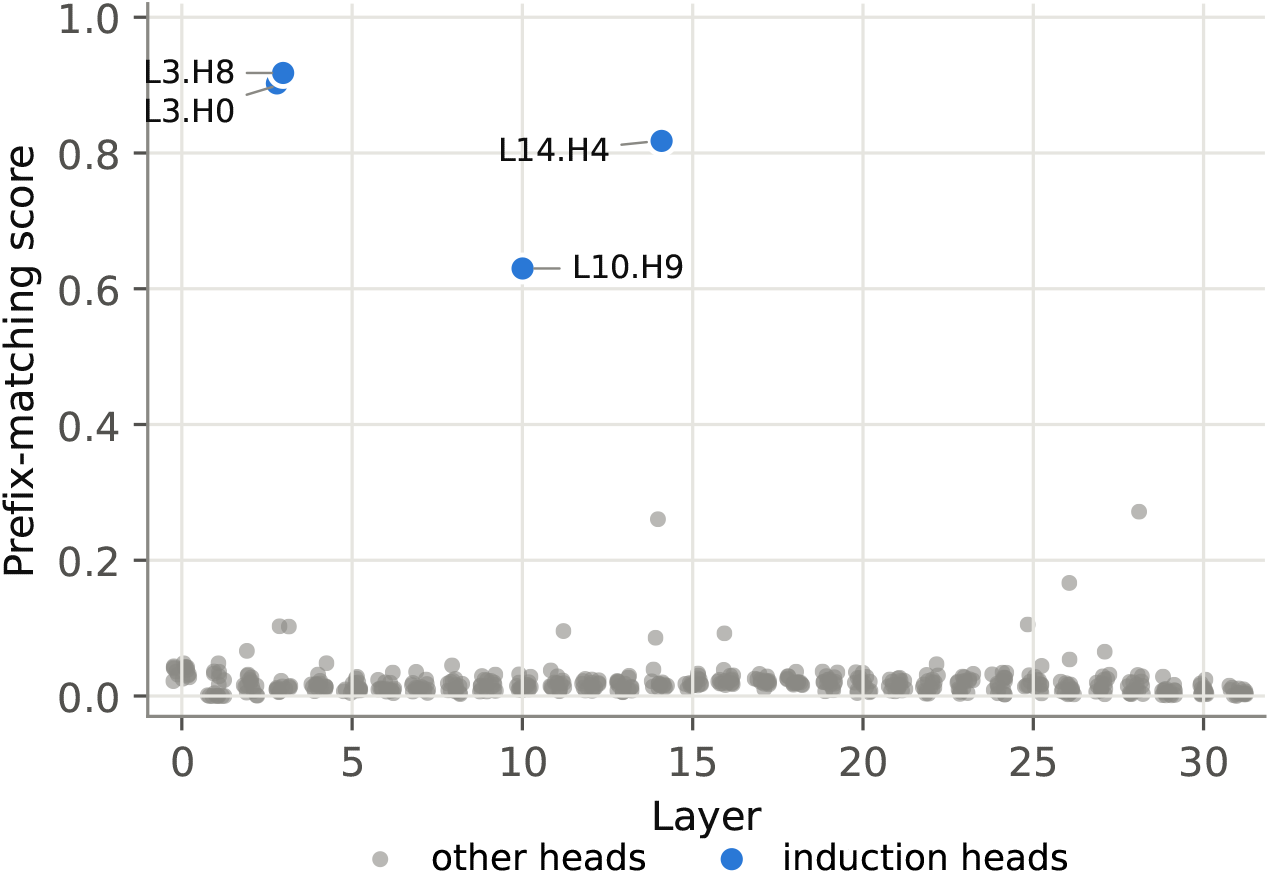
Prefix matching scatter in layers surrounding induction heads of ProGen2-xlarge.

**Figure 61:**
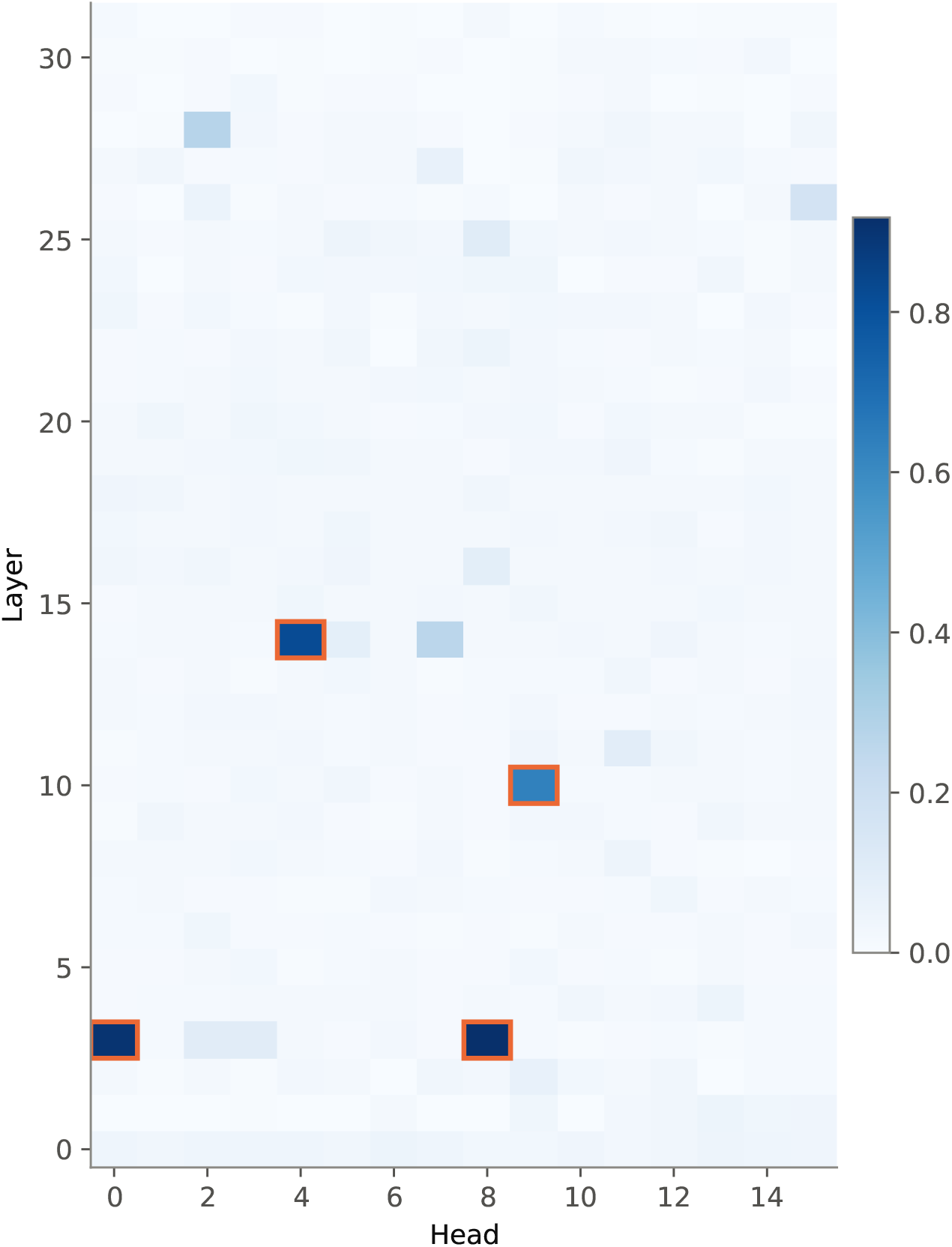
Prefix matching heatmap of induction-head and nearby layers in ProGen2-xlarge. Induction heads are highlighted with orange border.

**Figure 62:**
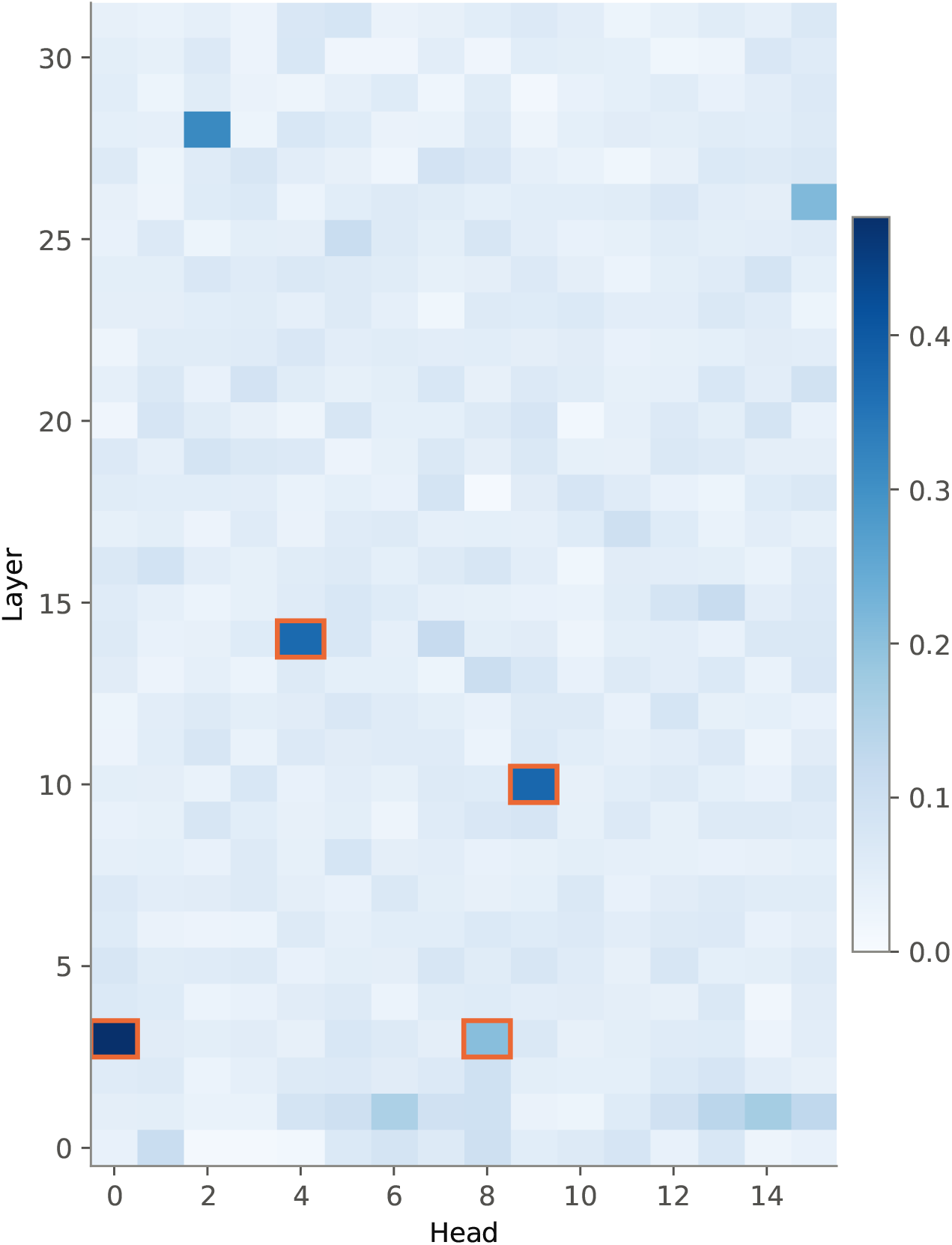
Copying heatmap of induction-head and nearby layers in ProGen2-xlarge.

**Figure 63:**
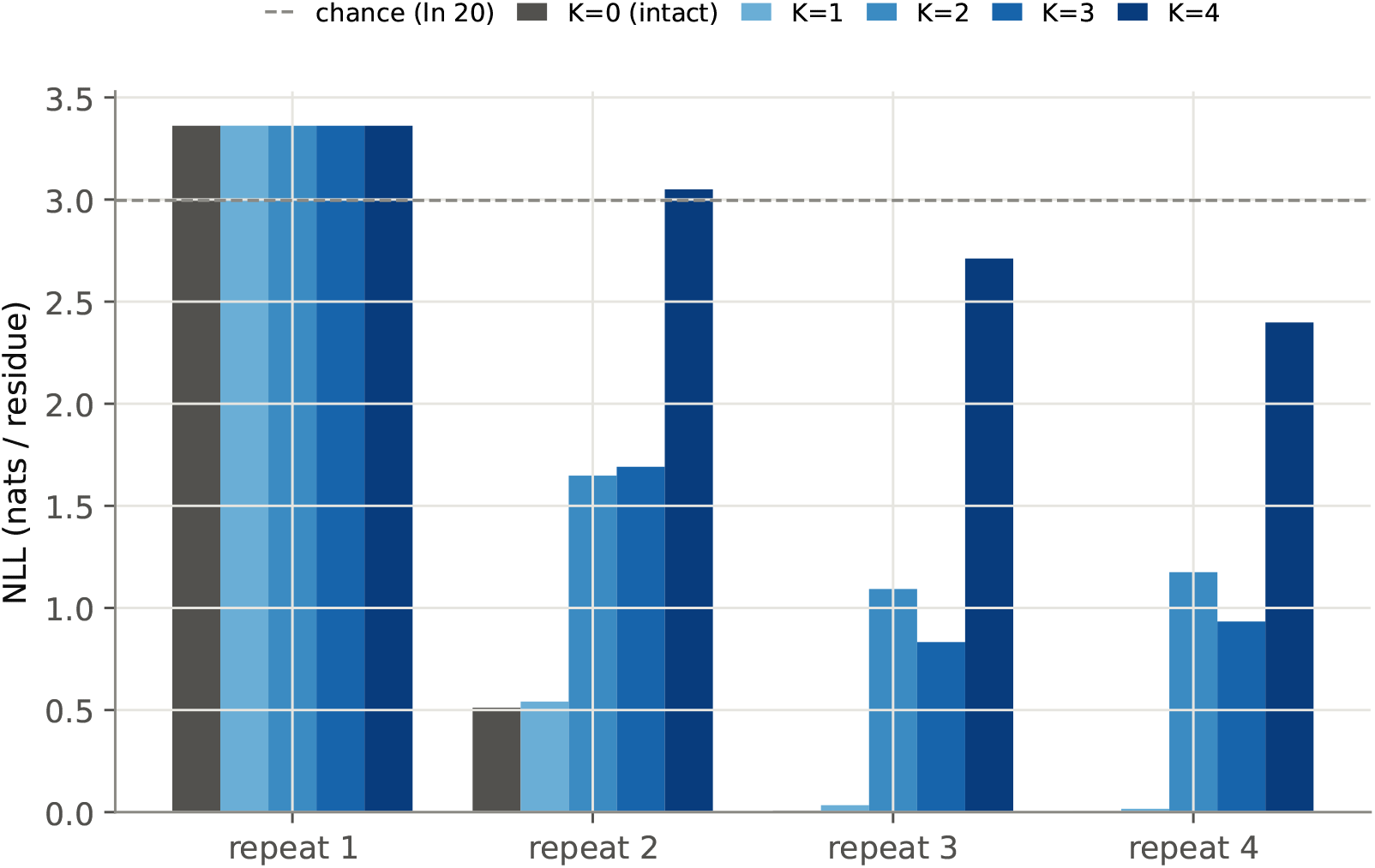
NLL loss-per-residue across all random sequence repeats during ProGen2-xlarge inference, with *K* induction head(s) zero-ed out.

**Figure 64:**
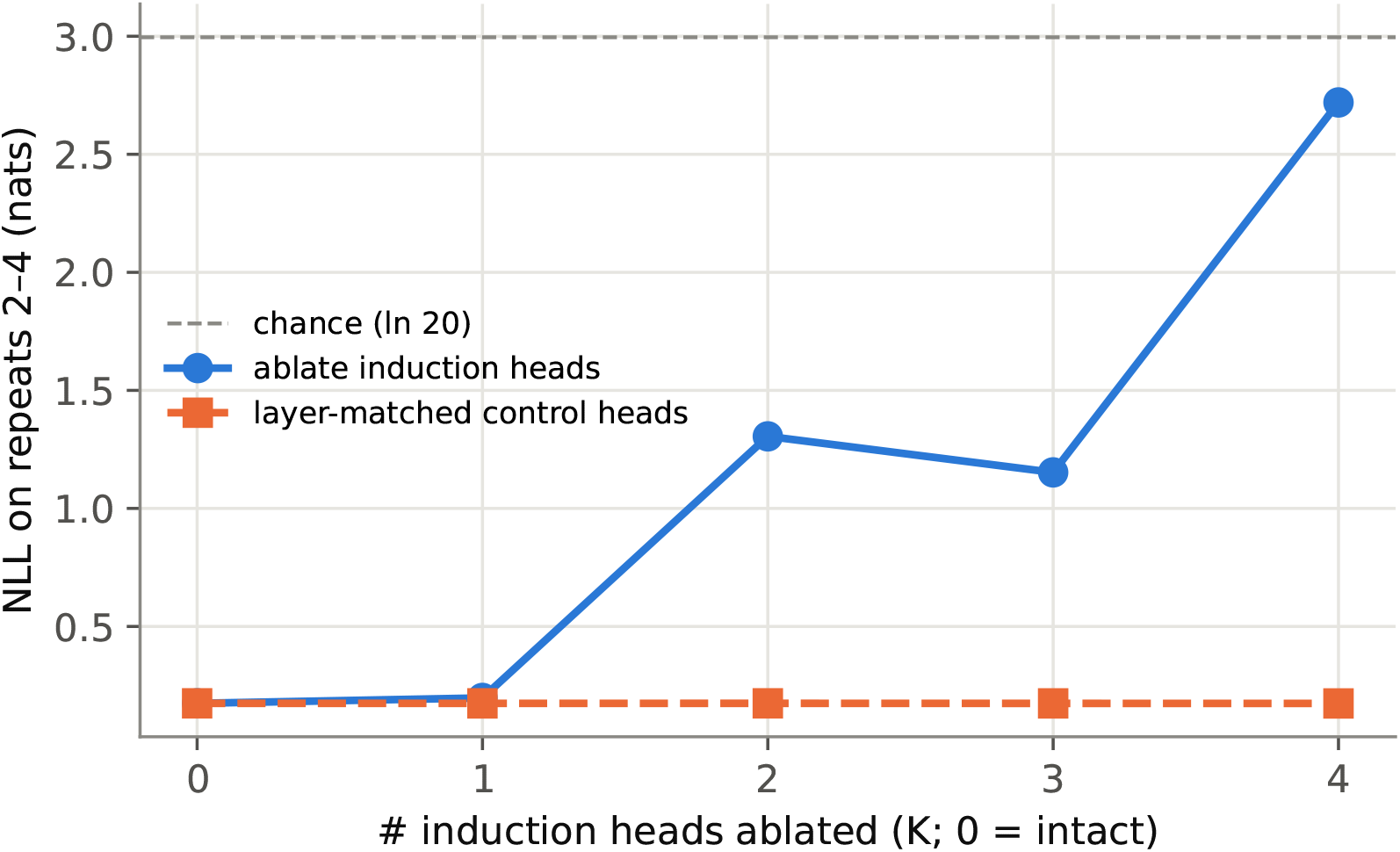
NLL loss-per-residue across random sequences repeats 2-4 between induction head ablation and control (ablating the same number of random, non-induction heads in the same induction layers) during ProGen2-xlarge inference.

**Figure 65:**
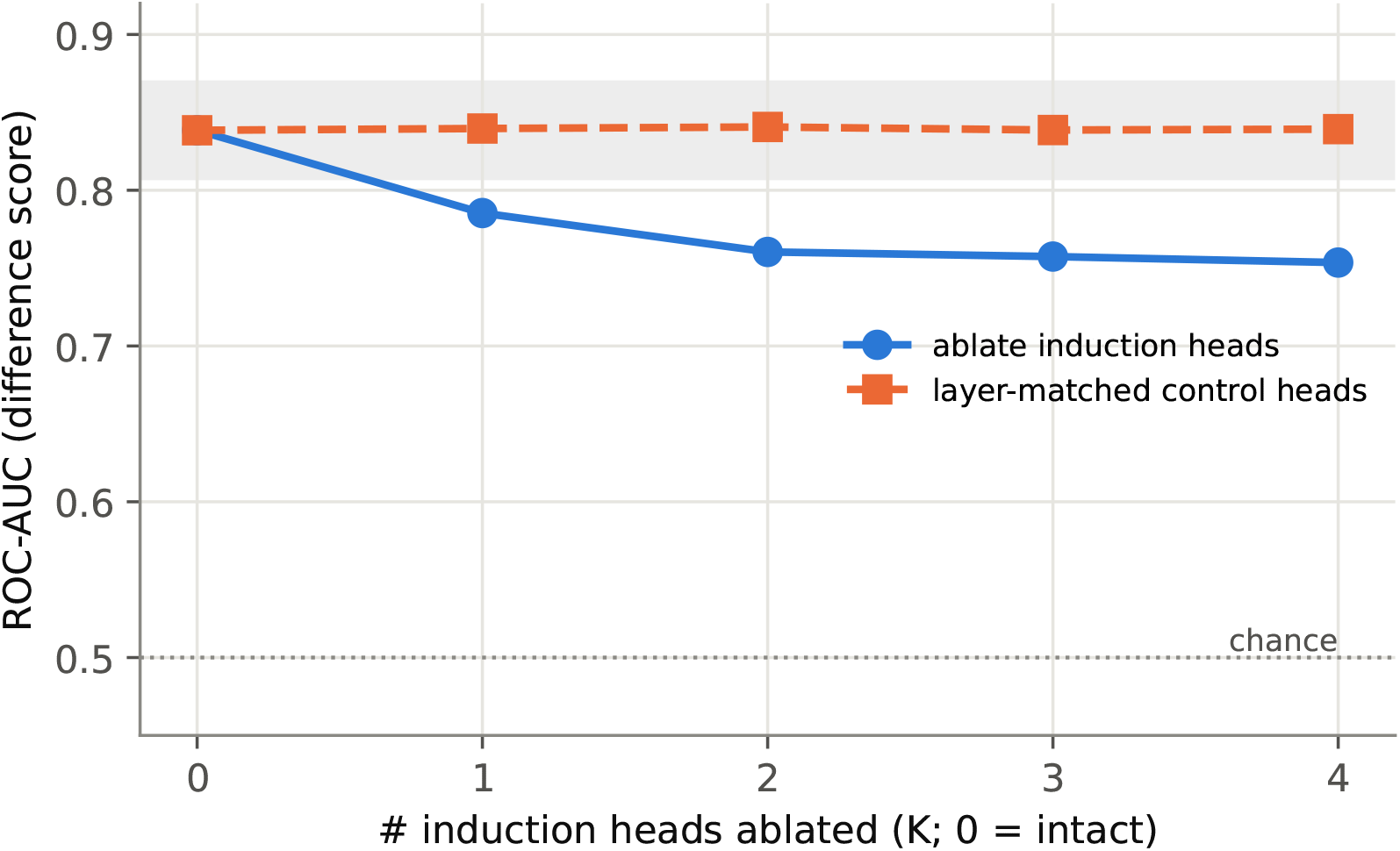
ProGen2-xlarge binder screen after ablating *K* induction heads, using 16-example prompt. Grey area depicts noise band of the intact model.

**Figure 66:**
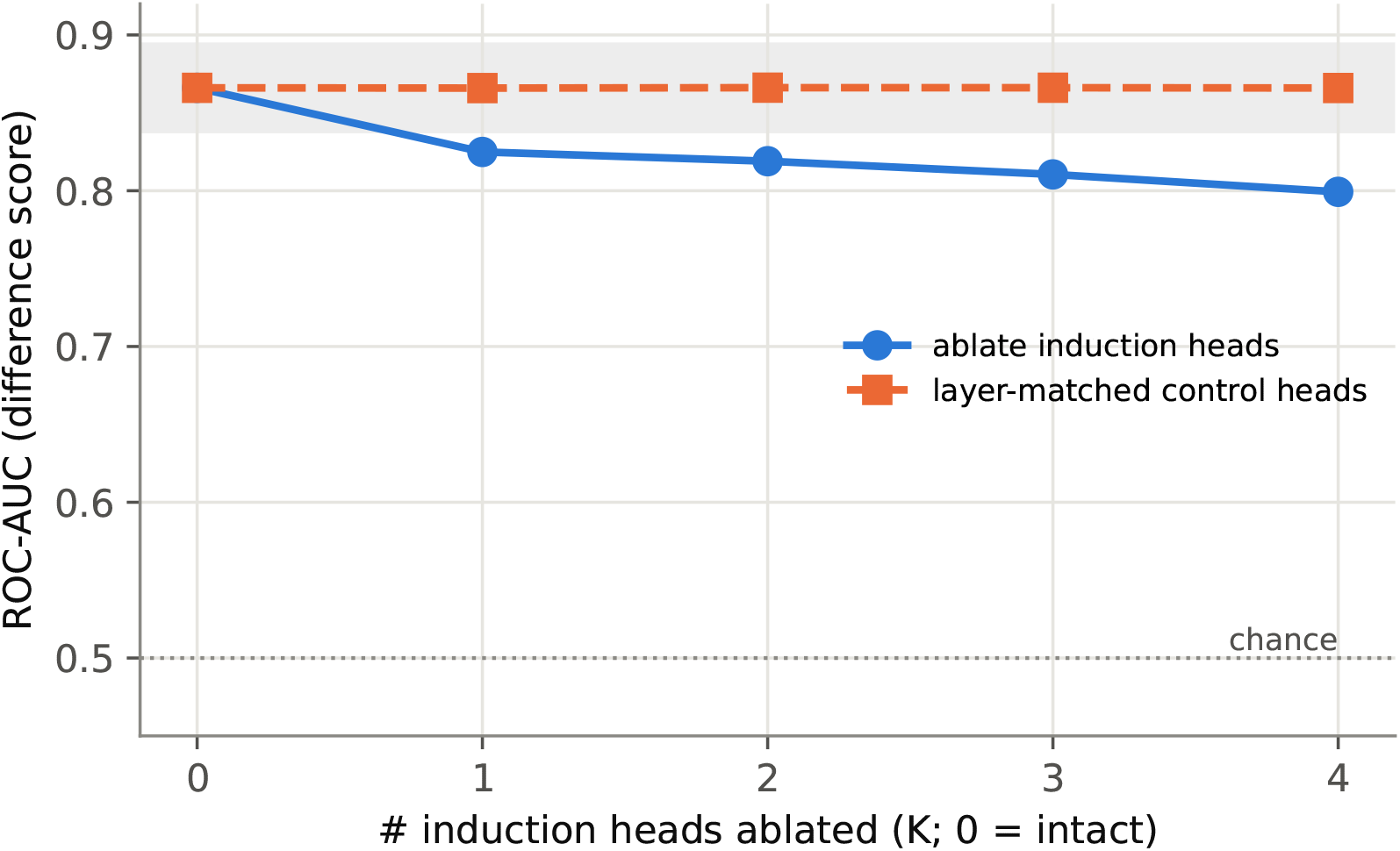
ProGen2-xlarge binder screen after ablating *K* induction heads, using 32-example prompt. Grey area depicts noise band of the intact model.

